# Impact of ligand binding on VEGFR1, VEGFR2, and NRP1 localization in human endothelial cells

**DOI:** 10.1101/2024.09.29.615728

**Authors:** Sarvenaz Sarabipour, Karina Kinghorn, Kaitlyn M Quigley, Anita Kovacs-Kasa, Brian H Annex, Victoria L Bautch, Feilim Mac Gabhann

## Abstract

The vascular endothelial growth factor receptors (VEGFRs) bind to cognate ligands to facilitate signaling pathways critical for angiogenesis, the growth of new capillaries from existing vasculature. Intracellular trafficking regulates the availability of receptors on the cell surface to bind ligands, which regulate activation, and the movement of activated receptors between the surface and intracellular pools, where they can initiate different signaling pathways. Using experimental data and computational modeling, we recently demonstrated and quantified the differential trafficking of three VEGF receptors, VEGFR1, VEGFR2, and coreceptor Neuropilin-1 (NRP1). Here, we expand that approach to quantify how the binding of different VEGF ligands alters the trafficking of these VEGF receptors and demonstrate the consequences of receptor localization and ligand binding on the localization and dynamics of signal initiation complexes. We include simulations of four different splice isoforms of VEGF-A and PLGF, each of which binds to different combinations of the VEGF receptors, and we use new experimental data for two of these ligands to parameterize and validate our model. We show that VEGFR2 trafficking is altered in response to ligand binding, but that trafficking of VEGFR1 is not; we also show that the altered trafficking can be explained by a single mechanistic process, increased internalization of the VEGFR2 receptor when bound to ligand; other processes are unaffected. We further show that even though the canonical view of receptor tyrosine kinases is of activation on the cell surface, most of the ligand-receptor complexes for both VEGFR1 and VEGFR2 are intracellular. We also explore the competition between the receptors for ligand binding, the so-called ‘decoy effect’, and show that while *in vitro* on the cell surface minimal such effect would be observed, inside the cell the effect can be substantial and may influence signaling. We term this location dependence the ‘reservoir effect’ as the size of the local ligand reservoir (large outside the cell, small inside the cell) plays an integral role in the receptor-receptor competition. These results expand our understanding of receptor-ligand trafficking dynamics and are critical for the design of therapeutic agents to regulate ligand availability to VEGFR1 and hence VEGF receptor signaling in angiogenesis.

## Introduction

Blood vessel formation is driven by sprouting angiogenesis of endothelial cells (Bautch, 2012; Jeltsch *et al*, 2013; Carmeliet, 2000). Angiogenesis plays a vital role in a host of physiological and pathological phenomena such as development, wound healing and tumor growth. Therapeutic induction of angiogenesis has long been a goal for ischemic diseases including peripheral artery disease and diabetes (Annex, 2013; Annex & Cooke, 2021). However, regulating angiogenesis in patients by manipulating its key cytokine drivers, including but not limited to members of the vascular endothelial growth factor (VEGF) family, has not yet succeeded, suggesting that the system and its regulation is not yet fully understood. Thus, a more comprehensive and mechanistic understanding of the molecular drivers of sprouting angiogenesis is required for the design of effective proangiogenic molecular therapies.

The VEGF family of ligands is encoded by five human genes including VEGF-A and placenta growth factor (PLGF), which produce bivalent ligands for the endothelial cell-surface VEGF receptors (VEGFRs). The protein ligands we study here – VEGF_121a_, VEGF_165a_, PLGF_1_, and PLGF_2_ – are prominent splice isoforms of these genes, and as constitutive antiparallel homodimers, each has two symmetric binding sites that enables the ligand to couple together two VEGFRs. The VEGFRs can pre-dimerize to some extent on endothelial cells in the absence of ligands (Sarabipour *et al*, 2016; da Rocha-Azevedo *et al*, 2020), although they are not fully active in signaling until bound and dimerized by their ligands (Sarabipour *et al*, 2016; Prahst *et al*, 2008). Upon binding of a ligand to two VEGFR1s or two VEGFR2s, the dimerized receptors trans-phosphorylate and initiate downstream signaling pathways (Simons *et al*, 2016; Lee *et al*, 2022; Dewerchin & Carmeliet, 2012; Markovic-Mueller *et al*, 2017). PLGF_2_ and VEGF_165a_ also contain a heparin binding domain (while PLGF_1_ and VEGF_121a_ do not), enabling these ligands to bind to heparan sulfate proteoglycans (HSPGs); this domain also harbors a binding site for the VEGFR co-receptor NRP1, and so these longer isoforms can bind NRP1, which facilitates binding of VEGF_165a_ to VEGFR2 and influences VEGFR2 trafficking (Prahst *et al*, 2008; Ballmer-Hofer *et al*, 2011).

VEGF and PLGF isoforms have tissue-specific expression patterns (Nowak *et al*, 2008; Rivron *et al*, 2012; Ruhrberg *et al*, 2002; Yonekura *et al*, 1999; Persico *et al*, 1999). Each isoform likely plays specific roles due to their different receptor binding and matrix-binding profiles. For example, while expression of VEGF_165a_ alone results in phenotypically normal vasculature features, VEGF_121a_ results in larger, more tortuous, less branched vessels (Ruhrberg *et al*, 2002; Vempati *et al*, 2014; Ng *et al*, 2001; Grunstein *et al*, 2000; Lee *et al*, 2005). PLGF is involved in activation of macrophages that can secrete angiogenic and lymphangiogenic factors (Selvaraj *et al*, 2003). PLGF also modulates tumor angiogenesis (Dewerchin & Carmeliet, 2012; Yang *et al*, 2013). However, PLGF is not required for normal murine development (Carmeliet *et al*, 2001), nor for exercise-induced angiogenesis (Gigante *et al*, 2004). Moreover, anti-PLGF antibodies do not block primary tumor angiogenesis (Bais *et al*, 2010) and PLGF knockdown also does not affect endothelial cell number, migration, or tube formation in cell culture (Xiang *et al*, 2014).

The membrane integral receptor VEGFR1 is considered a decoy receptor on endothelial cells during developmental angiogenesis (Nesmith *et al*, 2017; Kappas *et al*, 2008; Hiratsuka *et al*, 1998; Autiero *et al*, 2003). VEGF_165a_ and VEGF_121a_ bind to and initiate signaling complexes with both VEGFR1 and VEGFR2 dimers, while the PLGF ligands bind to VEGFR1 but not VEGFR2. As a result, it is hypothesized that VEGFR1 binding VEGF ligands decreases the binding of those ligands to VEGFR2 and thus modulates VEGFR2 signaling. We will characterize and quantify the VEGFR1 ‘decoy effect’ in this study. Likewise, PLGF ligand binding to VEGFR1 has been hypothesized to displace VEGF ligands from VEGFR1 and thus increase their binding to VEGFR2 and VEGFR2 signaling. A previous computational study suggested that this effect would be small (Mac Gabhann & Popel, 2004). Here we will quantify the potential for this effect in a more detailed model that includes receptor dimerization and much more detailed receptor trafficking. In particular, ligand-bound receptor dimers will be present both at the cell surface and on endosomes due to receptor trafficking (Bruns *et al*, 2010; Fearnley *et al*, 2014), and thus we may observe location-specific differences in the effects of VEGFR1 and PLGF on VEGFR2 activation.

The localization of membrane-integral receptors such as VEGFR1, VEGFR2, or NRP1 is dependent on trafficking within the cell. Simply put, ligand-receptor binding requires that the ligands and receptors be in the same place at the same time. Extracellular ligands can bind to receptors on the cell surface, while intracellular ligands in endosomes can bind to receptors in those locations. In a previous combined experimental-computational study, we quantified the trafficking of VEGFR1, VEGFR2, and NRP1 in the absence of exogenous ligands (Sarabipour *et al*, 2024); here, we examine how ligand binding can alter the trafficking of these receptors. In our model, we assume that ligated and activated receptor tyrosine kinases are phosphorylated on multiple sites intracellularly, and thus can interact with different cellular components that influence trafficking pathways or rates. Capturing these differences is important because the localization of these ligand-receptor signal initiation complexes likely lead to differential downstream signaling (Simons *et al*, 2016; Bruns *et al*, 2010).

Although the VEGF_165a_ and PLGF_1_ contributions to VEGFR2, VEGFR1 and NRP1 signaling have been studied *in vitro*, the combined effects of these ligands on VEGFR1 trafficking *in vitro* and *in silico* are not well understood. We aimed to understand the competition of VEGFR1 and VEGFR2 for VEGF binding, and the competition of VEGF and PLGF for binding to VEGFR1. The goals of this study were to: (1) elucidate the effect of VEGF_165a_ and PLGF_1_ on VEGFR1, VEGFR2, and NRP1 trafficking in human endothelial cells; (2) predict the distribution of VEGFR1, VEGFR2, and NRP1 on the cell surface and internal compartments as a function of VEGF_165a_ and PLGF_1_ treatment time and dosing; (3) investigate how this modified receptor trafficking impacts the distribution of ligated, signaling receptors; (4) compare the predicted effects of VEGF_165a_ vs VEGF_121a_ and PLGF_1_ vs PLGF_2_ to parse the effect of NRP1 binding on VEGFR1 and VEGFR2 trafficking; (5) quantify VEGFR1’s decoy effect on VEGFR2; and (6) quantify PLGF’s contribution to the decoy effect. Computational models provide key insights into the combinatorial effects of multiple receptors and ligands in regulation of signaling pathways *in vitro* and *in vivo* (Mac Gabhann *et al*, 2010; Wu *et al*, 2010b; Sarabipour *et al*, 2024). Here, we present such measurements and modeling with specific parameters for ligand-bound receptor trafficking that are quantified here for the first time in any cell type. To our knowledge, this is the first set of VEGFR1, VEGFR2, and NRP1 trafficking experiments on the same human cell type treated with VEGF_165a_ or PLGF_1_, and the first computational model to include detailed trafficking of VEGFR1, VEGFR2, and NRP1 explicitly under ligand treatment.

## Materials and Methods

### Computational Model Construction

We previously developed and parameterized a molecularly-detailed mechanistic computational model of the dynamics of membrane-integral VEGF receptors, VEGFR1, VEGFR2, and NRP1, specifically their synthesis, trafficking, and degradation in human umbilical vein endothelial cells (HUVECs) (Sarabipour *et al*, 2024). Here, we build on and extend this model to include ligand-receptor binding for four key ligands (VEGF_165a_, VEGF_121a_, PLGF_1_, and PLGF_2_), and to account for the ability of this ligand-receptor binding to alter receptor trafficking. The model includes all the reactions and parameters identified for HUVECs in our previous study, and here we estimate values for newly-added ligand-involved processes using experimental data from human endothelial cell culture.

The model uses coupled, nonlinear, deterministic ordinary differential equations describing the key biochemical and biophysical reactions including receptor-receptor dimerization (coupling), ligand-receptor binding, and trafficking of receptors and ligand-receptor complexes (Figure 1). The ligand-receptor binding and receptor-receptor coupling reactions can occur in every model compartment where the associated species are present. We use experimentally-derived surface receptor expression levels and interaction kinetics for receptors expressed in HUVECs that have been previously measured or derived, and in running the simulations we incorporate the specific geometry and initial concentrations of the *in vitro* HUVEC culture experiments being simulated.

**Figure 1.**
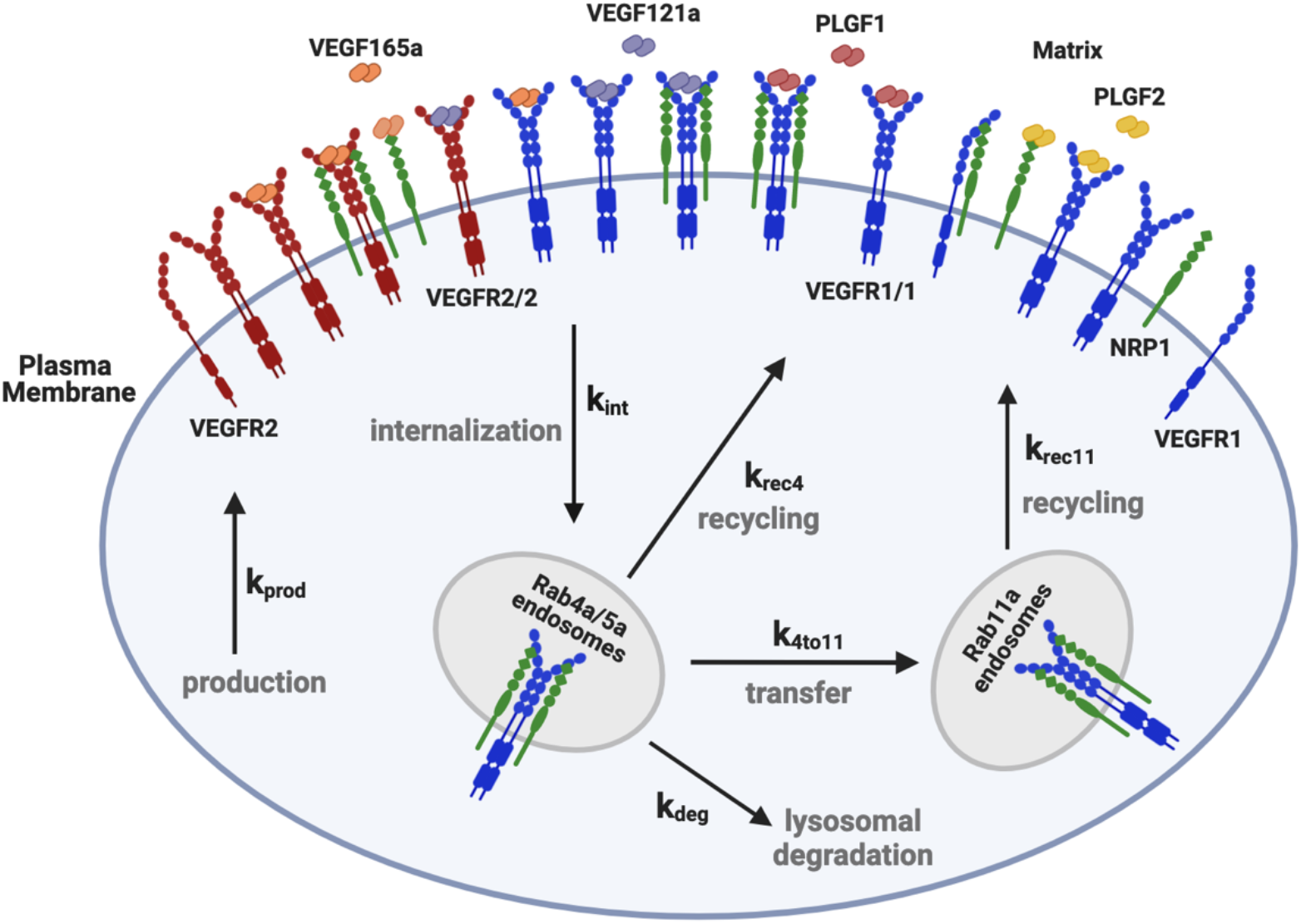
Diagram of molecular interactions of the computational model. Cellular biophysical and biochemical reactions between VEGFR1, VEGFR2 and NRP1 receptors, extracellular matrix components and VEGF_121a_, VEGF_165a_, PLGF_1_ and PLGF_2_ ligands. The receptors can homodimerize (at a lower level than that induced by ligands), and unlike VEGFR2, VEGFR1 can form a complex with NRP1 in the absence of ligands (Fuh *et al*, 2000). PLGF_1_ binds to VEGFR1 in the absence or presence of NRP1 binding to VEGFR1. VEGF_165a_ binds to VEGFR1, VEGFR2 and NRP1 and the matrix proteins. VEGF_165a_ can only bind to NRP1 or matrix, not both simultaneously. PLGF_2_ and VEGF_165a_ can only bind to VEGFR1 when VEGFR1 is not bound to NRP1. VEGF_165a_ can bind to NRP1 and VEGFR2 simultaneously. During trafficking, surface protein complexes (monomeric, dimeric, or higher order) can be internalized (rate constants denoted k_int_). Early endosomal (“Rab4a/5a”) receptors can be degraded (rate constant k_deg_), recycled (rate constant k_rec4_), or transferred to the Rab11a compartment (rate constant k_4to11_) which leads to an additional recycling pathway (rate constant k_rec11_). New surface receptors (monomers) are produced at rate k_prod_. Each of the rate constant values can be different for the different receptors. Reaction rates and species concentrations are detailed in Supplementary Tables S1-S18.

#### Ligand-receptor interactions

All four of the ligands studied here bind to VEGFR1; the VEGFA isoforms can also bind to VEGFR2 while the PLGF isoforms do not. Binding to the co-receptor NRP1 requires a NRP1-binding domain that shorter isoforms typically lack; of the ligands in our model, only VEGFA_165a_ and PLGF_2_ bind to NRP1. VEGFA_165a_ can thus bridge VEGFR2 and NRP1 by binding to each via non-overlapping binding sites. However, this NRP1 binding is prevented when NRP1 is associated with VEGFR1 (Fuh *et al*, 2000), perhaps due to steric hindrance, so only the shorter isoforms (VEGFA_121a_ and PLGF_1_) bind to VEGFR1-NRP1 complexes, and only via VEGFR1 binding. These ligand-receptor interactions are summarized in Supplementary Table S1, and along with the receptor-receptor interactions and perhaps most crucially the receptor-to-liganded-receptor interactions (i.e., dimerization), form the basis for the many molecular complexes that can be formed (Supplementary Tables S2-S11).

#### Compartments

The computational model has three subcellular compartments representing different locations within the cell: the cell surface (specifically, the outward-facing portion through which receptors interact with extracellular ligands) which includes the fluid space outside the cell (i.e., culture media); early endosomes (Rab4a/Rab5a-expressing); and recycling endosomes (Rab11a-expressing). The receptors and receptor complexes move between these compartments via trafficking pathways at different rates, though we do make some simplifying assumptions to avoid an excessive number of independent parameters. The model also includes a degradation compartment, representing lysosomes or other degradative pathways, and crucially this is not included in counts of receptor numbers for the purpose of comparison to experimental results. Instead, the purpose of this compartment is to keep track of the cumulative amount of degraded receptors; the process of protein degradation in the model is represented by receptors or ligand-receptor complexes moving to the degradation compartment. We assume that the individual surface and endosomal compartments are well mixed, i.e., they are of uniform (but not constant) concentration, and we assume that receptor levels are sufficiently high to justify the continuum approximation, as previously shown (Mac Gabhann *et al*, 2005), and thus can be represented with deterministic ordinary differential equations.

#### Protein synthesis and degradation

In the model, there is continual constant synthesis of new receptors to balance receptor degradation, thus representing the continuous turnover of proteins in the cell. Once synthesized, receptors are inserted into the cell surface compartment. In these simulations, ligands are added exogenously to the culture media as a step change in concentration, following a period of pre-simulation, in which the dynamics of receptor production, trafficking, and degradation proceed to steady state, to set up receptor levels on the cell surface and in the intracellular locations. Production and secretion of ligand by the cell could be incorporated, but the levels of exogenous ligand (50 ng.ml^-1^ or similar concentrations, Supplementary Table S12) are sufficiently high to likely overwhelm any endogenous ligand. Receptor production rate values are identified using optimization to match the simulated cell surface receptor levels to published measurements, and the rates are given in Supplementary Table S13. The degradation of receptors and internalized ligands occurs as a first-order process from the intracellular pool.

#### Receptor trafficking processes

The trafficking processes, moving receptors and ligand-receptor complexes between subcellular compartments, are summarized in Figure 1, and these processes are modeled as first-order transport rates. Internalization moves surface receptors to the early endosome compartment; from these endosomes, receptors can be degraded, recycled to the surface, or trafficked to a second endosome compartment for recycling. Two recycling pathways – one via Rab4a and one via Rab11a – are included based on evidence that VEGFR2 and NRP1 use both pathways in endothelial cells (Ballmer-Hofer *et al*, 2011). Appropriate receptor-specific and HUVEC-specific values of the production, trafficking, and degradation rate constants for the unliganded receptors were identified in our recent study (Sarabipour *et al*, 2024) and can be found in the Supplementary Table S14.

For ligand-bound receptors, the rates of trafficking may be different – as part of this study we explored the effect of multiple different ligands on VEGF receptor localization, leading to identification of a parsimonious mechanism to explain the observations (see Results). For trafficking processes that we identified to be ligand-dependent, the rate constants are different for those receptors when ligands are bound to them; if not ligand-dependent, then we assumed that the rate constants for liganded and unliganded receptors were the same. This greatly reduced the number of independent parameters needed to describe trafficking of the many receptor complexes in the system.

#### Receptor-receptor coupling and the impact of compartment geometry

In the absence of ligands, VEGFR1 and VEGFR2 are present as a mix of monomers and dimers due to reversible homodimerization in the absence of ligands (Sarabipour *et al*, 2024, 2016; da Rocha-Azevedo *et al*, 2020); this is in addition to the ligand-induced dimerization that will be described below. VEGFR1 and NRP1 can also bind (couple) directly to each other, while VEGFR2 and NRP1 do not; VEGFR2 and NRP1 appear to only interact when bridged by a ligand that binds both. The computational model describes the receptor levels at each location in units of #/cell (i.e., the number of molecules per cell), which makes incorporating first-order trafficking parameters simpler, and makes it straightforward to match calculated levels to experimental measurements; however, this approach impacts on the rate constants describing the receptor coupling reactions. This is described in detail in our previous work (Sarabipour *et al*, 2024); briefly, units of #/cell does not take into account the density (and therefore likelihood of interaction) of the receptors in compartments of different sizes. Assuming that the rate constants are the same on a per-surface-area basis (for two surface-associated molecules), then we can use the compartment surface areas (Supplementary Table S12) to calculate appropriate local rate constants:

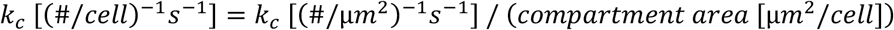

This reduces the number of unique parameters in the model (making the model simulations more tractable). The resulting local parameter values for receptor-receptor coupling are given in Supplementary Table S15.

#### Ligand concentrations, ligand-receptor binding, and the impact of compartment geometry

As noted above, the units used in the model are #/cell, and therefore the concentration of the ligand added to the extracellular media (typically expressed in ng.ml^-1^) is converted to #/cell using the following conversion factor using molecular weight (g/mol), the volume of fluid in the well (ml/well), and the number of cells per well (cell/well):

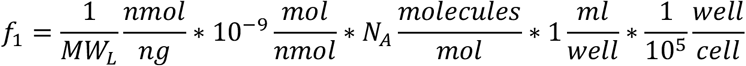

Where MW_L_ = molecular weight of ligand (VEGFA_165a_, 44 kDa; VEGFA_121a_, 28 kDa; PLGF_1_, 29.7 kDa; PLGF_2_, 34.6 kDa), N_A_ = Avogadro’s number (6.022 x 10^23^ molecules/mole), and thus, the values of *f_1_* for the four ligands are, respectively, 1.37 x 10^5^, 2.15 x 10^5^, 2.03 x 10^5^, and 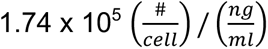.

Further, the ligand-receptor binding rate constants must be adjusted from the standard reported form based on experimental measurements (Supplementary Table S16) to account for three things: the full dimerization model used here compared to the more common 1:1 binding model assumed to generate the parameters; to match the #/cell units of the model; and because binding can occur in each subcellular location, the volume associated with that location affects the local rate constant. The resulting parameter values are given in Supplementary Table S17. The binding rate constants are effectively one-quarter of those measured, to account for the bivalency of both ligands and receptor dimers. The unbinding rate constants are the square root of half the measured rate, to account for the two unbinding events that must occur for the ligand to fully dissociate from a dimer (for more on the derivation of these relationships, see our previous work on dimerization (Mac Gabhann & Popel, 2007)).

The binding rate constants are then adjusted using local volumes to “#/cell” units as described above for ligand concentrations, while unbinding rate constants need no adjustment due to being first order processes. As previously described (Sarabipour *et al*, 2024), the endosomes enclose approximately 15 fL (about 1.5% of HUVEC volume), and assuming a 3:1 Rab4a:Rab11a split for both volume and cell surface area, this results in 11.25 fL/cell and 3.75 fL/cell for the two intracellular compartments, respectively, as compared to the 10^7^ fL/cell of extracellular volume (based on 1 mL media and 10^5^ cells per well; see Supplementary Table S12). As a result, the apparent (volume-adjusted) rate constants for ligands binding to receptors are much higher in endosomes than extracellularly (Supplementary Table S15), which accounts for the “#/cell” levels of ligand being concentrated in a smaller volume. It is important to have these units aligned, because the difference in size between the extracellular reservoir and the intracellular reservoir has a significant effect on ligand-receptor binding and competition.

#### Ligand-induced receptor coupling

Following ligand binding either to a receptor monomer or to a receptor dimer, the bivalent ligand can now bind to a second monomer or to the second receptor in the dimer. These events are related but somewhat different; binding to a second monomer is a second-order reaction involving molecules diffusing in the plane of the membrane (rate constant *k_c,LR_*), while binding to the second receptor in a dimer is effectively an intramolecular rearrangement, and thus a first order reaction (rate constant *k_Δ,LR_*). We use the Δ symbol to denote that the molecule formed has three bonds – two ligand-receptor bonds and one receptor-receptor bond between the two ligand-bound receptors. The values for these rate constants are obtained as described previously (Mac Gabhann & Popel, 2007); briefly,

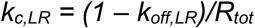

Where *R_tot_* is the receptor level on the surface (typically where experimental binding assays are sampling). The resulting values are then modified by surface area for the endosomal compartments (Supplementary Table S18). Comparing the receptor-receptor coupling and ligand-receptor coupling rate constants, we see that the ratio (*k_c,RR_/k_c,LR_*) is small (Supplementary Table S18), suggesting that ligand coupling of receptors is stronger than receptor-receptor coupling, which makes sense for ligand-induced dimerization.

Using the previously demonstrated (Mac Gabhann & Popel, 2007) relationships that

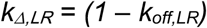

and

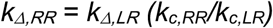

We obtain values for these rate constants which represent, respectively, the coupling of a ligand to a second receptor in the same complex, and the coupling of two receptors that are already both bound to the same ligand (Supplementary Table S18). The similarity of the dimerization model parameterized in this way to the typical 1:1 ligand-receptor binding model can be seen in the virtual Scatchard plots generated using the models (Supplementary Figure S1).

#### Pre-simulation and simulating ligand additions and perturbations

The balance of synthesis, degradation, and trafficking of receptors results in a steady-state, constant surface level of VEGFR1, VEGFR2, and NRP1 populations in the absence of ligand. These rates and steady state levels of receptors have previously been optimized and analyzed (Sarabipour *et al*, 2024). Here, we study what happens when that steady state is perturbed by ligand addition. We assume that addition of ligands is a sudden step-change in the extracellular concentration of ligands, and then use the model to simulate several hours of experimental time. Other perturbations can be included either as sudden changes to parameter values at the time of administration (e.g., siRNA knockdown of Rab4a and Rab11a to decrease in the levels of those proteins), or as changes incorporated into the pre-simulation because they are administered long before the ligand and thus the new state of the cells must be simulated, for example receptor knockdowns.

#### Model solution, model outputs, and comparison with experimental data

The complete model contains 281 molecular complexes (Supplementary Table S2, and Supplementary Tables S3-S11), including separately tracking the levels of each ligand, receptor, receptor-receptor complex, and ligand-receptor complex at each subcellular location (plasma membrane, Rab4a/5a endosomes, Rab11a endosomes) plus the degraded molecules. Code to simulate this set of 281 coupled ordinary differential equations that comprise this model was generated using the rule-based BioNetGen software; we then used the BioNetGen Visual Studio extension (Harris *et al*, 2016) to turn our BioNetGen model code into a MATLAB-compatible code in m-file format. This m-file encodes the molecules, reactions, and equations, and we then modified the m-file’s header to function under the control of our bespoke drivers in MATLAB, for example allowing us to run simulations with different parameter values, optimization, and sensitivity analysis. The code is provided in an online repository (Sarabipour & Mac Gabhann, 2024).

The output of the model is the concentration of each molecule or molecular complex at each location over time. To facilitate direct comparison to experimental data, the output concentrations of specific molecules or complexes are combined into aggregated quantities of interest. For example, the total VEGFR1 levels on the cell surface is the sum of all VEGFR1 in that compartment, whether in monomer form or complexed with other receptors or involved in ligand binding. If a complex contains two VEGFR1, it contributes double to the total level compared to a complex with one VEGFR1. “Internal” receptors are the sum of receptors in Rab4a and Rab11a locations; and “Total” or “Whole cell” receptors are the sum of receptors in Surface, Rab4a, and Rab11a locations. Degraded receptors are not included in any of the aggregations. Depending on the experimental measurement, e.g., levels of proteins from whole cell lysates, or only surface proteins isolated via biotin labeling, we use the appropriate aggregation of simulation results to compare to the experimental results. For predictions of ‘active’ molecules, i.e., receptor complexes expected to be capable of signaling, we used the sums of complexes containing a ligand that is bound to two VEGFR1 molecules or two VEGFR2 molecules.

We normalized the experimental data in the form of western blot band intensity for proteins of interest to actin band intensity control for equal whole cell protein loading (all proteins in the cell). We further normalized the band intensity corresponding to ligand treatment (VEGF_165a_ or PLGF_1_) conditions to the no ligand condition for each protein and experiment. For biotin-labeling experiments, the normalization is instead to PECAM (for surface receptors) or tubulin (for internal receptors) band intensity values. Since most of the experimental data is in relative units, not absolute units (exception is baseline surface receptor measurements, Supplementary Table S4), in order to compare experimental results to simulation results, we normalized the experimental data to experimental control values (no treatment conditions, or time zero condition depending on the experiment), and similarly normalized the simulation outputs described above.

#### Model parameter optimization

For HUVECs, we previously estimated the values of 15 receptor trafficking parameters and 3 receptor production rate parameters in the absence of exogenous ligands (Sarabipour *et al*, 2024). Rather than performing a complete additional optimization to estimate values for each of the equivalent 15 ligated receptor trafficking parameters, we expected that some of the trafficking parameters describing the movement of ligated receptors would be similar to their non-ligated counterparts. Thus, we performed simulations varying the values of trafficking rates one-by-one to identify parameters that most strongly affected model outputs, and then compared the results to experimental data for receptor localization following ligand addition. In this way we identified the most parsimonious model of how ligands impact trafficking parameters. As a result of this approach, there is sufficient data for the parameters being optimized to be identifiable (six data points to identify one parameter).

#### Sensitivity analysis

We performed univariate sensitivity analysis to identify parameters that most strongly affect model outputs. Key parameter values were varied as described in the text and the change in each selected output calculated. Model outputs selected included levels of expression of the receptor at the surface, inside the cell, and across the whole cell, over time (Supplementary Figures S2-S4) in response to ligand addition. We focused on the outcomes at 60 and 240 minutes (Supplementary Figures S5-S6) for comparison to experimental data.

#### Decoy effect

VEGFR1 has long been suggested to act as a decoy receptor; in other words, by binding ligand it prevents that ligand from then binding to the canonical signaling receptor VEGFR2. Functionally, this would require VEGFR1 to deplete the local ligand levels such that less ligand is available to bind to VEGFR2, since they compete for a common local pool of ligand. There is also considerable evidence that VEGFR1 is a signaling receptor in its own right, at least in adult animals (Bautch, 2012; Dewerchin & Carmeliet, 2012; Incio *et al*, 2016; Ganta *et al*, 2018). To better understand the balance between VEGFR1’s decoy role and its signaling role, it is useful to know the magnitude of the effect of VEGFR1 expression on the availability of VEGF for VEGFR2 binding. To quantify this, we created a *decoy effect* metric, expressed as the relative change in the formation of active VEGFR2 complexes due to the presence of VEGFR1. To calculate it, we simulate a scenario where VEGFR1 is absent (R1–) and compare this to a scenario with it present (R1+):

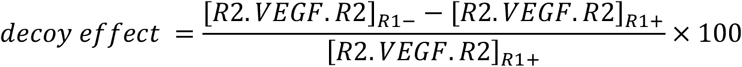

Expressed as a percentage, the size of this metric quantifies the degree to which the activation of VEGFR2 increases when VEGFR1 is removed; and thus reflects the amount that the presence of VEGFR1 modulates VEGFR2 activation downwards (the decoy effect). A positive value for the decoy effect metric suggests that VEGFR1 does have an impact via competition on VEGFR2 activation by VEGF, separate from any downstream signaling crosstalk effects. As with other metrics of binding and activation, the size of this decoy effect will depend on the expression and co-localization of the receptors (and of co-receptors), which are themselves dependent on the trafficking parameters; receptors cannot have an effect if they are not locally available. The above metric calculates VEGFR1’s effect on VEGFR2; we calculate similar metrics to quantify the effect of VEGFR2 deletion on VEGFR1 activation, and the effect of NRP1 deletion on VEGFR1 and VEGFR2 activation.

### Experimental Methods

#### Cell culture

Human umbilical vein endothelial cells (HUVECs) were cultured at 37 °C in EBM-2 medium supplemented with a bullet kit (EGM-2) and used at passages 3–5. For each experiment, cells were used at confluence (2-3 days post-plating) (see Supplementary Table S19 for manufacturers, catalog numbers, and other details of reagents).

#### Perturbations

Several treatments were used to inhibit specific processes in the receptor trafficking system (Figure 1). To deplete expression of recycling-associated proteins Rab4a and Rab11a in HUVECs, cells were grown to 70-80% confluency in 10 cm^2^ plates and incubated with 200 pmoles of siRNA (Silencer Select Locked Nucleic Acids) and 20 μL Lipofectamine 3000 (ThermoFisher, #L3000015). Cells were transfected with control (non-targeting) siRNA duplex and siRNA duplex targeting Rab4a and Rab11a. After 48 hours, the cells were serum-starved (OptiMem + 0.1% FBS) for 18 hours, followed by ligand addition.

#### Whole cell protein isolation

For selected experiments, total protein expression of VEGFR1, VEGFR2, and NRP1 from whole cell lysates was measured. At appropriate time points, RIPA buffer (50 mM Tris HCl, 150 mM NaCl, 1.0% (v/v) NP-40, 0.5% (w/v) Sodium Deoxycholate, 1.0 mM EDTA, 0.1% (w/v) SDS and 0.01% (w/v) Sodium Azide) and protease/phosphatase inhibitor (Cell Signaling Technology) at a pH of 7.4 was added to the plates, lysates were collected with cell scrapers, subsequently boiled for 5 minutes at 95°C, and used for immunoblot analysis as described below. The experiments were repeated three times.

#### Internal and cell surface protein isolation

For other experiments, the expression of surface receptors and intracellular receptors was separated and measured via biotinylation. Biotinylation and isolation of cell surface proteins was performed according to the manufacturer’s protocol (Pierce™ Cell Surface Biotinylation and Protein Isolation Kit, cat#A44390). Briefly, serum-starved HUVECs were washed twice in 6 ml of ice-cold PBS (Sigma-Aldrich) to stop internalization. Surface VEGFR1, VEGFR2, and NRP1 were labeled with 0.25 mg.ml^-1^ of the membrane-impermeant biotinylation reagent EZ-link Sulfo-NHS-SS-Biotin (ThermoFisher Scientific) at 4°C for 30 minutes with constant rocking. The unreacted biotinylation reagent was quenched by incubation with 50 mM glycine. HUVECs were washed twice with ice-cold PBS and removed from the plate by gently scraping. After centrifugation at 800g for 5 minutes at 4°C, cells were lysed using the provided lysis buffer supplemented with protease/phosphatase inhibitors (Cell Signaling Technology). A percentage of the total lysate was reserved for total protein analysis, and the rest was put over a NeutrAvidin Agarose slurry-containing spin column (ThermoFisher Scientific) at 4°C for 1 hour. Flow-through was collected in a given volume (same for all samples), and the column was then eluted with 1 M Dithiothreitol (DTT), and the eluate was resuspended in the same volume as the flow-through, to facilitate determination of surface:internal ratios. Experiments were repeated three times.

#### Ligand stimulation

HUVECs were starved in EBM-2 with 0.1% serum for 12 hours prior to ligand addition. Ligand stimulation by 50 ng.mL^-1^ of either Human VEGF_165a_ or PLGF_1_ (Genscript #Z02689, RnD #264-PGB-010) for 0 min, 15 min, 30 min, 60 min, 120 min and 240 min was followed by cell surface biotinylation, lysis and immunoblotting.

#### Immunoblots

Immunoblots were performed as previously described (Boucher *et al*, 2017). Briefly, whole cell lysates, or cell surface and internal fractions, were collected as described above. For whole cell lysates, approximately 10 µg of protein was separated by SDS-PAGE on 10% Tris-Acetate gels and transferred onto PVDF membranes. For biotin labeling experiments, equal volumes of the flow-through (∼10µg) and eluate were loaded to facilitate determination of surface:internal ratios. Membranes were blocked for 1 hour in OneBlock (Prometheus) and incubated at 4°C with primary antibodies (Supplementary Table S19) in OneBlock overnight. Membranes were washed 3 times in PBST (PBS 0.1% Tween-20) before adding HRP-conjugated secondary antibodies for 1 hour at room temperature. Secondary antibodies were removed, and membranes washed four times in PBST before addition of Luminata Forte (Millipore). See Supplementary Table S7 for details of antibodies, inhibitors and siRNAs used for this study. BioRad software was used to image the western blots, and ImageJ software was used to isolate and quantify the specific bands produced by the antibody. Loading control (typically ɑ-tubulin) was used to normalize the amount of total protein present.

## Results

### Impact of VEGF_165a_ and PLGF_1_ on VEGF receptor trafficking in HUVECs

We previously estimated the values of 15 receptor trafficking parameters and 3 receptor production rate parameters in the absence of exogenous ligands in HUVECs (Sarabipour *et al*, 2024). To study the impact on receptor trafficking of PLGF_1_ binding to VEGFR1 and VEGF_165a_ binding to VEGFR1, VEGFR2, and NRP1, we measured whole-cell expression levels of VEGFR1, VEGFR2, and NRP1 in HUVECs following addition of these exogenous ligands. Cells were serum starved for 12 hours, then treated with 50 ng.mL^-1^ of VEGF_165a_ or PLGF_1_ for 15 min, 30 min, 1 hr, 2 hr, and 4 hr, and compared to the no-ligand-added case. Over time, we did not observe consistent changes in whole-cell levels of VEGFR1 or NRP1 under treatment with either ligand (Figure 2A-D). Whole-cell VEGFR2 levels seemed lower at later times (down 42% and 31% after 2 hours and 4 hours) following VEGF_165a_, but with high variability between replicates (Figure 2A, 2C); there was no similar response to PLGF_1_ (Figure 2B, 2D).

**Figure 2.**
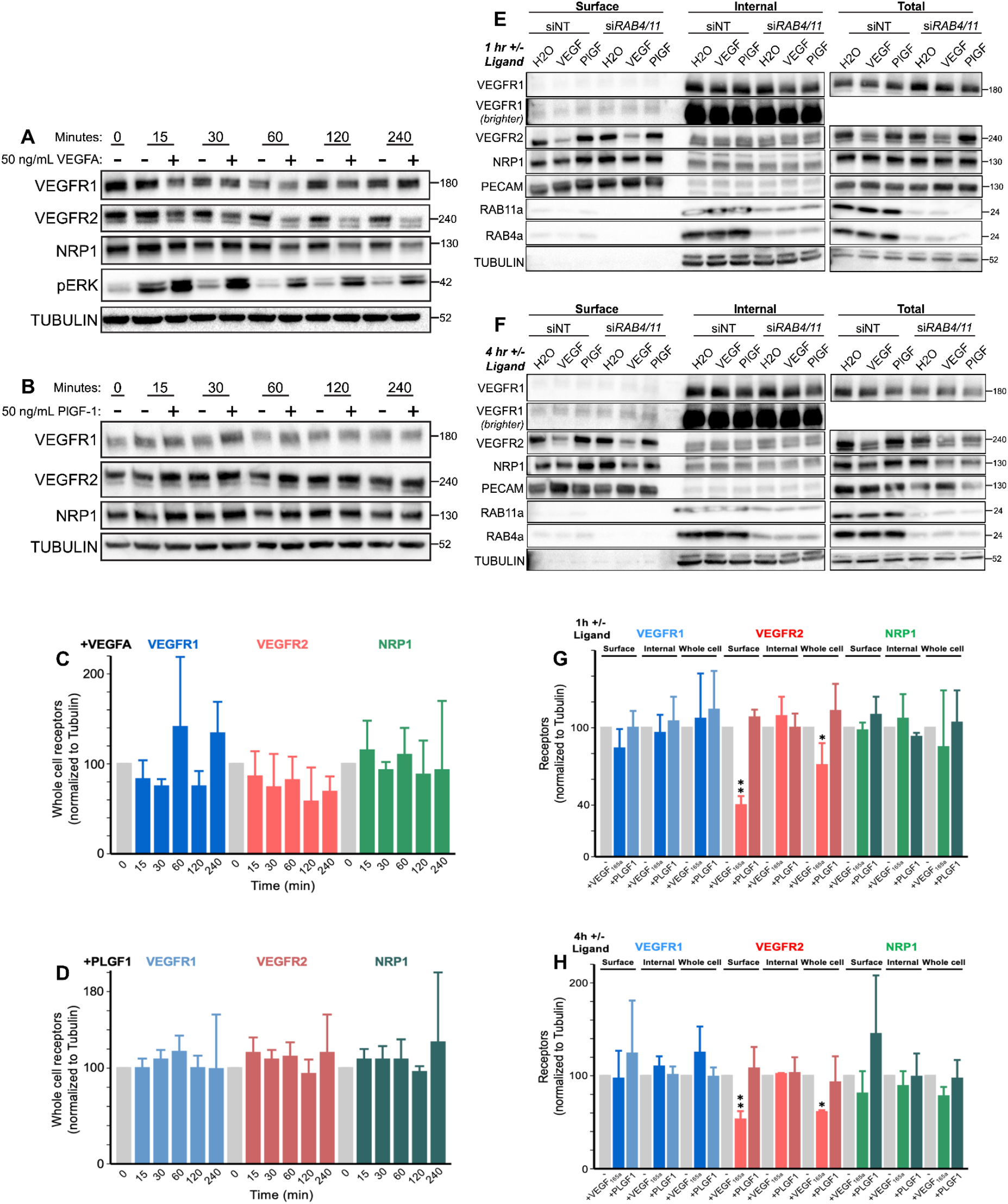
Experimental measures of whole-cell and surface levels of VEGFR1, VEGFR2, and NRP1 in HUVECs. Western blot of total HUVEC lysates treated as indicated for **A,** 1 and **B,** 4 hours with 50 µg.mL^-1^ of VEGF_165a_ or PLGF_1_ and stained for VEGFR1, VEGFR2 and NRP1. Representative of n=3 replicates. **C, D,** Quantification of whole cell receptor levels shown in A,B. **E, F,** Western blot of Biotin labeling assay to measure surface and internal VEGFR1, VEGFR2, and NRP1 levels in HUVECs; for 1 hour and 4 hours with 50 µg.mL^-1^ of VEGF_165a_ or PLGF_1_ and stained for VEGFR1, VEGFR2 and NRP1. Representative of n=3 replicates. Western blot showing the effect on VEGFR1, VEGFR2 and NRP1 levels of depletion of both Rab4a and Rab11a for 1 hour and 4 hours ligand treatment; Representative of n=3 replicates. Reagents used for the experiments are detailed in Supplementary Table S19. **G, H,** Quantification of whole cell receptor levels shown in D,E. Single factor ANOVA analysis for ligated whole cell receptor levels shown in panels C-D did not yield significant differences between receptor levels at different lysis timepoints (p>0.5), however the same analysis for panels G-H revealed significant decreases in ligated receptor levels at 1 hour and 4 hours for surface VEGFR2 and whole cell VEGFR2 receptor levels following VEGF_165a_ addition compared to no ligand addition (*, p<0.01; **, p<0.0001).

We previously showed that whole-cell receptor expression does not tell the full story; quantifying changes to receptor levels in different cellular compartments can reveal trafficking. To quantify receptor levels in the cellular compartments, we used surface biotin labeling to separate and quantify the surface and internal pools of NRP1, VEGFR2, and VEGFR1 in the absence and presence of ligands. Surface biotinylation labeled total surface proteins on serum-starved HUVECs, and these labeled proteins were then captured on streptavidin beads. We then used western blotting to quantify the levels of surface (biotinylated) and intracellular (non-biotinylated) NRP1 (Figure 2E-H). From this, we can see that surface VEGFR2 levels decrease following VEGF_165a_ treatment and stay decreased over time, consistent with other published observations of VEGFR2 decreases following ligand addition (Fearnley *et al*, 2016; Kofler *et al*, 2018). Interestingly, the internal receptor levels increase only slightly and transiently, returning to pre-ligand levels; and the total receptor levels are decreased (but by a lesser amount than the surface VEGFR2). By contrast, VEGFR1 and NRP1 levels, both on the surface and intracellularly, were not significantly changed by either VEGF_165a_ or PLGF_1_ treatment; and VEGFR2’s surface and internal levels were also unchanged by PLGF_1_.

Given the involvement of Rab4a and Rab11a in recycling (Figure 1), we also explored the effect of Rab4a and Rab11a knockdown via siRNA on receptor expression following ligand treatment (siRNA was administered 18 hours before the ligand addition, to allow time for the knockdown of Rab4a and Rab11a proteins (Figure 2E-2F). These knockdowns did not substantially affect localized receptor expression levels.

Moving to the computational model, we explored potential explanations for the experimentally-observed temporal differences in localized VEGFR2 levels. We first explored increased internalization of VEGF_165a_-bound VEGFR2 (compared to unligated VEGFR2), because this is one of the known effects of VEGFR2 activation, typically understood to be due to enhanced association of phosphorylated VEGFR2 with clathrin internalization pathways (Bhattacharya *et al*, 2005; Bruns *et al*, 2010, 2012; Ewan *et al*, 2006; Gourlaouen *et al*, 2013; Lampugnani *et al*, 2006; Lee *et al*, 2014; Nakayama *et al*, 2013). Testing different levels of increase to the internalization rate constant, we found that the lower surface levels of VEGFR2 are consistent with ligand-activated VEGFR2 having an internalization rate constant three times that of unactivated VEGFR2 (Figure 3A). As validation, we also compared the simulation-predicted impact of ligand binding on the intracellular and whole cell VEGFR2 levels to our experimental measurements (Figure 3B and 3C respectively), and these were consistent; we found that the transient increase in internal VEGFR2 levels and decreased whole cell VEGFR2 levels are recreated by the same parameter change. At first, the transient internal VEGFR2 change was somewhat surprising; since the increased internalization depletes VEGFR2 surface levels, we might expect compensatory increased internal VEGFR2 levels. However, what the model suggests is happening is that the increased internalization briefly increases the internal levels, but then degradation returns the internal VEGFR2 to pre-ligand levels. The model predicts this transient behavior without any change to the intrinsic degradation rate constant; rather, the higher VEGFR2 levels themselves increase the rate, and the driver for internal steady state levels (both before ligand and after ligand) is the balance of receptor production (assumed not to change) and degradation. Thus, the internal levels have to return to pre-ligand levels even with the increased internalization. This kind of nonobvious mechanistic insight is part of why experimental-computational combined approaches are powerful.

**Figure 3.**
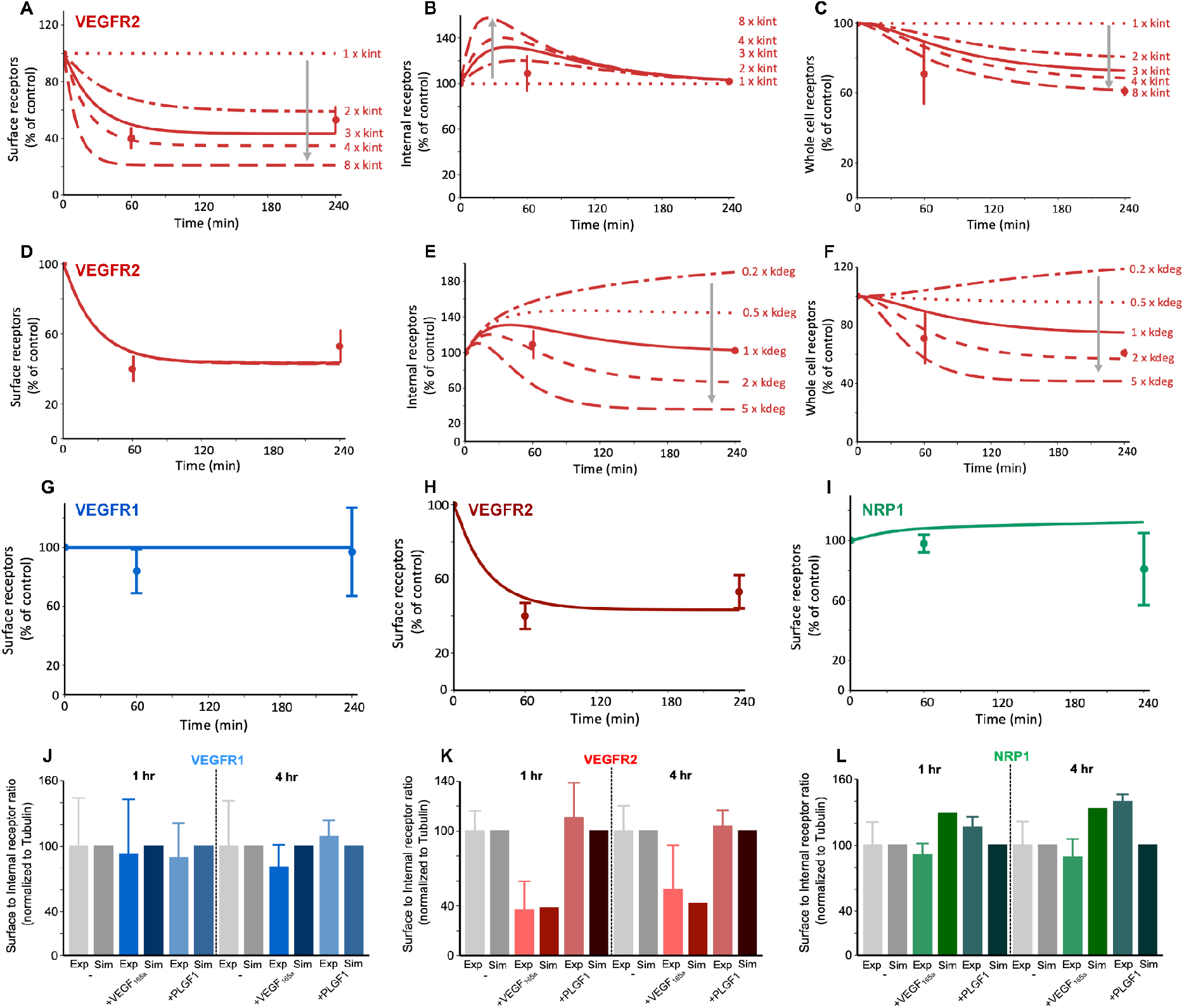
Ligand treatment - here, VEGF_165a_ - alters VEGFR2 distribution but not VEGFR1 or NRP1 distribution. **A-C**, total (ligated and unligated) receptor levels on the cell surface, inside the cell, and combined (whole cell) following VEGF_165a_ treatment. The experimentally-observed decline in surface and whole cell levels, simultaneous with a stable internal level, is consistent with a VEGF-induced three-fold increase in VEGFR2 internalization (V.R2.kint), compared to the unligated receptors, lines represent 8x, 4x, 3x, 2x, 1x, **D-F,** total (ligated and unligated) receptor levels on the cell surface, inside the cell, and combined (whole cell) following VEGF_165a_ treatment. Using the increased VEGFR2 internalization as a baseline, changes to ligand-induced degradation of VEGFR2 (V.R2.kdeg) do not improve the fit to experimental observations, lines represent 5x, 2x, 1x, 0.5x, 0.2x, **G-I,** the simulated results with enhanced VEGFR2 internalization shows a match between the predicted and experimentally changes in surface receptor levels for VEGFR1, VEGFR2, and NRP1. **J-L,** Experimental and simulated surface to internal ratios for VEGFR1, VEGFR2 and NRP1 in the absence of ligands and under 1h and 4h ligand treatment conditions (normalized to the unligated values). See Figures S2-S7.

Among the various trafficking parameters, degradation was the only other parameter to which VEGFR2 levels were sensitive in our previous ligand-free study (Sarabipour *et al*, 2024), because VEGFR2 recycling was not quantitatively important. Thus, we checked whether, along with or instead of VEGFR2 internalization, ligand-modified VEGFR2 degradation rate constants could explain the observations; however, altering the degradation rate constant did not affect surface VEGFR2 levels (Figure 3D), and thus could not explain those levels, and furthermore it resulted in long-term changes to internal VEGFR2 levels (Figure 3E), and thus could not explain those observations either. Altering the recycling rate constants for ligand-bound VEGFR2 did not affect VEGFR2 levels at any location, as expected (Supplementary Figure S2G-O).

Simulating the potential impact of ligands on VEGFR1 trafficking (Supplementary Figures S3-S4) has some similarities and some differences to VEGFR2. For example, like VEGFR2, increased internalization would decrease surface VEGFR1 (Figure S3A, S4A), but it would have even less impact on internal and whole cell VEGFR1 than on VEGFR2, because most of VEGFR1 is already internal (Sarabipour *et al*, 2024). Ligand-induced alterations to the degradation rate of VEGFR1 would be predicted to impact both surface and internal levels of VEGFR1 (Figure S3D-F, S4D-F); increased degradation lowers internal levels, which decreases the number of receptors available for recycling to the surface. That’s different from VEGFR2, for which the surface level depends only on production and internalization, because recycling is low in these cells. Increasing recycling rates increase surface VEGFR1 levels (though the second step of Rab11a-recycling is not the bottleneck and therefore sees no change (Figure S3M-O)).

To explore this more systematically, we performed simulations varying the values of individual trafficking rate constants and evaluated receptor levels at each location, to identify parameters that most strongly affected model outputs (Supplementary Figures S5-S6). Since we know that in the experiments we didn’t observe substantial changes to whole cell, surface, and internal levels of either VEGFR1 or NRP1, we assumed that the trafficking rate constants for these receptors - at least those that in simulations result in changes to the observed metrics - are unaffected by ligand binding. We should note, however, that because the amount of VEGFR1 on the surface is already low (10% or less of cellular VEGFR1; (Sarabipour *et al*, 2024)), the effect of changes to the internalization rate constant for VEGFR1 (Figure S4) may not show up in experimental data - first, because the impact on surface VEGFR1 would be to deplete an already hard-to-measure low level; and second, because the effect on internal and whole cell VEGFR1 is so small (even smaller than the effect for VEGFR2). Therefore, we proceed with an assumption that VEGFR1 internalization rate is not affected by ligand binding, and while we can’t rule out that this may be incorrect, if it is then the level of effect on the already-skewed VEGFR1 localization is expected to be negligible. Overall, the picture that emerges is that there are no trafficking alterations by ligand binding needed to match the observations for VEGFR1 or NRP1, and thus the most parsimonious and consistent conclusion is that only the internalization of VEGFR2 is affected, as evidenced by its effects on VEGFR2 levels and their consistency with experimental data (Figure 3).

As another way of looking at these trafficking changes, we calculated the overall transport rates (i.e., rate constant multiplied by concentration) for the receptors moving between subcellular locations (Supplementary Figure S7). The overall rates of VEGFR1 movement are higher than VEGFR2 but are unaffected by ligand addition. VEGFR2 internalization and degradation are transiently increased (1 hour, Figure S7E) until a new steady state is established with different receptor localization (4 hours, Figure S7H). Ligand addition also has an impact on NRP1 trafficking rates as NRP1 coupling to VEGFR2 via the ligand will exhibit slower transport than NRP1 coupled to VEGFR1 (Figure S7C,F,I). However, overall trafficking remains balanced and the levels of NRP1 in different locations are not predicted to change substantially (Figure 3I, 3L), which matches the experimental measurements (Figure 2, Figure 3I, 3L).

As additional validation of the model, we simulated disruption to the Rab4a/Rab11a recycling pathways (Supplementary Figure S8) and compared it to experimental siRNA-mediated knockdown of both Rab4a and Rab11a in HUVECs (Figure 2E-H). Expression of VEGFR1, VEGFR2, and NRP1 was measured following siRNA-Rab4a/siRNA-Rab11a treatment, in the absence and presence of 50 ng.mL^-1^ of VEGF_165a_ or PLGF_1_ at 1 hour and 4 hour timepoints following knockdown and serum starvation to assess changes to stability/turnover. The surface, internal or whole-cell levels of VEGFR1, VEGFR2, and NRP1 were essentially unaffected by Rab4a/11a knockdown experiments (Figure 2E-F) and in the equivalent simulations (Supplementary Figure S8), receptor stability in response to ligands was similarly unaffected by the Rab4a/Rab11a knockdown.

### Impact of receptor trafficking on ligand-induced receptor activation

To this point we have focused on receptor localization, counting both ligated and unligated receptors. Signaling is generally initiated by ligated, activated receptors, and thus we explored how trafficking of the three different receptors might result in different localized levels of active receptors. This differential localization of active receptors is important, given previous studies supporting the concept that signal initiation in different subcellular locations can result in activation of different signaling pathways and different cell behaviors (Wendel Clegg & Mac Gabhann, 2015; Lampugnani *et al*, 2006; Lamalice *et al*, 2006; Tan *et al*, 2013; Goh & Sorkin, 2013).

We define active VEGFR1 and VEGFR2 as being in complexes where two receptors are bound to the same bivalent ligand (receptor-ligand-receptor complexes). Activation of receptors may have additional complexity, including the differential localization of phosphatases that can site-selectively dephosphorylate tyrosines in the intracellular domains of the receptors, but here we focus on the potential for ligand-induced receptor activation by location. We simulated four ligands: VEGF_165a_ and PLGF1 based on the experimental data, plus VEGF_121a_ and PLGF_2_ using the same trafficking rates as their isoforms but reflecting the distinct receptor-binding capabilities as described earlier (Supplementary Table S1).

VEGFR2 on the surface of the cells first encounters ligands that are outside the cell, and on binding forms signal-capable complexes; as these complexes internalize faster than other VEGFR2, these complexes then decline over time (Figure 4A). The active complexes moved inside (Figure 4B), and ultimately more signal-capable complexes are present inside the cell than on the surface of the cell (Figure 4A-C), in contrast to the initial 50-50 distribution of VEGFR2 between the cell surface and inside the cell. Signal-capable complexes decline over time, but slowly, as degradation inside the cell is mostly balanced by new receptor production, and there is a plentiful supply of ligands in the extracellular media to bind to these new receptors. The apparent higher activation of VEGFR2 by VEGF_121a_ vs VEGF_165a_ is largely due to higher initial molar ligand concentrations (in nM) for the same weight-based concentrations (in ng/ml) as VEGF_121a_ has a lower molecular weight (Table S12).

**Figure 4.**
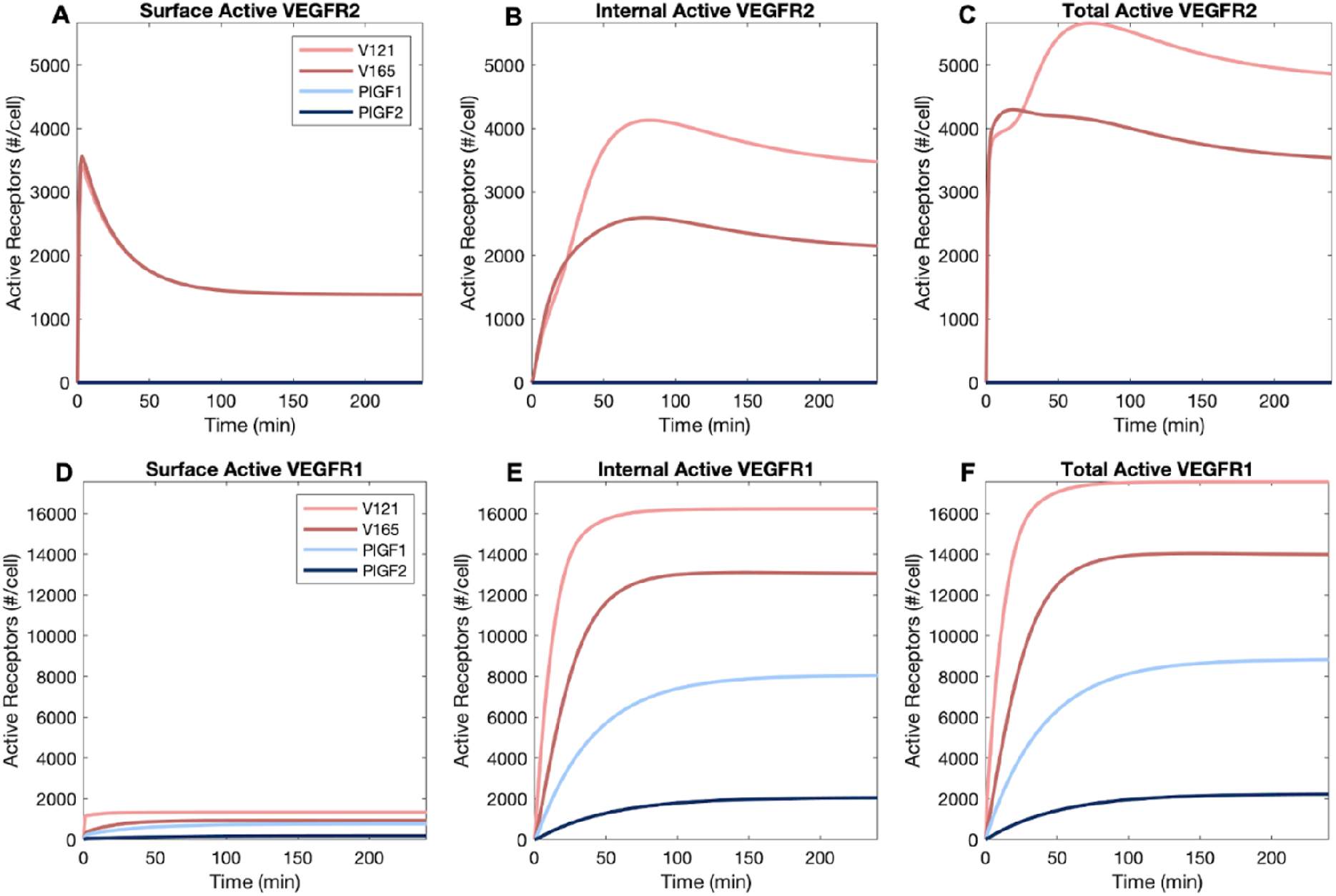
Distribution of ligated and active VEGFR1 and VEGFR2 receptors following addition of one ligand. **A-C**, Cell surface (A), internal (B), and whole cell (C) VEGFR2 following 50 ng.ml^-1^ VEGF_121a_ or VEGF_165a_, **D-F,** cell surface (D), internal (E), and whole cell (F) VEGFR1 following 50 ng.mL^-1^ VEGF_121a_, VEGF_165a_, PLGF_1_ or PLGF_2_.

As for VEGFR1, we know that the expression of the receptor on the surface is very low, and as a result few active receptor complexes are formed (Figure 4D). The question then is, with the great majority of VEGFR1 inside the cell, whether a sufficient ligand can internalize and ligate those receptors? Clearly, the answer is yes (Figure 4E). The exchange on VEGFR1 via internalization and recycling is sufficient to bring ligated receptors into the cell where they account for the vast majority of the ligated VEGFR1 across the whole cell (Figure 4D-F). We considered that perhaps ligands enter the cell via binding the more numerous surface VEGFR2 and then rebinding to internal VEGFR1 once inside the cell, and likely this does contribute, but it is not required, as PLGF_1_ binds only to VEGFR1 (Supplementary Table S1) yet shows significant internal ligation (Figure 4E). PLGF_2_ is predicted to induce less activation than PLGF_1_, largely due to inability to bind the R1-N1 complex (Supplementary Table S1).

The bias towards intracellular active receptor complexes is illustrated more clearly when compared side-by-side after 1 hour (Figure 5A) and 4 hours (Figure 5B) of ligand stimulation. Another consequence active receptor internalization is that ligands can unbind from (and rebind to) receptors while inside the cell, resulting in the presence of free (unbound) ligands inside endosomes (Figure 5C-D) and a different equilibrium between ligands and receptors inside the cell than pertains at the cell surface.

**Figure 5.**
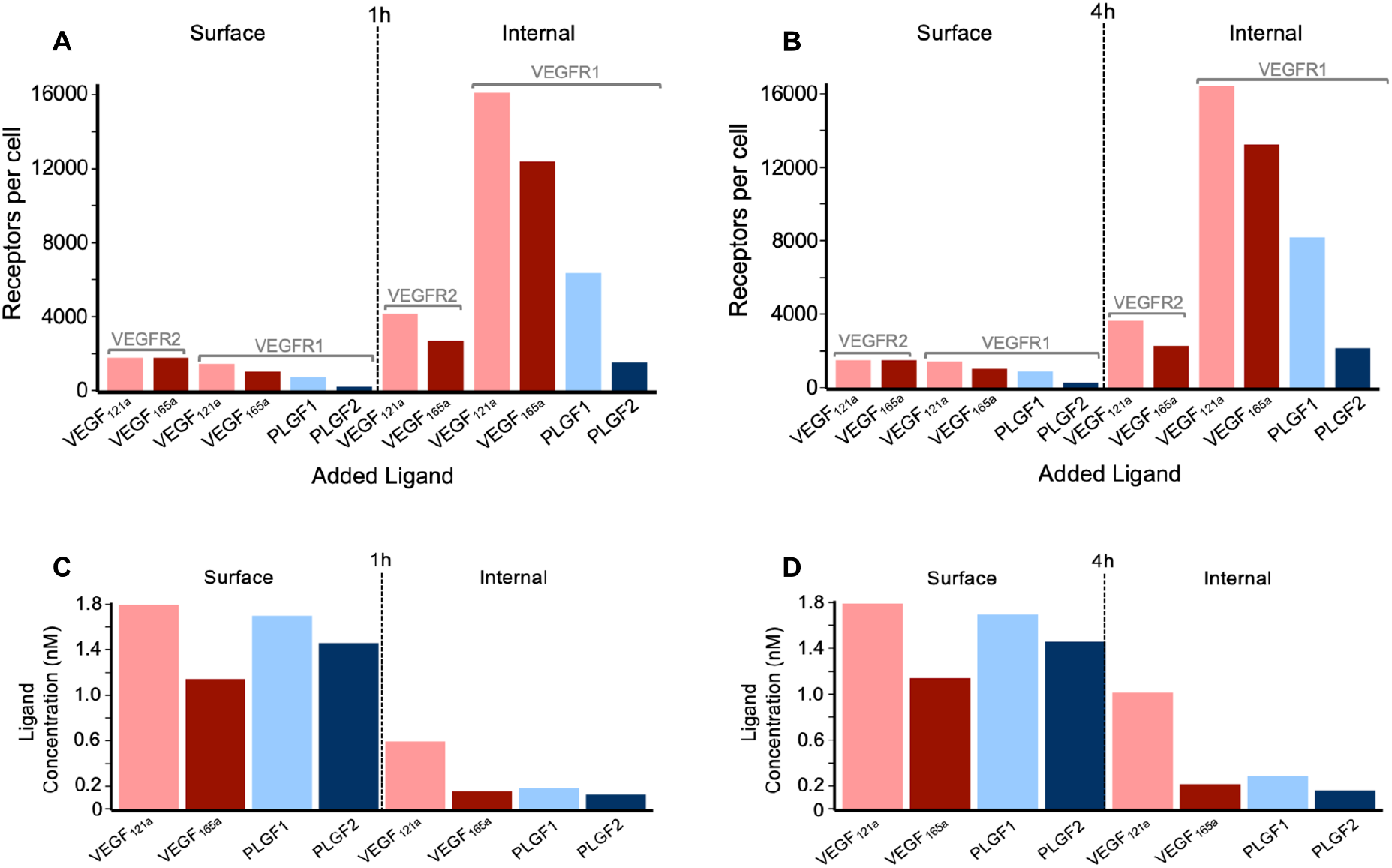
Distribution of ligands and of active ligated VEGF receptors. **A, B,** Levels of active VEGFR1 and VEGFR2 on the cell surface and internally, following 1 hour (A) or 4 hour (B) treatment with 50 ng.mL^-1^ of VEGF_165a_, VEGF_121a_, PLGF_1_, or PLGF_2_. **C, D,** Concentration of extracellular ligand (“surface”) and intracellular ligand (“internal”) following 1 hour (C) or 4 hour (D) treatment with 50 ng.mL^-1^ of VEGF_165a_, VEGF_121a_, PLGF_1_, or PLGF_2_. See the dose-dependent impacts on these levels in Figures S13-S14.

We also ran simulations across a range of extracellular ligand doses, showing similar observations across the range of concentrations - e.g., higher surface VEGFR2 activation at earlier times and decrease at later times (Supplementary Figure S9), and high levels of internal VEGFR1 activation (Supplementary Figure S10). At lower concentrations, VEGF_165a_ has an advantage over VEGF_121a_ in VEGFR2 activation due to its ability to bind to NRP1 (Supplementary Figure S9), while at higher concentrations, as receptors get saturated, VEGF_121a_ becomes a more potent inducer of VEGFR2. VEGF_121a_ is predicted to induce higher VEGFR1 activation than VEGF_165a_ (Supplementary Figure S10), due to the high levels of NRP1-VEGFR1 heterodimers, which permit binding of VEGF_121a_ but not VEGF_165a_ (Supplementary Table S1). For similar reasons, PLGF_1_ is predicted to be a stronger inducer of VEGFR1 than PLGF_2_ (Supplementary Figure S10). An alternate illustration of both the dose-dependency and the internal bias is shown in Supplementary Figures S11 and S12.

One notable feature of the ligand dose-dependence curves (Figures S9, S10) is that at high concentrations, there is a dip in several of the curves, i.e., increasing the added ligand concentration further decreases receptor ligation and activation. This phenomenon can also be seen when exploring the dimerization of receptors (Supplementary Figure S13): in the absence of exogenous ligands, some VEGFR1 and VEGFR2 are dimerized (though inactive). As ligands are added, at low ligand concentrations, ligands bind receptor monomers (or bind to receptor dimers but only once) not increasing dimerization; then at intermediate concentrations, dimerization increases alongside activation; and at higher ligand levels, receptor monomers become saturated with bivalent ligand and increased ligand binding decreases dimerization. The heterodimers formed by VEGFR1 and NRP1 exhibit only slightly increasing prevalence in the presence of ligands that bind the heterodimers (Supplementary Figure S14), while they decrease at higher concentrations of ligands that do not (e.g., VEGF_165_).

#### Decoy effects and the impact of competition

As noted earlier, VEGFR1 may have both intrinsic signaling function and modulate VEGFR2 signaling via decoy function. This ‘decoy effect’ is due to competition for ligand binding; ligands that bind VEGFR1 decrease the available pool of ligands for binding to VEGFR2. To quantify the likely size of the decoy effect, we predicted the number of active ligated VEGFR2 induced by VEGF_165_, both with and without the presence of VEGFR1. In this way, we estimated the size of VEGFR1’s decoy effect on VEGFR2, the calculations for which are described in *Methods*.

Aggregating across the whole cell, in the presence of VEGFR1 (Figure 6A) increasing VEGF levels result in higher levels of VEGFR2 signal-initiation complexes, resulting in thousands of active VEGFR2 per cell. The early peak and partial decline over time is also visible. Without VEGFR1 (Figure 6B), the same patterns are present, however the overall level of active VEGFR2 is higher; the removal of VEGFR1 appears to have freed up ligand that now can bind to and activate VEGFR2. The net effect of this is strongest at lower VEGF concentrations (Figure 6C); this makes sense, as (a) the higher the VEGF concentration in the media, the smaller the impact that VEGFR1 binding has on VEGF concentration, and consequently on the binding to VEGFR2; and (b) at higher concentrations, VEGFR2 is already close to saturation and so additional ligand will not add many additional active complexes.

**Figure 6.**
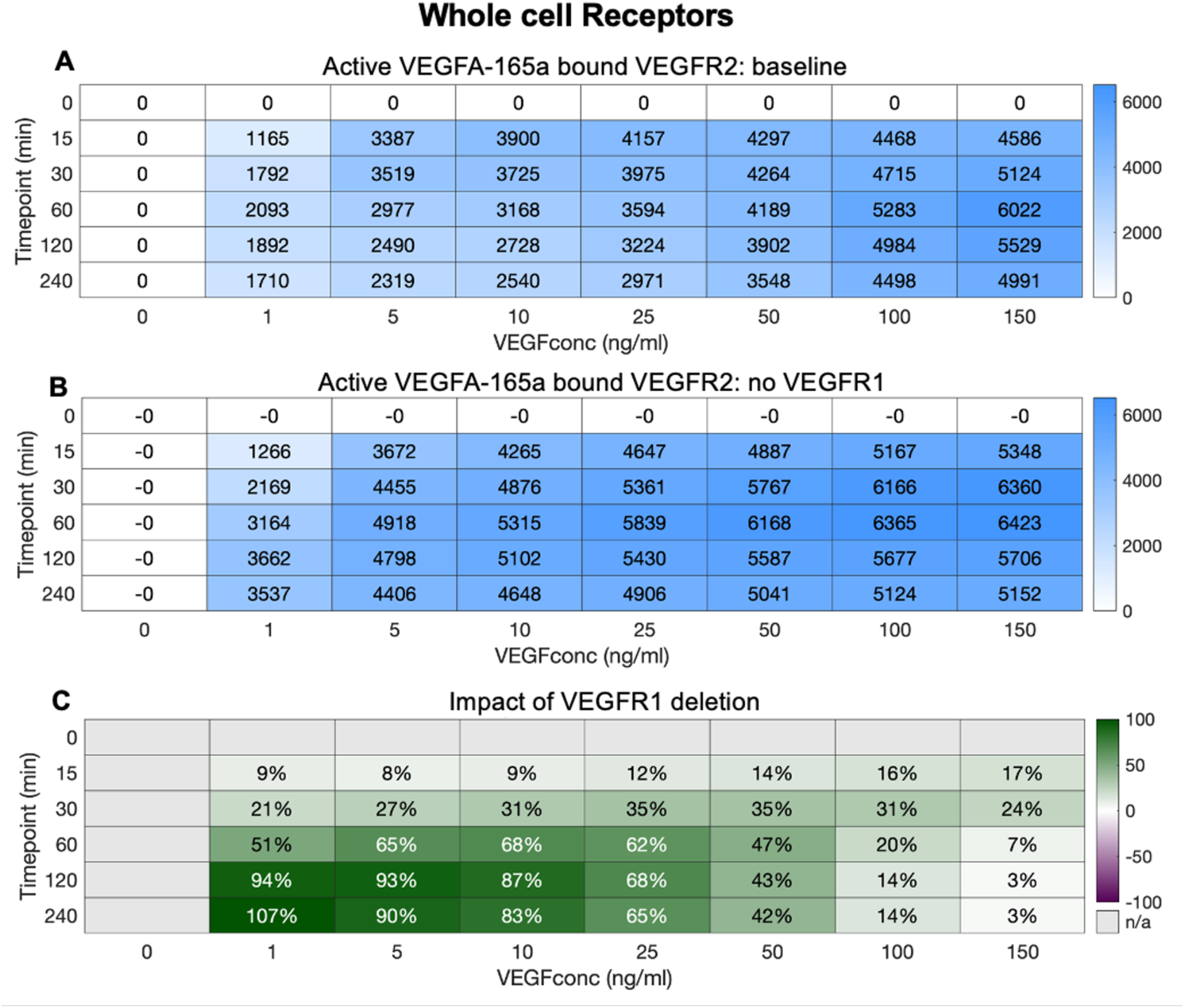
Impact of VEGFR1 on whole cell VEGFR2 ligation by VEGF_165a_. **A**, changes in VEGFR2.VEGF_165a_.VEGFR2 levels under varying VEGF_165a_ concentrations **B,** changes in VEGFR2.VEGF_165a_.VEGFR2 in the absence of VEGFR1 levels under varying VEGF_165a_ concentrations **C,** percent change in VEGFR2.VEGF_165a_.VEGFR2 levels under loss of VEGFR1 molecules and under varying VEGF_165a_ concentrations. When VEGFR1 is removed, whole cell VEGFR2.VEGF_165a_.VEGFR2 levels change.

However, we previously explored this decoy effect, with a model of VEGFR1-VEGFR2 competition that had less detail on receptor trafficking (Mac Gabhann & Popel, 2004; Wu *et al*, 2010b, 2010a), that predicted minimal decoy effect, and little impact of VEGFR1 on VEGFR2 ligation. So why the difference? In the previous work, receptors were only present on the cell surface; they could be dynamically produced and internalized but were considered degraded once inside the cell. In this updated model with detailed trafficking, we can separate out the effects of VEGFR1 on VEGFR2 on the surface (Figure 7C) and inside the cell (Figure 7F). What we see is entirely consistent with that previous work - on the surface, there is essentially no decoy effect. This is not just because the VEGFR1 is present at lower levels on the surface than inside the cell, though that may contribute slightly; the real distinction here is that the cell surface receptors are facing out into a large reservoir of ligand in the extracellular media. Even if the concentrations are similar outside the cell and in endosomes, the difference in volume between the two locations is crucial. VEGFR1 on the surface binding VEGF results in very little decline in the overall concentration extracellularly because, relatively, there is so much ligand available in the large extracellular volume. By contrast, VEGFR1 binding VEGF inside the cell can significantly deplete VEGF concentration in endosomes because the actual amount of VEGF there (amount = concentration times volume) is lower. And thus, the decoy effect can be seen to be a phenomenon of intracellular receptors (Figure 7F) rather than surface receptors, with a major impact on intracellular VEGFR2 activation at all but the highest (already saturating) ligand concentrations (Figure 7F). We define the impact ligand reservoir size on the decoy effect as the ‘reservoir effect’.

**Figure 7.**
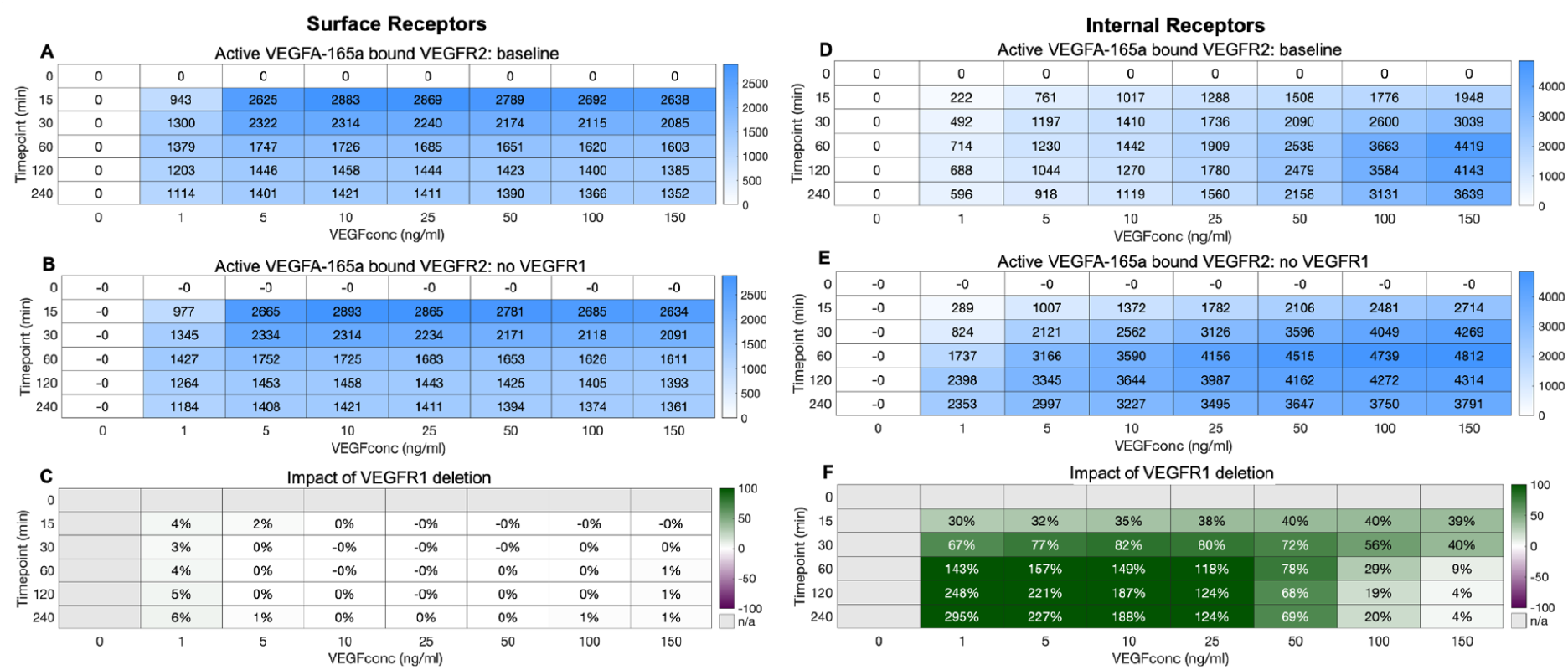
Impact of VEGFR1 on **A,B,C,** Surface and **D,E,F,** Internal VEGFR2 ligation by VEGF_165a_. Changes in VEGFR2.VEGF_165a_.VEGFR2 levels under varying VEGF_165a_ concentrations. Change in VEGFR2.VEGF_165a_.VEGFR2 levels under loss of VEGFR1 molecules and under varying VEGF_165a_ concentrations (percent change shown in B, and D,). When VEGFR1 is removed, surface VEGFR2.VEGF_165a_.VEGFR2 does not change (panel B) but internal VEGFR2.VEGF_165a_.VEGFR2 levels change over a range of VEGF_165a_ concentrations (panel D).

**Figure 8.**
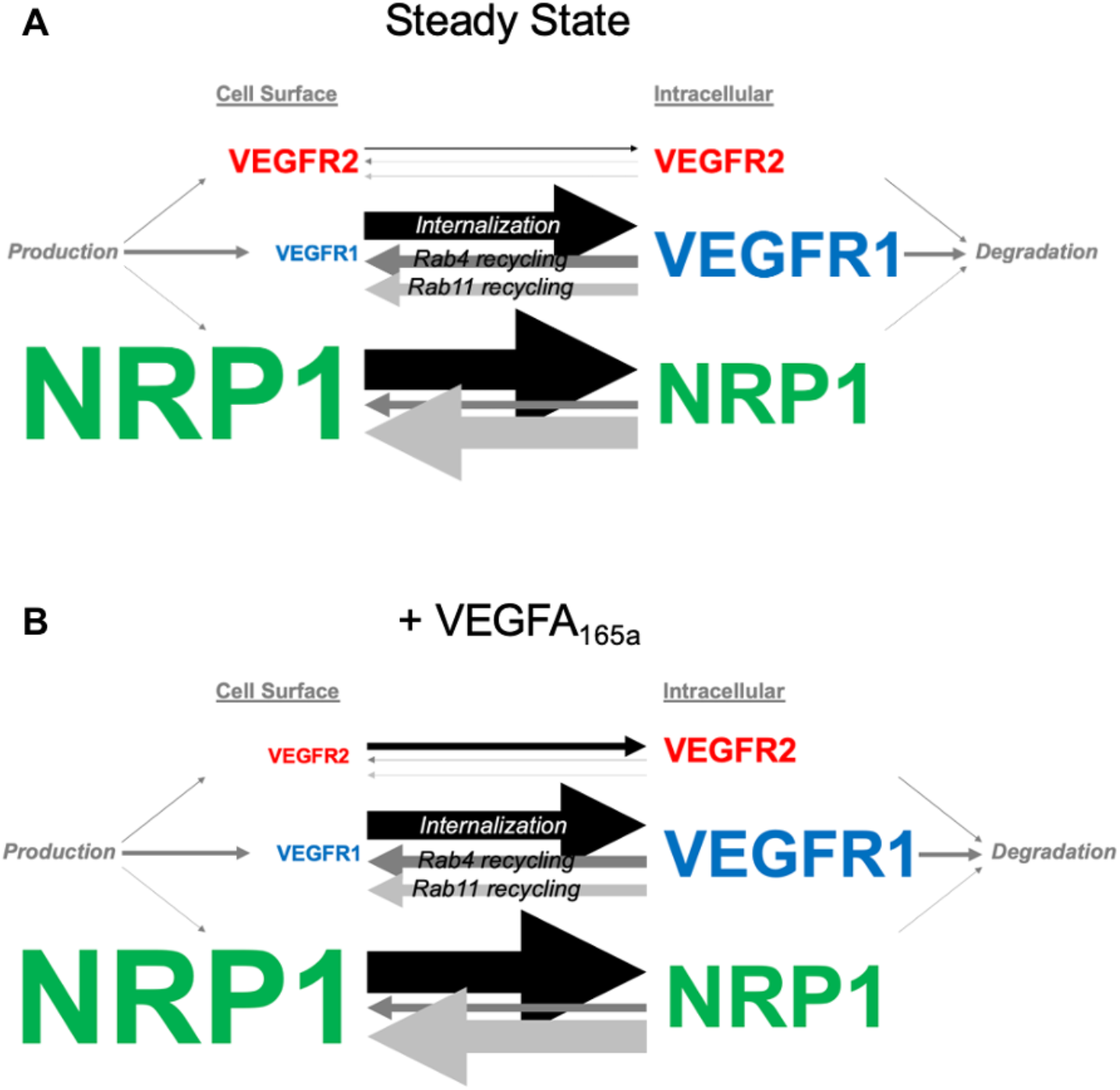
Differences in VEGFR1, VEGFR2 and NRP1 trafficking in the absence of ligands (steady state) and presence of ligands in HUVECs. **A,** VEGFR1, VEGFR2, NRP1 are constitutively trafficked in the absence of ligands in HUVECs. **B,** VEGFR2 internalization is faster as a result of VEGF-A treatment. PLGF treatment has no impact on VEGFR1, VEGFR2 or NRP1 trafficking in HUVECs. The size of the arrow represents the amount of receptor movement. The size of the receptor represents the expression level relative to other receptors shown here.

As noted earlier, NRP1 is a co-receptor for some VEGF/PLGF isoforms and not others (Supplementary Table S1) and leads to impacts on the binding of different ligands to the receptor tyrosine kinases and their resulting activation (Figures 4, 5). This can be seen to be a ligand-dose-dependent effect (Supplementary Figures S9, S10) that is stronger at lower ligand levels. We computationally deleted NRP1 and calculated the impact on VEGF_165a_ activation of VEGFR2 (Supplementary Figures S15-S17) and VEGFR1 (Supplementary Figures S18-S20). Indeed, the loss of NRP1 causes a sizable decrease in VEGFR2 activation at lower concentrations, due to the loss of the avidity provided by the VEGFR2-VEGF-NRP1 complexes. At higher ligand concentrations, there is a small but opposite effect, likely due to the ligand coming close to saturating the receptors, and the removal of NRP1 now eliminating a large sink for ligand binding. There are slight quantitative differences in these effects on the cell surface (Figure S16) vs internally (Figure S17) due to different relative levels of the receptors in these locations, but the trends are the same. By contrast, VEGF_165a_ activation of VEGFR1 is strongly enhanced by NRP1 deletion (Supplementary Figures S18-S20), because the VEGFR1-NRP1 heterodimers are no longer formed, making VEGFR1 more available for VEGF_165a_ binding. This effect is stronger at lower ligand concentrations, again because at higher concentrations the barriers to binding are more easily overcome. Putting these insights together, we can see that the presence of NRP1, at low or moderate concentrations, would be expected to enhance binding to VEGFR2 and diminish binding to VEGFR1 - both elements pushing the system in the direction of more VEGFR2 signaling. Of note, at higher ligand concentrations typical of *in vitro* studies (such as 50 ng/ml), the impact of NRP1 is much smaller; however, concentrations *in vivo* are typically considerably lower than those used *in vitro* (Clegg *et al*, 2017), and thus we might expect NRP1 to have a more sizeable effect *in vivo*.

Just as we have seen the impact of VEGFR1 on VEGFR2 activation, there can be a reciprocal impact of VEGFR2 on VEGFR1 ligand binding and activation (Supplementary Figures S21-23). Again, we see a stronger effect of VEGFR2 on VEGFR1 at lower ligand concentrations, but perhaps surprisingly, while VEGFR1 deletion increases VEGFR2 activation (Figures 6, 7), loss of VEGFR2 is predicted to decrease activation of VEGFR1. This holds both on the surface (Supplementary Figure S22) and inside the cell (Supplementary Figure S23). The effect of VEGFR2 on VEGFR1 is also due to the presence of NRP1. The presence of VEGFR2 allows the formation of VEGFR2-VEGF-NRP1 complexes, which reduces the formation of VEGFR1-NRP1 complexes that prevent the binding of VEGF_165_. Thus, with VEGFR2 deleted, VEGFR1-NRP1 complexes dominate and VEGF_165a_ binding to VEGFR1 is lower.

Having quantified the effects of receptors competing for ligand binding, we can also quantify the effects of ligands competing for receptor binding. Specifically, PLGF and VEGF both bind to VEGFR1. Our simulations thus far manipulated one ligand at a time, and under these conditions PLGF exhibits time- and dose-dependent activation of VEGFR1 (Supplementary Figure S24). However, tissues often express multiple isoforms, so we explored co-administration of two ligands (Supplementary Figures S25-28). Unlike receptors, where competition was higher at low ligand concentrations, these simulations reveal that competition is higher at high ligand concentrations, likely because at these high levels depletion of available free receptors leads to the competition effects. For example, at 5 ng/ml of both VEGF_165a_ and PLGF_1_, VEGF-bound active VEGFR1 is reduced only 3.7% by the addition of PLGF1 (Supplementary Figure S25), while this raises to 20% and 23% at 50 ng/ml and 150 ng.ml^-1^, respectively. VEGF_165a_ has a much stronger effect on PLGF_1_; VEGFR1 activated by PLGF is reduced 15%, 50%, and 68% for initial ligand concentrations of 5, 50, and 150 ng.ml^-1^ (Supplementary Figure S26). Unlike competition by receptors for ligands, competition by ligands for receptors is not primarily mediated by changes to ligand levels (Supplementary Figure S27), which reflects saturated binding (and the scarcity of unligated receptors) as the key mediator. What about the consequences of VEGF-PLGF competition on VEGFR2 activation? PLGF cannot bind VEGFR2, but if PLGF binding VEGFR1 increases the VEGF available to bind to VEGFR2, then perhaps some competition will be observed. In this case, high levels of competition (and therefore an impact of PLGF on VEGF activation of VEGFR2) is only seen at low VEGF and high PLGF concentrations (Supplementary Figure S28). At high VEGF levels, VEGFR2 is already well activated and thus not responsive to PLGF addition; at low levels of VEGF, only high levels of PLGF will sufficiently saturate VEGFR1 to encourage VEGF transfer to VEGFR2; and even this effect is only seen inside the cell, because at the cell surface the large reservoir again provides too large a buffer. Even the largest such changes are likely too small to observe reliably in an experiment.

Finally, it is known that pH can be different, typically lower, in endosomes than outside the cell and that this can alter the binding rate constants for ligand-receptor binding. We explored whether pH-based differences in ligand-receptor binding would impact some of the key insights here (Supplementary Figures S29-S30). While some small differences in the overall levels of ligand-receptor binding are observed (intracellularly) as a result, the overall system is not greatly changed, for example the decoy effects of VEGFR2 on VEGFR1 and of VEGFR1 and VEGFR2 are essentially unchanged.

## Discussion

Proteins must be in the right cellular location at the right time to interact with other proteins and function properly. For transmembrane proteins such as growth factor receptors, endosomal trafficking brings them to the surface and shuttles them between the surface and intracellular locations. These intracellular locations may facilitate receptor degradation, initiation of alternate signaling pathways, or simply be a repository for additional proteins that can come to the surface when required. Whatever their purpose, the localization of the receptors (and therefore their function) is controlled by the processes of intracellular trafficking. Receptors that bind VEGF family ligands, such as the receptor tyrosine kinases VEGFR1 and VEGFR2, and the co-receptor Neuropilin-1 (NRP1) are no different; to bind to extracellular ligands, they must be on the cell surface. Intracellular-located receptors can only bind ligands if the ligands are localized to the endosomes via autocrine production, pinocytosis, or trafficking inwards while bound to receptors.

We recently used a combined computational-experimental approach to quantify the trafficking and localization of VEGFR1, VEGFR2, and NRP1 on human endothelial cells for the first time (Sarabipour *et al*, 2024). In the absence of ligands, we showed that the receptors are constitutively moved around the cell, and that the trafficking rates and thus localization were different for each receptor. For example, VEGFR1 is less stable (has faster turnover) than VEGFR2 or NRP1; and due to the balance of trafficking processes, VEGFR1 is predominantly located inside the cell, while VEGFR2 is more balanced and NRP1 is more prevalent on the cell-surface. These localization distinctions have implications for ligand interactions, as ligand-receptor binding is considered a primary signal initiation event in many cases. However, we did not know whether ligand binding alters receptor trafficking, beyond previous observations of increased VEGFR2 internalization with ligand binding and receptor activation (Bhattacharya *et al*, 2005; Bruns *et al*, 2010, 2012; Ewan *et al*, 2006; Gourlaouen *et al*, 2013; Lampugnani *et al*, 2006; Lee *et al*, 2014; Nakayama *et al*, 2013). Therefore, the aim of this study was to explore trafficking in response to exogenously administered ligands, and to predict how the ligation of cell surface and internal receptors differs.

Using novel experimental data of VEGFR1, VEGFR2, and NRP1 expression, and additional data of VEGFR1, VEGFR2, and NRP1 localization to cell surface or internal cell locations, before and after exposure to two key VEGF family ligands (VEGF_165a_ and the VEGFR1-specific PLGF_1_), we demonstrated that of the three receptors, only VEGFR2 exhibited significantly different trafficking with ligand engagement. This is the first quantification of ligand-responsive trafficking of VEGFR1 and NRP1, and it is interesting that they did not follow the pattern of higher internalization that VEGFR2 exhibits. In addition, the pattern of the VEGFR2 changes was at first surprising: surface VEGFR2 was decreased as expected, and whole-cell VEGFR2 decreased, but internal VEGFR2 was neither substantially increased in compensation with the surface loss, nor was it decreased reflecting overall receptor loss throughout the cell. Instead, it stayed relatively steady.

Turning to our computational mechanistic model, we used separate observations to parameterize and validate the model, and to explain the mechanistic underpinnings of the observations. We used sensitivity analysis to explore how ligand-induced changes to each of the fifteen trafficking parameters might explain the outcomes. First, the surface loss of VEGFR2 could be parsimoniously explained by a single effect: three-fold increased internalization of ligand-bound VEGFR2 compared to unligated VEGFR2. Higher internalization of ligand-activated receptor tyrosine kinases has been observed in multiple receptor families, including EGFR (Starbuck & Lauffenburger, 1992; Wiley *et al*, 2003), and is thought to be due to increased association with clathrin pathways (Alfonzo-Méndez *et al*, 2022). Second, without changing any additional parameters beyond the ligand-induced internalization, the internal and whole-cell levels of VEGFR2 also matched the experimental observations. Along with validating the model, the model showed that the reason the internal receptors did not increase substantially (and only transiently, in the simulations) is that the production of the receptors is balanced by degradation, and thus in the absence of degradation changes the intracellular levels return to their pre-ligand levels quickly. The whole-cell slight decrease is due to a combination of the surface decrease and the steady internal levels. Further validation of the model was obtained by comparing simulations and experiments using siRNA to downregulate the recycling-associated proteins Rab4a and Rab11a.

Now that we have a handle on receptor trafficking, and how ligand binding affects it, what about the functional reason for studying the system - the formation of signaling-pathway-initiating ligand-receptor complexes? We gained several surprising insights from the simulations. First, active VEGFR2 are predicted to be mostly internal, even though VEGFR2 is evenly distributed between the cell surface and internally before ligand exposure, and even though all ligands in the model were initially extracellular. Second, active VEGFR1 is also mostly internal, which correlates with 90% of VEGFR1 being internal. However, the small VEGFR1 population on the surface was predicted to be sufficient to bring ligands inside the cell. The fast trafficking of VEGFR1 shuttles ligands in; and we know that it’s not just ligands internalized via VEGFR2 binding and later released inside the cell, because it works for PLGF_1_ as well. Third, NRP1 is predicted to exhibit slower trafficking due to its increased association with VEGFR2 (via VEGF_165a_), which has lower trafficking rate constants than VEGFR1 and VEGFR1-NRP1; but NRP1 levels on the surface and inside the cell are not substantially altered.

What is the meaning of predictions that active VEGFR2 and active VEGFR1 are predominantly intracellular? We know that cell signaling pathways are not solely initiated from cell surface receptors. While that has been the typical picture for illustrating ligand-induced receptor signaling, for VEGFR2 and for other receptors we know that (a) intracellular receptors can initiate signaling pathways, and (b) that intracellular receptors can initiate signaling pathways that are different than those initiated by cell surface receptors (Simons, 2012; Lampugnani *et al*, 2006; Wendel Clegg & Mac Gabhann, 2015). Just as for ligand-receptor binding, context and location are everything. Different subcellular locations host different phosphatases, scaffold proteins, and more to enable differential signaling. Overall observed downstream signaling and cellular behavior is due to a combination of these different signaling pathways following ligand exposure. Second, it is notable that the timing of the signaling can be different - for example, VEGFR2 on the surface has an early activity peak and then declines (due to internalization), while intracellular active VEGFR2 is more sustained (Simons, 2012; Lampugnani *et al*, 2006; Wendel Clegg & Mac Gabhann, 2015). There are examples of both types of signaling responses in endothelial cells - early peaks or sustained signaling - depending on the signaling pathway and the context. These temporally regulated effects may be linked to the localization of signal initiation, although here we only simulated ligand binding and receptor activation, and did not include additional regulation such as dephosphorylation that might alter these dynamics further. The complex regulation of the downstream pathways themselves, including the known differential localization of site-specific tyrosine phosphatases (Goh & Sorkin, 2013; Lampugnani *et al*, 2006; Lamalice *et al*, 2006; Wendel Clegg & Mac Gabhann, 2015) may alter both the identify and dynamics of which pathways are activated by the receptor from which locations.

We explored the impact of NRP1 on VEGFR1 and VEGFR2 activation in multiple ways, including simulating a VEGF isoform that does not bind NRP1 (VEGF_121a_) and a PLGF isoform that does (PLGF_2_). Interestingly, the effects of NRP1 are dose dependent; for example, it is at lower concentrations, which are more typical of *in vivo* conditions than of *in vitro* cell culture, that we see enhanced activation of VEGFR2 activation by VEGF_165a_ vs the non-NRP1 binding VEGF_121a_, and we can also see this effect via a computational knockdown of NRP1 expression. NRP1 also exerts an isoform-specific effect on VEGFR1 activation, by associating directly with VEGFR1 and facilitating VEGF_121a_ and PLGF_1_ binding but not that of VEGF_165a_ or PLGF_2_; this effect is again stronger at lower ligand concentrations typical of *in vivo* conditions.

With a working multi-receptor mechanistic model, we returned to a concept that has been raised before with regard to VEGFR1 and VEGFR2, specifically that VEGFR1, whose kinase and signaling activity is difficult to document in endothelial cells, serves as a ‘decoy receptor’ for VEGFR2, i.e., that by binding VEGF ligands it takes those ligands away from VEGFR2 and lowers the signaling amplitude of the canonical receptor. There is compelling evidence that this mechanism is operative during development, as VEGFR1 lacking the cytoplasmic domain is sufficient for vascular development while total VEGFR1 deletion leads to embryonic lethality with vascular overgrowth (Fong *et al*, 1995, 1999; Hiratsuka *et al*, 1998; Kappas *et al*, 2008; Kearney *et al*, 2004; Roberts *et al*, 2004). In adult angiogenesis, VEGFR1 may play a signaling role (Nesmith *et al*, 2017; Ganta *et al*, 2018, 2016). Even while signaling, VEGFR1 may function as a decoy, and thus we developed a metric to quantify this decoy role using computational simulations with and without VEGFR1 present.

Several interesting observations followed. First, in our simulations, VEGFR1 did function as a decoy, reducing the local VEGF available for VEGFR2 to bind and become activated. However, this only occurred intracellularly in our model, where the local VEGF reservoir, even at similar concentration levels, is much smaller than the extracellular reservoir (due to smaller intracellular volumes). We term this the ‘reservoir effect’. The decoy effect works by competing for a shared ligand pool; if ligand is not rate-limiting, there is likely no competition. In this model, in intracellular reservoirs the impact on local concentration of one ligand binding/unbinding a receptor is larger due to small volumes. In essence, VEGFR1 can exert a much stronger influence on VEGFR2 inside the cell than outside the cell. The lack of a decoy effect on the cell surface is consistent with previous explorations which did not account for intracellular trafficking (Mac Gabhann & Popel, 2004; Wu *et al*, 2010b, 2010a), and we also note that *in vivo*, the surface-area-to-volume ratio extracellularly would be much higher than it is *in vitro*, and thus some decoy effect on the surface may occur *in vivo*.

Second, the decoy effect that is observed in the in vitro simulations is much stronger at lower ligand concentrations. This is likely because at high ligand concentrations, the receptors don’t have to compete for ligand binding. The effective extracellular ligand concentration is predicted to be lower *in vivo* than concentrations used *in vitro*, and thus we would expect that the decoy effect would be substantial in that situation, possibly even at the cell surface.

There is a second type of decoy effect - rather than receptors competing for ligands, ligands can compete for binding to VEGFR1; in particular, because PLGF1 binds only VEGFR1, it has been proposed to be able to displace other VEGF ligands from VEGFR1 and thus increase binding to VEGFR2. While our simulations show that this effect can occur, unlike receptor competition it is a phenomenon of high ligand levels rather than low ligand levels. Because it is not predicted to occur at lower ligand concentrations, we expect that PLGF-VEGF competition may not be a significant contributor to decoy effects *in vivo*, though in pathological situations ligand and receptor expression may be altered and the effect may come into play.

Here, we did not include autocrine expression of ligands by the endothelial cells themselves, which can occur; for example PLGF and VEGF_165a_ have different expression patterns in vascular endothelial cells (Domigan *et al*, 2015; Lee *et al*, 2007). The exclusion is justified for *in vitro* cell cultures we are simulating here, as any autocrine expression would be swamped by the high levels of exogenous ligand added. *In vivo*, expression of VEGF ligands by endothelial cells would likely further increase intracellular activation if the ligands encounter receptors before secretion. Another aspect of VEGF biology we have not explored here is the potential for formation of VEGFR1-VEGFR2 heterodimers, which could themselves initiate different signaling pathways than either homodimer. We didn’t include it here to avoid further complexity, and because information is not currently available about the relative prevalence of these heterodimers, but also because the two major cellular locations (surface and internal) have two different receptor-expressing regimes: VEGFR2 in excess of VEGFR1 on the surface, VEGFR1 in excess of VEGFR2 intracellularly. This reduces the number of heterodimers that would likely form, though interestingly computational predictions suggest it may also suppress formation of the homodimer of the less abundant receptor (Mac Gabhann & Popel, 2007).

In summary, we have developed and validated a model of multi-VEGF-receptor trafficking under conditions of ligand stimulation in human endothelial cells. A similar approach could be used for other receptor systems and cell types. Many of the insights developed here for the VEGF-VEGFR system are likely also generalizable to other systems, including: the relative importance of receptor vs ligand competition at low or high ligand concentrations; competition being higher in small-reservoir environments such as intracellular spaces; and the importance of receptor localization on receptor function.

## Supporting information

Supplemental File_Sarabipour_etal_29Sep2024

## Author contributions

SS and FMG designed and wrote the code, performed data visualization, and wrote the original draft of the manuscript. KK, KMQ, and VLB planned and KK and KMQ performed the experiments and experimental data analysis. FMG, VLB, and BHA acquired resources and funding. SS, FMG, KMQ, KK, VLB, AKK, and BHA performed conceptualization, formal analysis, investigation, methodology, validation, and wrote, reviewed and edited the manuscript.

## Acknowledgements

This work was supported by National Institutes of Health grants R01-GM129074 (FMG, BHA, VLB), R01-HL101200 (BHA, FMG), and R35-HL139950 (VLB).

## Supplementary File

Contains 30 Figures and 19 Tables.

