## Supplemental File_Sarabipour_etal_29Sep2024 for "Impact of ligand binding on VEGFR1, VEGFR2, and NRP1 localization in human endothelial cells"

**Supplementary Table S1. Ligand-receptor interactions.** While all four ligands bind VEGFR1, PLGF ligands do not bind VEGFR2. Longer isoforms include a NRP1-binding domain. VEGFR1 and NRP1 can directly interact, and the coupled receptor does not admit the NRP1-binding ligands, rather the non-NRP1-binding ligands can bind to the VEGFR1 in the complex.

| | **VEGFR1** | **VEGFR2** | **NRP1** | **VEGFR1-NRP1** |
|---|---|---|---|---|
| **$VEGFA_{121a}$** | **X** | **X** | | **X** |
| **$VEGFA_{165a}$** | **X** | **X** | **X** | |
| **$PLGF_1$** | **X** | | | **X** |
| **$PLGF_2$** | **X** | | **X** | |

**Supplementary Table S2. Molecules included in the model.** The total number of molecules and molecular complexes in the model is 281. The comprehensive lists of these molecules, and the number by which each is identified in the code, are given in Tables S3-S11.

| Type | Number of complexes | See Table |
|---|---|---|
| Unligated Receptors and Receptor Complexes | 32 | S3 |
| Unbound Ligands | 16 | S4 |
| Ligand-bound monomeric R1 or R2 | 32 | S5 |
| Nonsignaling ligand-bound VEGFR1 dimers | 48 | S6 |
| Nonsignaling ligand-bound VEGFR2 dimers | 24 | S7 |
| Nonsignaling NRP1-only complexes | 16 | S8 |
| Signaling ligand-bound VEGFR1 dimers | 64 | S9 |
| Signaling ligand-bound VEGFR2 dimers | 32 | S10 |
| Matrix-bound ligands and receptors | 17 | S11 |

**Supplementary Table S3. Unligated receptors and receptor complexes.** This table gives the number by which each molecule or molecular complex is identified in the model code. Dots indicate direct binding. These unligated receptors were included in our previous model of the trafficking of only receptors (Sarabipour *et al*, 2024).

| Molecule/Complex | Surface | Rab4a5a | Rab11a | Lysosome (degraded) |
|---|---|---|---|---|
| R1 | 5 | 22 | 70 | 77 |
| R2 | 6 | 23 | 71 | 78 |
| N1 | 7 | 24 | 72 | 79 |
| R1.R1 | 9 | 31 | 90 | 147 |
| R2.R2 | 10 | 32 | 91 | 148 |
| N1.R1 | 11 | 33 | 92 | 149 |
| N1.R1.R1 | 29 | 88 | 160 | 206 |
| N1.R1.R1.N1 | 30 | 89 | 161 | 207 |

**Supplementary Table S4. Unbound ligands.** This table gives the number by which each molecule is identified in the model code. V165 represents $VEGFA_{165a}$, V121 represents $VEGFA_{121a}$, P1 represents $PLGF_1$, and P2 represents $PLGF_2$.

| Molecule/Complex | Surface | Rab4a5a | Rab11a | Lysosome (degraded) |
|---|---|---|---|---|
| V165 | 1 | 25 | 73 | 80 |
| V121 | 2 | 28 | 76 | 83 |
| P1 | 3 | 26 | 74 | 81 |
| P2 | 4 | 27 | 75 | 82 |

**Supplementary Table S5. Ligand-bound monomeric VEGFR1 or VEGFR2.** These molecular complexes are non-signaling and primarily are intermediates to ligands binding two receptors. Dots indicate direct binding. This table gives the number by which each molecule or molecular complex is identified in the model code. V165 represents $VEGFA_{165a}$, V121 represents $VEGFA_{121a}$, P1 represents $PLGF_1$, and P2 represents $PLGF_2$.

| Molecule/Complex | Surface | Rab4a5a | Rab11a | Lysosome (degraded) |
|---|---|---|---|---|
| V165.R1 | 18 | 52 | 109 | 153 |
| V121.R1 | 19 | 53 | 110 | 154 |
| V121.R1.N1 | 46 | 103 | 172 | 219 |
| P1.R1 | 20 | 54 | 111 | 155 |
| P1.R1.N1 | 47 | 104 | 173 | 224 |
| P2.R1 | 21 | 55 | 112 | 157 |
| V165.R2 | 16 | 50 | 107 | 151 |
| V121.R2 | 17 | 51 | 108 | 152 |

**Supplementary Table S6. Non-signaling ligand-bound VEGFR1 dimers.** Although these complexes include two receptors, they are not coupled by the ligand (i.e. the ligand is bound only to one of the two receptors) and thus are not in higher-propensity signaling conformation. These are also likely to be intermediate forms. Dots and parentheses indicate direct binding; for example, P1.R1(N1).R1 means one $PLGF_1$ is bound to one VEGFR1, and this VEGFR1 is itself bound to another VEGFR1 and to a NRP1. This table gives the number by which each molecule or molecular complex is identified in the model code. V165 represents $VEGFA_{165a}$, V121 represents $VEGFA_{121a}$, P1 represents $PLGF_1$, and P2 represents $PLGF_2$.

| Molecule/Complex | Surface | Rab4a5a | Rab11a | Lysosome (degraded) |
|---|---|---|---|---|
| V165.R1.R1 | 42 | 99 | 168 | 216 |
| V165.R1.R1.N1 | 86 | 158 | 230 | 261 |
| V121.R1.R1 | 43 | 100 | 169 | 218 |
| V121.R1.R1.N1 | 66 | 143 | 202 | 220 |
| V121.R1(N1).R1 | 93 | 162 | 232 | 263 |
| V121.R1(N1).R1.N1 | 94 | 163 | 233 | 265 |
| P1.R1.R1 | 44 | 101 | 170 | 223 |
| P1.R1.R1.N1 | 68 | 145 | 204 | 225 |
| P1.R1(N1).R1 | 95 | 164 | 234 | 268 |
| P1.R1(N1).R1.N1 | 96 | 165 | 235 | 270 |
| P2.R1.R1 | 45 | 102 | 171 | 229 |
| P2.R1.R1.N1 | 87 | 159 | 231 | 273 |

**Supplementary Table S7. Non-signaling ligand-bound VEGFR2 dimers.** Although these complexes include two receptors, they are not coupled by the ligand (i.e. the ligand is bound only to one of the two receptors) and thus are not in higher-propensity signaling conformation. These are also likely to be intermediate forms. Dots and parentheses indicate direct binding. This table gives the number by which each molecule or molecular complex is identified in the model code. V165 represents $VEGFA_{165a}$ and V121 represents $VEGFA_{121a}$.

| Molecule/Complex | Surface | Rab4a5a | Rab11a | Lysosome (degraded) |
|---|---|---|---|---|
| N1.V165.R2 | 64 | 141 | 200 | 211 |
| N1.V165(N1).R2 | 128 | 187 | 249 | 257 |
| V165.R2.R2 | 40 | 97 | 166 | 210 |
| N1.V165.R2.R2 | 65 | 142 | 201 | 212 |
| N1.V165(N1).R2.R2 | 130 | 189 | 251 | 258 |
| V121.R2.R2 | 41 | 98 | 167 | 213 |

**Supplementary Table S8. Nonsignaling NRP1-only complexes.** Dots indicate direct binding. This table gives the number by which each molecule or molecular complex is identified in the model code. V165 represents $VEGFA_{165a}$ and P2 represents $PLGF_2$.

| Molecule/Complex | Surface | Rab4a5a | Rab11a | Lysosome (degraded) |
|---|---|---|---|---|
| V165.N1 | 14 | 48 | 105 | 150 |
| P2.N1 | 15 | 49 | 106 | 156 |
| N1.V165.N1 | 56 | 133 | 192 | 208 |
| N1.P2.N1 | 57 | 134 | 193 | 227 |

**Supplementary Table S9. Signaling ligand-bound VEGFR1 dimers.** This table gives the number by which each molecule or molecular complex is identified in the model code. Dots indicate direct binding. A Δ symbol indicates that the ligand is bound to both VEGFR1, and that the two VEGFR1 are also directly associated with each other. V165 represents $VEGFA_{165a}$, V121 represents $VEGFA_{121a}$, P1 represents $PLGF_1$, and P2 represents $PLGF_2$.

| Molecule/Complex | Surface | Rab4a5a | Rab11a | Lysosome (degraded) |
|---|---|---|---|---|
| R1.V165.R1 | 60 | 137 | 196 | 215 |
| R1.V121.R1 | 61 | 138 | 197 | 217 |
| R1.V121.R1.N1 | 67 | 144 | 203 | 221 |
| N1.R1.V121.R1.N1 | 131 | 190 | 252 | 266 |
| R1.V165.R1 Δ | 121 | 181 | 243 | 260 |
| R1.V121.R1 Δ | 122 | 182 | 244 | 262 |
| R1.V121.R1.N1 Δ | 123 | 183 | 245 | 264 |
| N1.R1.V121.R1.N1 Δ | 175 | 237 | 275 | 280 |
| R1.P1.R1 | 62 | 139 | 198 | 222 |
| R1.P1.R1.N1 | 69 | 146 | 205 | 226 |
| N1.R1.P1.R1.N1 | 132 | 191 | 253 | 271 |
| R1.P2.R1 | 63 | 140 | 199 | 228 |
| R1.P1.R1 Δ | 124 | 184 | 246 | 267 |
| R1.P1.R1.N1 Δ | 125 | 185 | 247 | 269 |
| N1.R1.P1.R1.N1 Δ | 176 | 238 | 276 | 281 |
| R1.P2.R1 Δ | 127 | 186 | 248 | 272 |

**Supplementary Table S10. Signaling ligand-bound VEGFR2 dimers.** This table gives the number by which each molecule or molecular complex is identified in the model code. Dots and parentheses indicate direct binding. A ∆ symbol indicates that the ligand is bound to both VEGFR2, and that the two VEGFR2 are also directly associated with each other. V165 represents $VEGFA_{165a}$ and V121 represents $VEGFA_{121a}$ .

| **Molecule/Complex** | **Surface** | **Rab4a5a** | **Rab11a** | **Lysosome (degraded)** |
|---|---|---|---|---|
| R2.V165.R2 | 58 | 135 | 194 | 209 |
| R2.V165(N1).R2 | 129 | 188 | 250 | 255 |
| R2.(N1)V165(N1).R2 | 177 | 239 | 277 | 278 |
| R2.V121.R2 | 59 | 136 | 195 | 214 |
| R2.V165.R2 ∆ | 117 | 178 | 240 | 254 |
| R2.V165(N1).R2 ∆ | 118 | 179 | 241 | 256 |
| R2.(N1)V165(N1).R2 ∆ | 174 | 236 | 274 | 279 |
| R2.V121.R2 ∆ | 119 | 180 | 242 | 259 |

**Supplementary Table S11. Matrix-bound complexes.** Due to the size of matrix molecules, these complexes are only at the surface and are not internalized. However, ligand-coupled receptors can still serve as signal initiation (Wendel Clegg & Mac Gabhann, 2015). This table gives the number by which each molecule or molecular complex is identified in the model code. Dots and parentheses indicate direct binding. A ∆ symbol indicates that the ligand is bound to both VEGFR, and that the two VEGFR are also associated. This table is included for complete description of the code provided, however in the present study matrix binding is not included and all these molecules will have zero concentration. V165 represents $VEGFA_{165a}$, V121 represents $VEGFA_{121a}$, P1 represents $PLGF_1$, and P2 represents $PLGF_2$.

| Molecule/Complex | Surface | Signaling |
|---|---|---|
| M | 8 | |
| M.V165 | 12 | |
| M.V165.R1 | 34 | |
| M.V165.R1.R1 | 37 | |
| M.V165.R1.R1.N1 | 84 | |
| R1.(M)V165.R1 | 114 | Yes |
| R1.(M)V165.R1 ∆ | 120 | Yes |
| M.V165.R2 | 36 | |
| M.V165.R2.R2 | 39 | |
| R2.(M)V165.R2 | 113 | Yes |
| R2.(M)V165.R2 ∆ | 116 | Yes |
| M.P2 | 13 | |
| M.P2.R1 | 35 | |
| M.P2.R1.R1 | 38 | |
| M.P2.R1.R1.N1 | 85 | |
| R1.(M)P2.R1 | 115 | Yes |
| R1.(M)P2.R1 ∆ | 126 | Yes |

**Supplementary Table S12. Model Parameters.** Compartment volumes, compartment surface areas, and initial concentrations of ligands and surface receptors. Ligand molecular weights were obtained from sources of experimental recombinant proteins.

| **Receptor species/ parameters** | **Value** | **Units** | **Reference or assumptions (MW: molecular weight)** |
|---|---|---|---|
| Extracellular volume | $10^{-8}$ | liter/cell | 1 ml cell culture media in one well with $10^5$ cells |
| Rab4a volume | $11.25 \times 10^{-15}$ | liter/cell | (Sarabipour *et al*, 2024) |
| Rab11a volume | $3.75 \times 10^{-15}$ | liter/cell | (Sarabipour *et al*, 2024) |
| Cell surface area | 1,000 | $\mu m^2$/cell | (Jaffe, 1987) |
| Rab4a surface area | 950 | $\mu m^2$/cell | (Sarabipour *et al*, 2024) |
| Rab11a surface area | 350 | $\mu m^2$/cell | (Sarabipour *et al*, 2024) |
| VEGFR1 (cell surface) | 1,800 | receptors per cell | (Imoukhuede & Popel, 2011) |
| VEGFR2 (cell surface) | 4,900 | receptors per cell | (Imoukhuede & Popel, 2011) |
| NRP1 (cell surface) | 68,000 | receptors per cell | (Imoukhuede & Popel, 2011) |
| $VEGFA_{121a}$ | $1.075 \times 10^7$ | molecules/cell | 50 ng.mL$^{-1}$ , MW ~28 kDa |
| $VEGFA_{165a}$ | $6.843 \times 10^6$ | molecules/cell | 50 ng.mL$^{-1}$ , MW ~44 kDa |
| $PLGF_1$ | $1.014 \times 10^7$ | molecules/cell | 50 ng.mL$^{-1}$, MW ~29.7 kDa |
| $PLGF_2$ | $8.702 \times 10^6$ | molecules/cell | 50 ng.mL$^{-1}$, MW ~34.6 kDa |

**Supplementary Table S13. Production rates for VEGF receptors.** These parameters are obtained from optimization in the absence of ligands, and result in steady state surface receptor densities in agreement with previous measurements (Sarabipour *et al*, 2024), when used in concert with the trafficking parameters in Table S14.

| | | | |
|---|---|---|---|
| Production $k_{prod}$ | VEGFR1 | 4.101 | $(\#/cell).s^{-1}$ |
| | VEGFR2 | 1.114 | $(\#/cell).s^{-1}$ |
| | NRP1 | 0.459 | $(\#/cell).s^{-1}$ |

**Supplementary Table S14. Trafficking parameter estimates for VEGFR1, VEGFR2, and NRP1 in the absence and presence of ligands based on experimental data from HUVEC.** This standard parameter set was used for model simulations presented in the main manuscript. Based on modeling and data from HUVECs (Human Umbilical Vein Endothelial Cells). Unligated VEGFR1, VEGFR2, and NRP1 trafficking parameters are from previous work (Sarabipour *et al*, 2024); ligated trafficking parameters are described in this study. R1: VEGFR1, R2: VEGFR2, N1: Neuropilin-1 or NRP1.

| | Receptor | Unliganded **$k$ ($s^{-1}$)** | VEGF-bound **$k$ ($s^{-1}$)** | PLGF-bound **$k$ ($s^{-1}$)** |
|---|---|---|---|---|
| Internalization $k_{int}$ | VEGFR1 | $1.3 \times 10^{-2}$ | $1.3 \times 10^{-2}$ | $1.3 \times 10^{-2}$ |
| | VEGFR2 | $2.3 \times 10^{-4}$ | $3 \times 2.3 \times 10^{-4}$ | |
| | NRP1 | $2.7 \times 10^{-4}$ | $2.7 \times 10^{-4}$ | $2.7 \times 10^{-4}$ |
| | R1.N1 | $1.3 \times 10^{-2}$ | $1.3 \times 10^{-2}$ | $1.3 \times 10^{-2}$ |
| | R2.N1 | | $3 \times 2.3 \times 10^{-4}$ | |
| Recycling to surface via Rab4a endosomes $k_{rec4}$ | VEGFR1 | $5.4 \times 10^{-4}$ | $5.4 \times 10^{-4}$ | $5.4 \times 10^{-4}$ |
| | VEGFR2 | $1.2 \times 10^{-6}$ | $1.2 \times 10^{-6}$ | |
| | NRP1 | $2.1 \times 10^{-2}$ | $2.1 \times 10^{-2}$ | $2.1 \times 10^{-2}$ |
| | R1.N1 | $5.4 \times 10^{-4}$ | $5.4 \times 10^{-4}$ | $5.4 \times 10^{-4}$ |
| | R2.N1 | | $1.2 \times 10^{-6}$ | |
| Transfer from Rab4a endosomes to Rab11a endosomes $k_{4to11}$ | VEGFR1 | $5.9 \times 10^{-4}$ | $5.9 \times 10^{-4}$ | $5.9 \times 10^{-4}$ |
| | VEGFR2 | $1.5 \times 10^{-6}$ | $1.5 \times 10^{-6}$ | |
| | NRP1 | $7.0 \times 10^{-2}$ | $7.0 \times 10^{-2}$ | $7.0 \times 10^{-2}$ |
| | R1.N1 | $5.9 \times 10^{-4}$ | $5.9 \times 10^{-4}$ | $5.9 \times 10^{-4}$ |
| | R2.N1 | | $1.5 \times 10^{-6}$ | |
| Recycling to surface via Rab11a endosomes $k_{rec11}$ | VEGFR1 | $1.0 \times 10^{-1}$ | $1.0 \times 10^{-1}$ | $1.0 \times 10^{-1}$ |
| | VEGFR2 | $8.9 \times 10^{-2}$ | $8.9 \times 10^{-2}$ | |
| | NRP1 | $7.9 \times 10^{-4}$ | $7.9 \times 10^{-4}$ | $7.9 \times 10^{-4}$ |
| | R1.N1 | $1.0 \times 10^{-1}$ | $1.0 \times 10^{-1}$ | $1.0 \times 10^{-1}$ |
| | R2.N1 | | $8.9 \times 10^{-2}$ | |
| Degradation $k_{deg}$ | VEGFR1 | $2.3 \times 10^{-4}$ | $2.3 \times 10^{-4}$ | $2.3 \times 10^{-4}$ |
| | VEGFR2 | $2.3 \times 10^{-4}$ | $2.3 \times 10^{-4}$ | |
| | NRP1 | $1.2 \times 10^{-6}$ | $1.2 \times 10^{-6}$ | $1.2 \times 10^{-6}$ |
| | R1.N1 | $2.3 \times 10^{-4}$ | $2.3 \times 10^{-4}$ | $2.3 \times 10^{-4}$ |
| | R2.N1 | | $2.3 \times 10^{-4}$ | |

**Supplementary Table S15. Receptor dimerization parameters.** R1: VEGFR1, R2: VEGFR2, N1: Neuropilin-1/NRP1. The unligated receptor dimerization rates were set in a previous study to yield 30-40% dimers of R1-R1, R2-R2, and N1-R1 (Sarabipour *et al*, 2024). Note that these base parameters (in units of molecules$^{-1}$.µm$^2$.s$^{-1}$) are adjusted to units of molecules$^{-1}$.cell.s$^{-1}$ at each location, using the appropriate membrane surface area (Table S12), as described previously (Sarabipour *et al*, 2024).

| | **Description** | **$k_{on}$ (molecules$^{-1}$.µm$^2$.s$^{-1}$)** | **$k_{off}$ (s$^{-1}$)** | **$K_D$ (molecules.µm$^{-2}$)** | **Reference** |
|---|---|---|---|---|---|
| R1-R1 | unligated VEGFR1 dimerization | $8.0 \times 10^{-4}$ | $1.0 \times 10^{-2}$ | 12.5 | See (Sarabipour *et al*, 2024) |
| R2-R2 | unligated VEGFR2 dimerization | $2.0 \times 10^{-3}$ | $1.0 \times 10^{-2}$ | 5 | (Sarabipour *et al*, 2016; da Rocha-Azevedo *et al*, 2020) & see (Sarabipour *et al*, 2024) |
| N1-R1 | unligated NRP1-VEGFR1 dimerization | $8.0 \times 10^{-4}$ | $1.0 \times 10^{-2}$ | 12.5 | (Wendel Clegg & Mac Gabhann, 2015; Wu *et al*, 2009; Fuh *et al*, 2000) & see (Sarabipour *et al*, 2024) |
| | **Description** | **$k_{on}$ Surface (molecules$^{-1}$.cell.s$^{-1}$)** | **$k_{on}$ Rab4a (molecules$^{-1}$.cell.s$^{-1}$)** | **$k_{on}$ Rab11a (molecules$^{-1}$.cell.s$^{-1}$)** | **Reference** |
| R1-R1 | unligated VEGFR1 dimerization | $8.0 \times 10^{-7}$ | $8.42 \times 10^{-7}$ | $2.46 \times 10^{-6}$ | See (Sarabipour *et al*, 2024) |
| R2-R2 | unligated VEGFR2 dimerization | $2.0 \times 10^{-6}$ | $2.11 \times 10^{-6}$ | $6.15 \times 10^{-6}$ | See (Sarabipour *et al*, 2024) |
| N1-R1 | unligated NRP1-VEGFR1 dimerization | $8.0 \times 10^{-7}$ | $8.42 \times 10^{-7}$ | $2.46 \times 10^{-6}$ | See (Sarabipour *et al*, 2024) |

**Supplementary Table S16. Experimentally-derived 1:1 Ligand-Receptor binding.** Measured and estimated rate constants and equilibrium constants assuming 1:1 interaction (i.e. monovalent ligand binds monovalent receptor dimer fully in one step; this is the most common assumption for estimating ligand-receptor binding experimentally). These are not the values used in our model, which is a dimerization-explicit model, but the parameters used (Table S17) are based on these as described in the Methods section. In each entry the units of the parameters shown are (top to bottom): $K_D$ (pM), $k_{on}$ ($pM^{-1}$ $s^{-1}$), and $k_{off}$ ($s^{-1}$). L: Ligand; R1: VEGFR1; R2: VEGFR2; N1: NRP1.

| Interaction | Parameter | $VEGFA_{121a}$ | $VEGFA_{165a}$ | $PLGF_1$ | $PLGF_2$ | Reference |
|---|---|---|---|---|---|---|
| L-R1 | $K_d$<br>$k_{on}$<br>$k_{off}$ | 33<br>$3 \times 10^{-5}$<br>$10^{-3}$ | 33<br>$3 \times 10^{-5}$<br>$10^{-3}$ | 233<br>$1.5 \times 10^{-6}$<br>$3.5 \times 10^{-4}$ | 233<br>$1.5 \times 10^{-6}$<br>$3.5 \times 10^{-4}$ | (Wendel Clegg & Mac Gabhann, 2015; Wu *et al*, 2009) |
| L-R2 | $K_d$<br>$k_{on}$<br>$k_{off}$ | 100<br>$1 \times 10^{-5}$<br>$10^{-3}$ | 100<br>$1 \times 10^{-5}$<br>$10^{-3}$ | | | (Wu *et al*, 2009; Mac Gabhann & Popel, 2004; Wendel Clegg & Mac Gabhann, 2017) |
| L-N1 | $K_d$<br>$k_{on}$<br>$k_{off}$ | | 1,200<br>$5 \times 10^{-7}$<br>$6 \times 10^{-4}$ | | 100,000<br>$1 \times 10^{-8}$<br>$1 \times 10^{-3}$ | (Wendel Clegg & Mac Gabhann, 2017; Vintonenko *et al*, 2011; Hoffmann *et al*, 2013) |
| L-(N1R1) | $K_d$<br>$k_{on}$<br>$k_{off}$ | 33<br>$3 \times 10^{-5}$<br>$1 \times 10^{-3}$ | | 233<br>$1.5 \times 10^{-6}$<br>$3.5 \times 10^{-4}$ | | (Wendel Clegg & Mac Gabhann, 2017; Vintonenko *et al*, 2011; Hoffmann *et al*, 2013) |

**Supplementary Table S17. Ligand-Receptor binding.** Calculated rate constants for first binding step to monomers or dimers assuming 1:2 interaction (i.e. one bivalent ligand can bind to two monovalent receptor monomers, or twice to one bivalent receptor dimer). The base on-rate constants ($k_{on}$) are assumed to be one quarter of the 1:1 rate constant (Table S16) to account for the bivalency of ligands and of receptor dimers. Thus, when considering these base on-rate constant values below, ligand binding to a receptor monomer is twice the rate constant (due to two receptor-binding sites on the ligand) and ligand binding to an unliganded receptor dimer is four times the rate constant (due to two receptor-binding sites on the ligands and two ligand-binding sites on the receptor dimer). L: Ligand; R1: VEGFR1; R2: VEGFR2; N1: NRP1.

| Interaction | Rate Constant | $VEGFA_{121a}$ | $VEGFA_{165a}$ | $PLGF_1$ | $PLGF_2$ |
|---|---|---|---|---|---|
| L-R1 | $k_{on}$ (pM$^{-1}$ s$^{-1}$) | 2* or 4*<br>$7.5 \times 10^{-6}$ | 2* or 4*<br>$7.5 \times 10^{-6}$ | 2* or 4*<br>$3.75 \times 10^{-7}$ | 2* or 4*<br>$3.75 \times 10^{-7}$ |
| | $k_{on}$ (#/cell)$^{-1}$s$^{-1}$ | 2* or 4* | 2* or 4* | 2* or 4* | 2* or 4* |
| | *surface* | $1.25 \times 10^{-9}$ | $1.25 \times 10^{-9}$ | $6.23 \times 10^{-11}$ | $6.23 \times 10^{-11}$ |
| | *rab4a* | $1.11 \times 10^{-3}$ | $1.11 \times 10^{-3}$ | $5.54 \times 10^{-5}$ | $5.54 \times 10^{-5}$ |
| | *rab11a* | $3.32 \times 10^{-3}$ | $3.32 \times 10^{-3}$ | $1.66 \times 10^{-4}$ | $1.66 \times 10^{-4}$ |
| | $k_{off}$ (s$^{-1}$) | $2.24 \times 10^{-2}$ | $2.24 \times 10^{-2}$ | $1.32 \times 10^{-2}$ | $1.32 \times 10^{-2}$ |
| L-R2 | $k_{on}$ (pM$^{-1}$ s$^{-1}$) | 2* or 4*<br>$2.5 \times 10^{-6}$ | 2* or 4*<br>$2.5 \times 10^{-6}$ | | |
| | $k_{on}$ (#/cell)$^{-1}$s$^{-1}$ | 2* or 4* | 2* or 4* | | |
| | *surface* | $4.15 \times 10^{-10}$ | $4.15 \times 10^{-10}$ | | |
| | *rab4a* | $3.69 \times 10^{-4}$ | $3.69 \times 10^{-4}$ | | |
| | *rab11a* | $1.11 \times 10^{-3}$ | $1.11 \times 10^{-3}$ | | |
| | $k_{off}$ (s$^{-1}$) | $2.24 \times 10^{-2}$ | $2.24 \times 10^{-2}$ | | |
| L-N1 | $k_{on}$ (pM$^{-1}$ s$^{-1}$) | | 2*<br>$1.25 \times 10^{-7}$ | | 2*<br>$2.5 \times 10^{-9}$ |
| | $k_{on}$ (#/cell)$^{-1}$s$^{-1}$ | | 2* | | 2* |
| | *surface* | | $2.08 \times 10^{-11}$ | | $4.15 \times 10^{-13}$ |
| | *rab4a* | | $1.85 \times 10^{-5}$ | | $3.69 \times 10^{-7}$ |
| | *rab11a* | | $5.54 \times 10^{-5}$ | | $1.11 \times 10^{-6}$ |
| | $k_{off}$ (s$^{-1}$) | | $1.73 \times 10^{-2}$ | | $2.24 \times 10^{-2}$ |
| L-(N1R1) | $k_{on}$ (pM$^{-1}$ s$^{-1}$) | 2* or 4*<br>$7.5 \times 10^{-6}$ | | 2* or 4*<br>$3.75 \times 10^{-7}$ | |
| | $k_{on}$ (#/cell)$^{-1}$s$^{-1}$ | 2* or 4* | | 2* or 4* | |
| | *surface* | $1.25 \times 10^{-9}$ | | $6.23 \times 10^{-11}$ | |
| | *rab4a* | $1.11 \times 10^{-3}$ | | $5.54 \times 10^{-5}$ | |
| | *rab11a* | $3.32 \times 10^{-3}$ | | $1.66 \times 10^{-4}$ | |
| | $k_{off}$ (s$^{-1}$) | $2.24 \times 10^{-2}$ | | $1.32 \times 10^{-2}$ | |

**Supplementary Table S18. Ligand-Receptor Coupling.** These are the values for intermolecular and intramolecular binding of ligands and receptors following the first ligand-receptor binding event. Values for receptor-receptor coupling without a ligand ($k_{c,RR}$) are given in Supplementary Table S15. The binding rate constant for a ligand (already bound to a receptor monomer) binding to a second receptor monomer ($k_{c,LR}$); the binding rate constant for a ligand (already bound to one receptor in a dimer) binding to the second receptor in a dimer ($k_{\Delta,LR}$); and the binding rate constant for a receptor binding to the second receptor of a dimer (when both are already bound to a ligand) ($k_{\Delta,RR}$) are calculated based on other known parameters as described in *Methods* and in a previous publication (Mac Gabhann & Popel, 2007). L: Ligand; R1: VEGFR1; R2: VEGFR2; N1: NRP1.

| Interaction | Rate Constant | $VEGFA_{121a}$ | $VEGFA_{165a}$ | $PLGF_1$ | $PLGF_2$ |
|---|---|---|---|---|---|
| L-R1 | $k_{c,LR}$ (#/cell)$^{-1}$s$^{-1}$<br>*surface*<br>*rab4a*<br>*rab11a*<br>$k_{c,RR}/k_{c,LR}$<br>$k_{\Delta,LR}$ (s$^{-1}$)<br>$k_{\Delta,RR}$ (s$^{-1}$) | <br>5.31 x 10$^{-4}$<br>5.59 x 10$^{-4}$<br>1.63 x 10$^{-3}$<br>1.51 x 10$^{-3}$<br>9.55 x 10$^{-1}$<br>1.44 x 10$^{-3}$ | <br>5.31 x 10$^{-4}$<br>5.59 x 10$^{-4}$<br>1.63 x 10$^{-3}$<br>1.51 x 10$^{-3}$<br>9.55 x 10$^{-1}$<br>1.44 x 10$^{-3}$ | <br>5.41 x 10$^{-4}$<br>5.69 x 10$^{-4}$<br>1.66 x 10$^{-3}$<br>1.48 x 10$^{-3}$<br>9.74 x 10$^{-1}$<br>1.44 x 10$^{-3}$ | <br>5.41 x 10$^{-4}$<br>5.69 x 10$^{-4}$<br>1.66 x 10$^{-3}$<br>1.48 x 10$^{-3}$<br>9.74 x 10$^{-1}$<br>1.44 x 10$^{-3}$ |
| L-R2 | $k_{c,LR}$ (#/cell)$^{-1}$s$^{-1}$<br>*surface*<br>*rab4a*<br>*rab11a*<br>$k_{c,RR}/k_{c,LR}$<br>$k_{\Delta,LR}$ (s$^{-1}$)<br>$k_{\Delta,RR}$ (s$^{-1}$) | <br>1.91 x 10$^{-4}$<br>2.01 x 10$^{-4}$<br>5.88 x 10$^{-4}$<br>1.05 x 10$^{-2}$<br>9.55 x 10$^{-1}$<br>1.00 x 10$^{-2}$ | <br>1.91 x 10$^{-4}$<br>2.01 x 10$^{-4}$<br>5.88 x 10$^{-4}$<br>1.05 x 10$^{-2}$<br>9.55 x 10$^{-1}$<br>1.00 x 10$^{-2}$ | | |
| L-N1 | $k_{c,LR}$ (#/cell)$^{-1}$s$^{-1}$<br>*surface*<br>*rab4a*<br>*rab11a* | | <br>1.42 x 10$^{-5}$<br>1.49 x 10$^{-5}$<br>4.37 x 10$^{-5}$ | | <br>1.41 x 10$^{-5}$<br>1.48 x 10$^{-5}$<br>4.32 x 10$^{-5}$ |
| L-(N1R1) | $k_{c,LR}$ (#/cell)$^{-1}$s$^{-1}$<br>*surface*<br>*rab4a*<br>*rab11a*<br>$k_{c,RR}/k_{c,LR}$<br>$k_{\Delta,LR}$ (s$^{-1}$)<br>$k_{\Delta,RR}$ (s$^{-1}$) | <br>5.31 x 10$^{-4}$<br>5.59 x 10$^{-4}$<br>1.63 x 10$^{-3}$<br>1.51 x 10$^{-3}$<br>9.55 x 10$^{-1}$<br>1.44 x 10$^{-3}$ | | <br>5.41 x 10$^{-4}$<br>5.69 x 10$^{-4}$<br>1.66 x 10$^{-3}$<br>1.48 x 10$^{-3}$<br>9.74 x 10$^{-1}$<br>1.44 x 10$^{-3}$ | |
| R2-L-N1 | $k_{c,RLN}$ (#/cell)$^{-1}$s$^{-1}$<br>*surface*<br>*rab4a*<br>*rab11a* | | <br>2.50 x 10$^{-6}$<br>2.63 x 10$^{-6}$<br>7.69 x 10$^{-6}$ | | |

**Supplementary Table S19. List of experimental reagents and antibodies.**

| **Reagents** | **Company (catalog #)** | |
|---|---|---|
| HUVECs | Lonza (#2519A)<br>Lot #s: 0000704189 and 0000661173 | |
| HUVEC culture media and supplements | EBM-2 medium supplemented with the bullet kit (EGM-2) (Lonza) | |
| siRNA Rab4a oligonucleotide | ThermoFisher Scientific 439084 (s11675) | |
| siRNA Rab11a oligonucleotide | ThermoFisher Scientific 4390824 (s16702)<br>or Santa Cruz Biotechnology (sc3630) | |
| siRNA transfection reagent Lipofectamine™ 3000 | Thermo Fisher Scientific (L3000001) | |
| Biotinylation kit | Pierce™ Cell Surface Biotinylation<br>and Protein Isolation Kit (#A44390) | |
| Recombinant Human $PLGF_1$ | R&D (264-PGB-010) | |
| $VEGF_{165a}$ | Genscript (Z02689) | |
| Antibody | Company | IB titer |
| VEGFR1 (membrane-integral) | CST (#2893) | 1:1000 |
| VEGFR2 (membrane-integral) | CST (#2479) | 1:10000 |
| NRP1 (membrane-integral) | R&D (AF3870) | 1:1000 |
| PECAM1 | CST (#3528) | 1:100000 |
| α-Tubulin | CST (#3873) | 1:100000 |
| β-Actin | CST (#3700) | 1:10000 |
| Rab4a | ThermoFisher (MA5-17161) | 1:1000 |
| Rab11a | Abcam (ab65200), BD Bio (610656) | 1:2000 |

| | | |
|---|---|---|
| Anti-mouse IgG, HRP-linked (secondary) | ThermoFisher (A16011) | 1:10000 |
| Anti-rabbit IgG, HRP-linked (secondary) | ThermoFisher (A16035) | 1:10000 |
| **Abbreviations**<br>**IB:** Immunoblot, **CST:** Cell Signaling Technologies<br>**HUVECs:** Human Umbilical Vein Endothelial Cells<br>**VEGFR:** Vascular Endothelial Growth Factor Receptor; **NRP:** Neuropilin<br>**PECAM:** Platelet Endothelial Cell Adhesion Molecule, also known as CD31 | | |

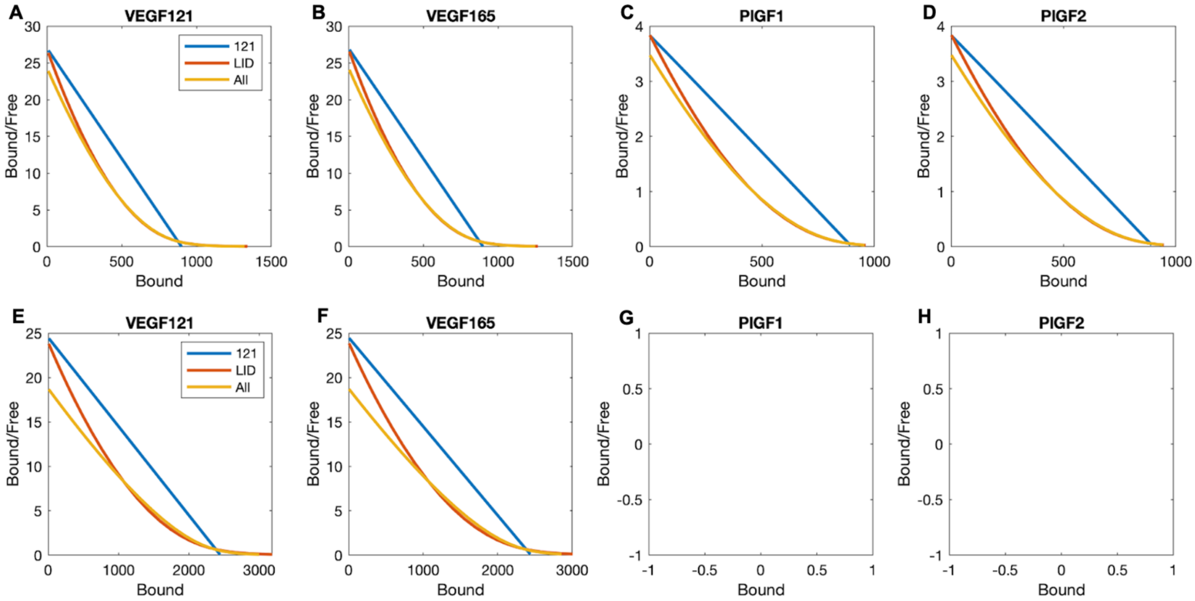

**Figure S1. Simulated Scatchard plots comparing different representations of VEGFR dimerization.** The system was simulated under three different assumptions: one-to-one ("121") ligand receptor binding, i.e. pre-dimerization of receptors and single step ligand binding/activation; ligand-induced dimerization ("LID"), i.e. no receptor pre-dimerization; and a full dimerization model including all dimerization paths ("All"). The similarity between the "121" and "All" lines indicates that the dimerization model represents the observed equilibrium data well. Top row: VEGFR1 expression only; bottom row: VEGFR2 expression only.

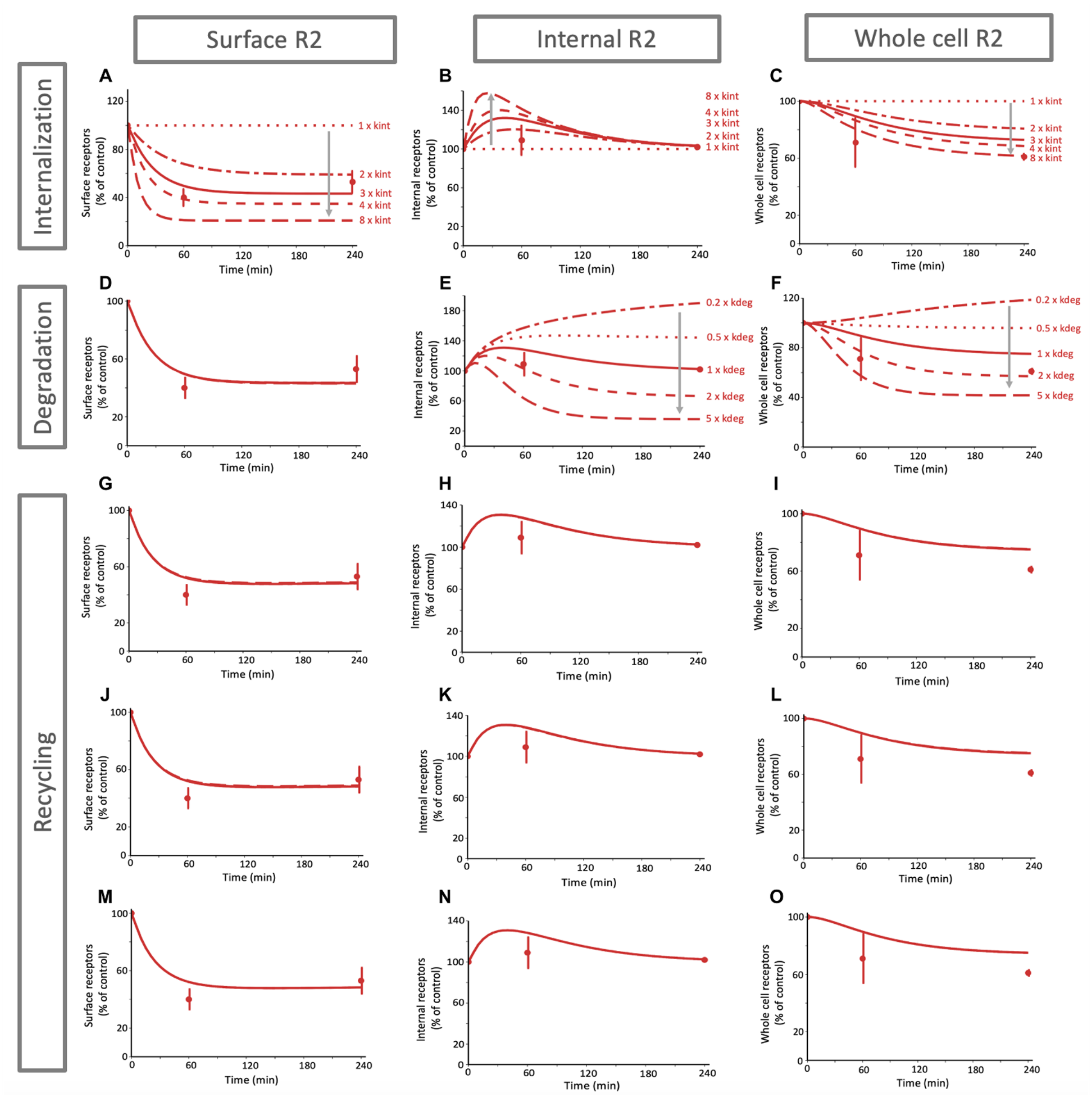

**Figure S2. Distribution of VEGFR2 over 4 hours of VEGF$_{165a}$ treatment.** Total (ligated and unligated) receptor levels on the cell surface, inside the cell, and across the whole cell in response to VEGF$_{165a}$ treatment at different levels of how the ligand binding affects the indicated VEGFR2 trafficking parameter. Simulations are shown as solid and dotted lines. Gray arrows indicate the direction of increasing parameter values. The same experimental data is shown in each row as dots and variance bars. **A-C,** VEGF$_{165a}$ binding causes an increase in the internalization rate constant for VEGFR2 (V.R2.kint and V.R2.N1.kint), compared to the unligated value; a three-fold increase (solid line) compared to the unligated receptor rate constant matched the observed data best. **D-O**, Taking this increased $k_{int}$ (receptor complex internalization rate) as a baseline for the remaining simulations, we explored variation in the other trafficking parameters. Lines represent 5x, 2x, 1x, 0.5x, 0.2x the baseline (unligated) trafficking rate; in panels D and G-O the lines overlap substantially. **D-F,** total surface, internal and whole cell receptors in response to VEGF$_{165a}$ treatment and change in ligated VEGFR2 degradation (altered $k_{deg}$ affecting both V.R2 and V.R2.N1 complexes). **G-O,** total surface, internal and whole cell receptors in response to $VEGF_{165a}$ treatment and change in ligated VEGFR2 recycling, including: recycling via Rab4a (altered $k_{rec4}$, G-I); transfer to Rab11a (altered $k_{4to11}$, J-L); and recycling via Rab11a (altered $k_{rec11}$, M-O). Altered rate constants affect both V.R2 and V.R2.N1 complexes. Panels A-F are identical to panels A-F of Figure 3; they are replicated here for ease of comparison. Gray arrows indicate the direction of increasing parameter values.

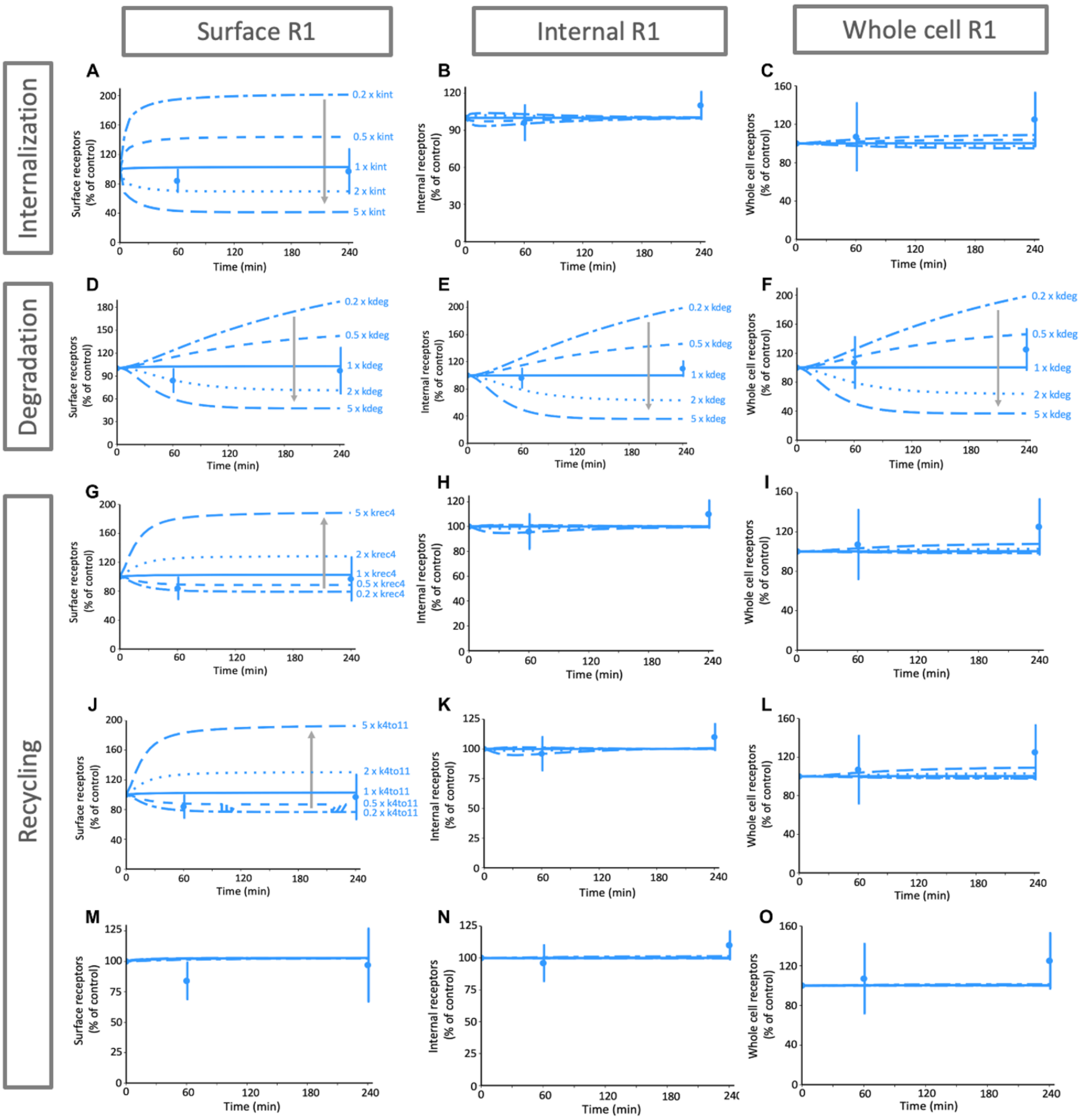

**Figure S3. Distribution of VEGFR1 over 4 hours of $VEGF_{165a}$ treatment.** Total (ligated and unligated) receptor levels on the cell surface, inside the cell, and across the whole cell in response to $VEGF_{165a}$ treatment at different levels of how the ligand binding affects the indicated VEGFR1 trafficking parameter. Simulations are shown as solid and dotted lines. Lines represent 5x, 2x, 1x, 0.5x, 0.2x the baseline (unligated) trafficking rate. Gray arrows indicate the direction of increasing parameter values. The same experimental data is shown in each row as dots and variance bars. **A-C,** Response to $VEGF_{165a}$ treatment for different values of the ligated VEGFR1 internalization parameter ($k_{int}$). . **D-F,** Response to $VEGF_{165a}$ treatment for different values of the ligated VEGFR1 degradation parameter ($k_{deg}$), **G-I,** Tesponse to $VEGF_{165a}$ treatment for different values of the ligated VEGFR1 fast recycling via Rab4a-expressing endosomes parameter ($k_{rec4}$), **J-L,** Response to $VEGF_{165a}$ treatment for different ligated VEGFR1 transfer from Rab4a-expressing endosomes to Rab11a-expressing endosomes parameter ($k_{4to11}$), **M-O,** Response to $VEGF_{165a}$ treatment for different values of the ligated VEGFR1 slow recycling via Rab11a-expressing endosomes parameter ($k_{rec11}$).

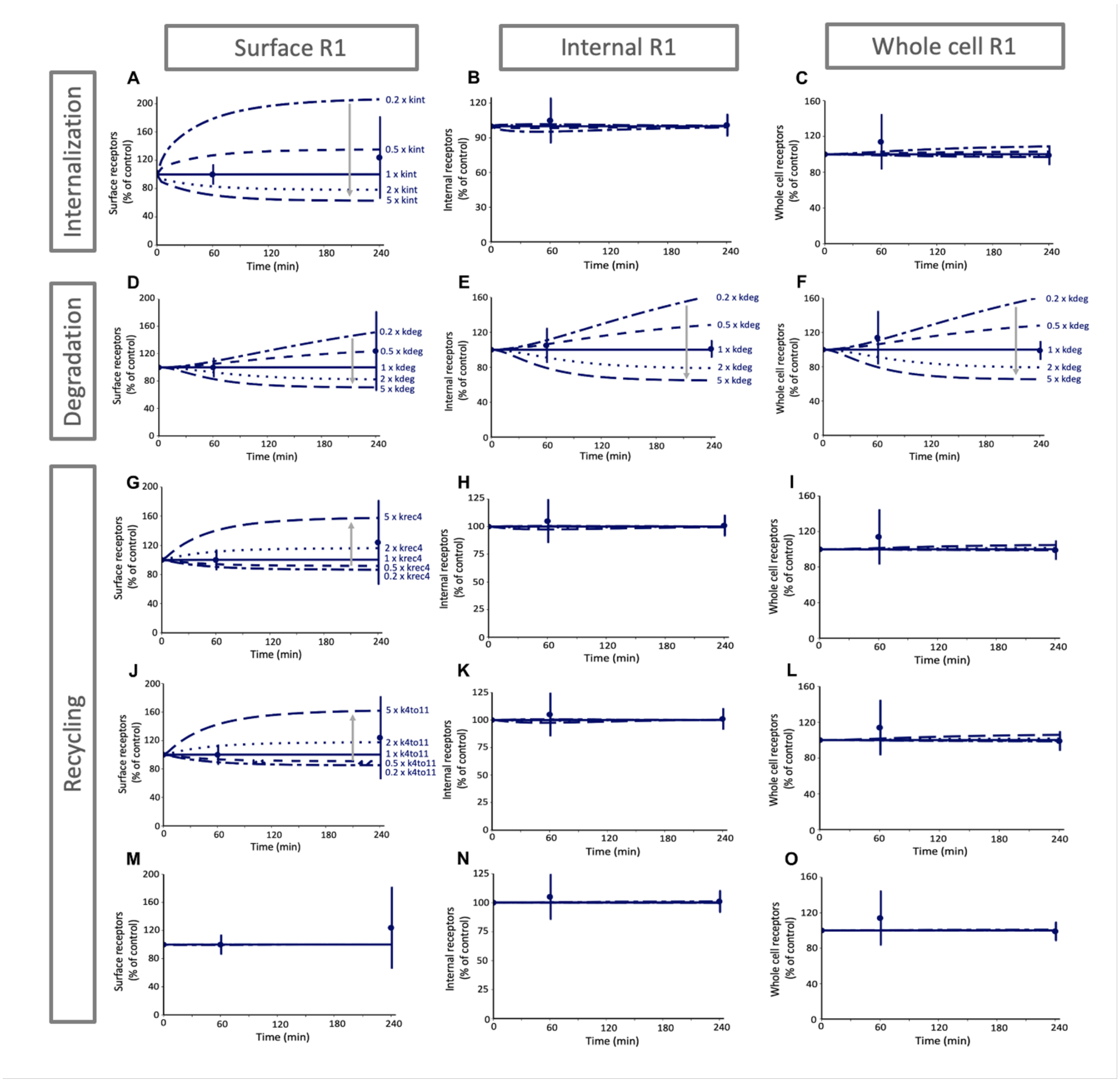

**Figure S4. Distribution of VEGFR1 over 4 hours of PLGF$_1$ treatment.** Total (ligated and unligated) receptor levels on the cell surface, inside the cell, and across the whole cell in response to PLGF$_1$ treatment at different levels of how the ligand binding affects the indicated VEGFR1 trafficking parameter. Simulations are shown as solid and dotted lines. Lines represent 5x, 2x, 1x, 0.5x, 0.2x the baseline (unligated) trafficking rate. Gray arrows indicate the direction of increasing parameter values. The same experimental data is shown in each row as dots and variance bars. **A-C,** Response to PLGF1 treatment for different values of the ligated VEGFR1 internalization parameter ($k_{int}$, applies to both P.R1 and P.R1.N1 complexes), **D-F,** Response to PLGF$_1$ treatment for different values of the ligated VEGFR1 degradation parameter ($k_{deg}$), **G-I,** Response to PLGF$_1$ treatment for different values of the ligated VEGFR1 fast recycling via Rab4a-expressing endosomes parameter ($k_{rec4}$), **J-L,** Response to PLGF$_1$ treatment for different values of the ligated VEGFR1 transfer from Rab4a-expressing endosomes to Rab11a-expressing endosomes parameter ($k_{4to11}$), **M-O,** Response to PLGF$_1$ treatment for different values of the ligated VEGFR1 slow recycling via Rab11a-expressing endosomes parameter ($k_{rec11}$).

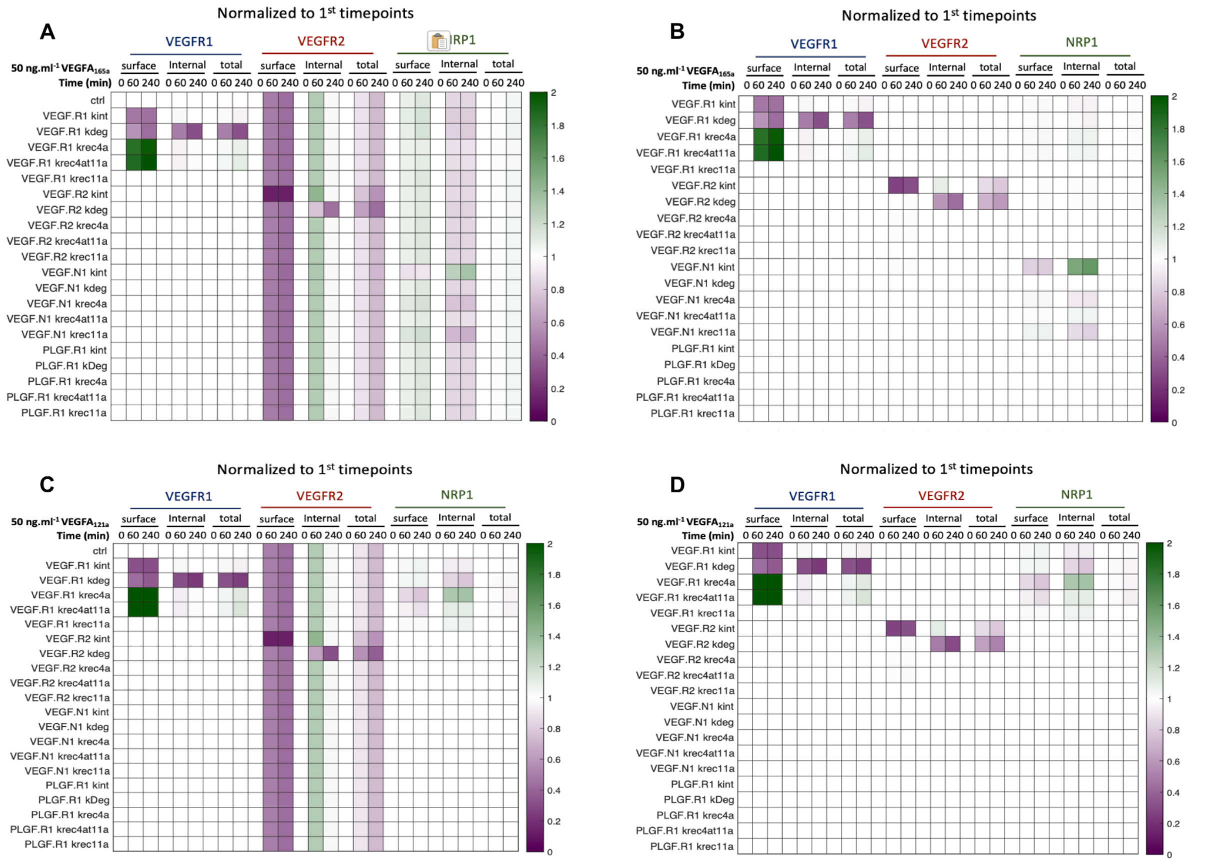

**Figure S5. Summary of impact of trafficking parameters on localization of VEGF receptors following VEGF ligation.** Predicted levels of VEGFR1, VEGFR2, and NRP1 at the surface, internally, and across the whole cell ("total") after 0, 60, or 240 mins treatment with 50 ng.mL$^{-1}$ of VEGF$_{165a}$ (A-B), or VEGF$_{121a}$ (C-D). Each row represents simulations with a different ligated receptor trafficking parameter increased five-fold. Changes in receptor levels normalized to the no-ligand condition (0 mins) are shown (A,C). These are further normalized to the no-parameter-change ("ctrl") condition (B,D), demonstrating the small number of parameters that alter receptor localization.

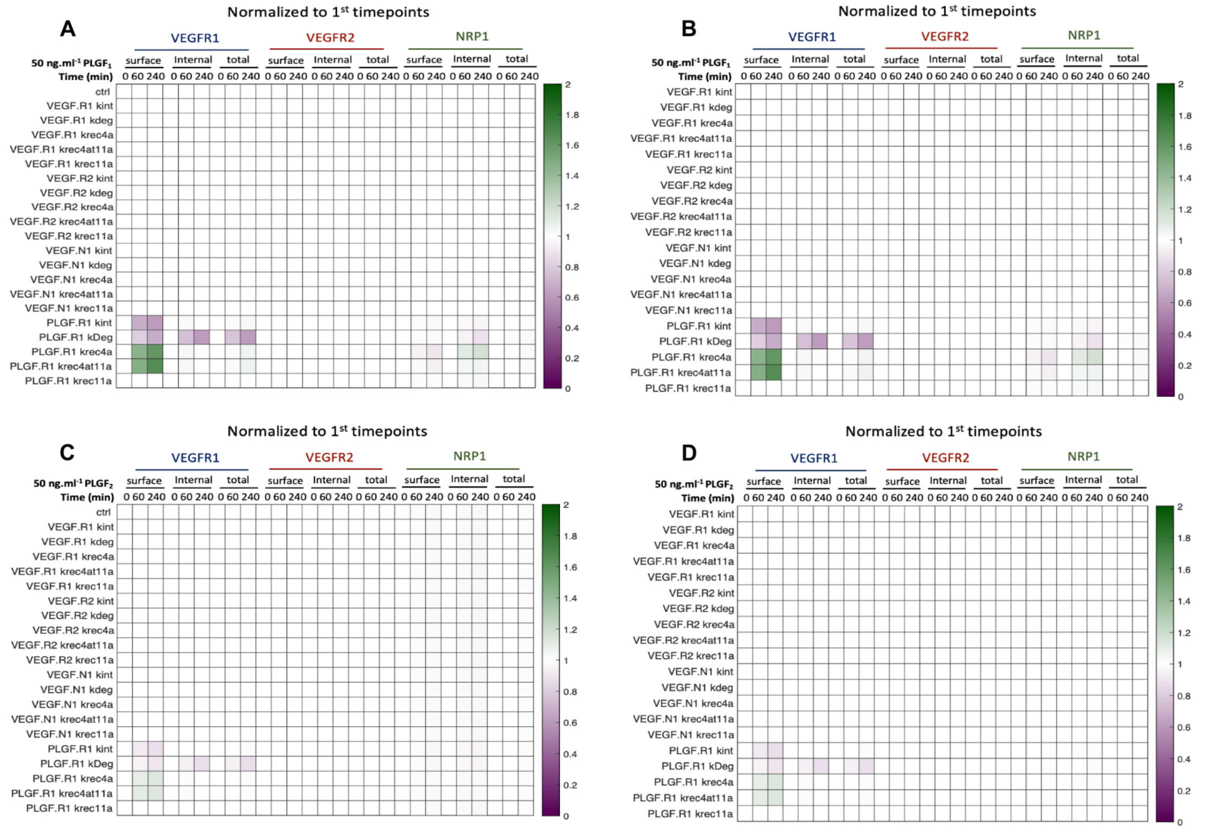

**Figure S6. Summary of impact of trafficking parameters on localization of VEGF receptors following PLGF ligation.** Predicted levels of VEGFR1, VEGFR2, and NRP1 at the surface, internally, and across the whole cell ("total") after 0, 60, or 240 mins treatment with 50 ng.mL$^{-1}$ of PLGF$_1$ (A-B) or PLGF$_2$ (C-D). Each row represents simulations with a different ligated receptor trafficking parameter increased five-fold. Changes in receptor levels normalized to the no-ligand condition (0 mins) are shown (A,C). These are further normalized to the no-parameter-change ("ctrl") condition (B,D), demonstrating the small number of parameters that alter receptor localization.

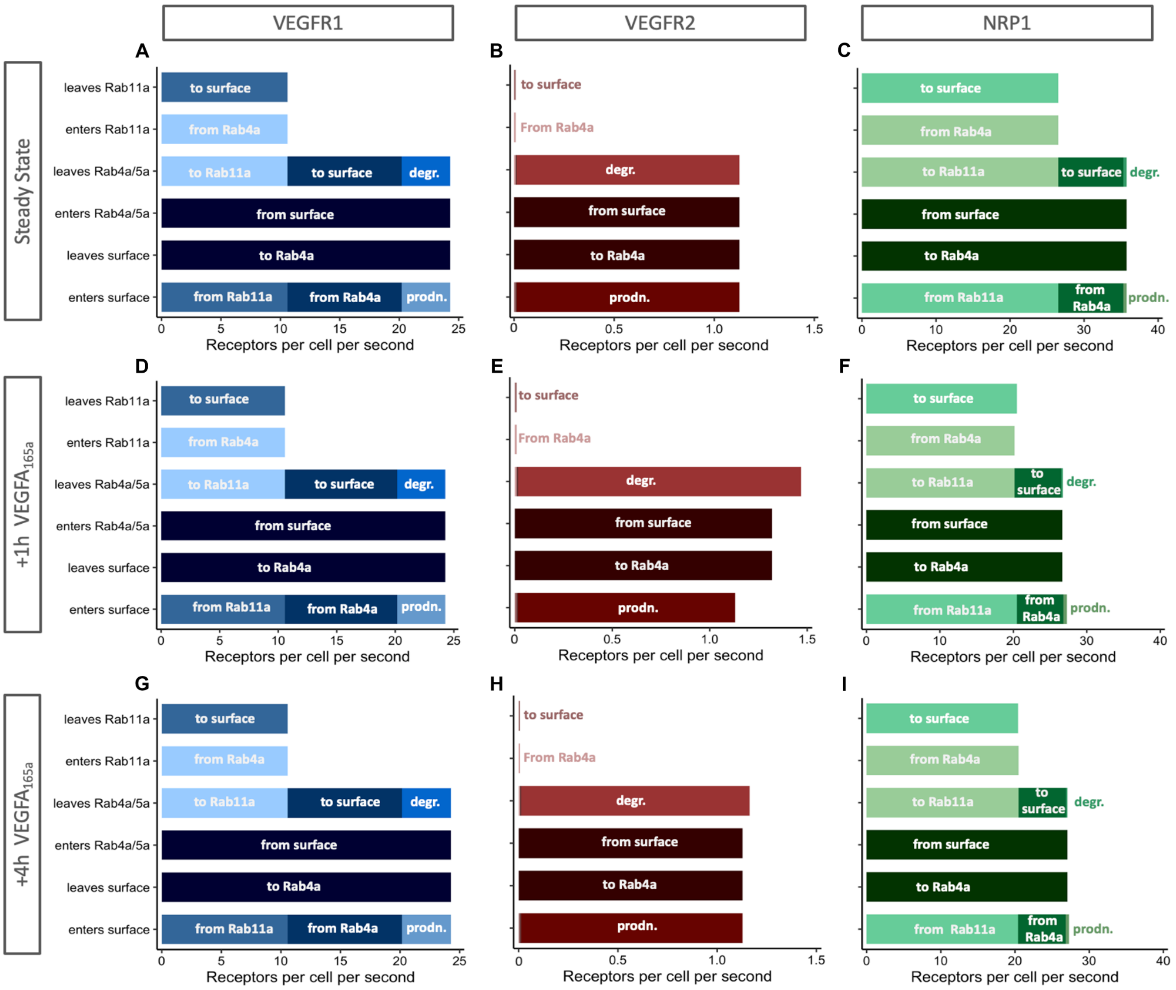

**Figure S7. Differential receptor fluxes.** The overall transport rates (rate constant multiplied by concentration) for VEGFR1, VEGFR2, and NRP1 in each subcellular location at steady state (no ligand treatment) and under different durations of ligand treatment. A rate constant may be high, but if the corresponding concentration is low (and we know that the receptors are not uniformly distributed across cellular compartments), then the rate of movement will be low. Note also that the x-axis scale is different for each receptor. At steady state, these overall rates in and out are balanced, so the 'net' rates are close to zero. For Rab11a: receptors arriving from Rab4a are balanced out by recycling; for Rab4a: receptors internalized to Rab4a are balanced out the sum of degraded and recycling; for surface: receptor internalization balances the sum of new synthesis and recycling. "degr." = degradation; "prodn." = production. **A-C,** transport rates for VEGFR1, VEGFR2, and NRP1 at steady state starvation conditions (no ligand treatment), **D-F,** transport rates for VEGFR1, VEGFR2, and NRP1 after 1 hour of 50 ng.mL$^{-1}$ VEGF$_{165a}$ treatment, **G-I,** transport rates for VEGFR1, VEGFR2, and NRP1 after 4 hours of 50 ng.mL$^{-1}$ VEGF$_{165a}$ treatment.

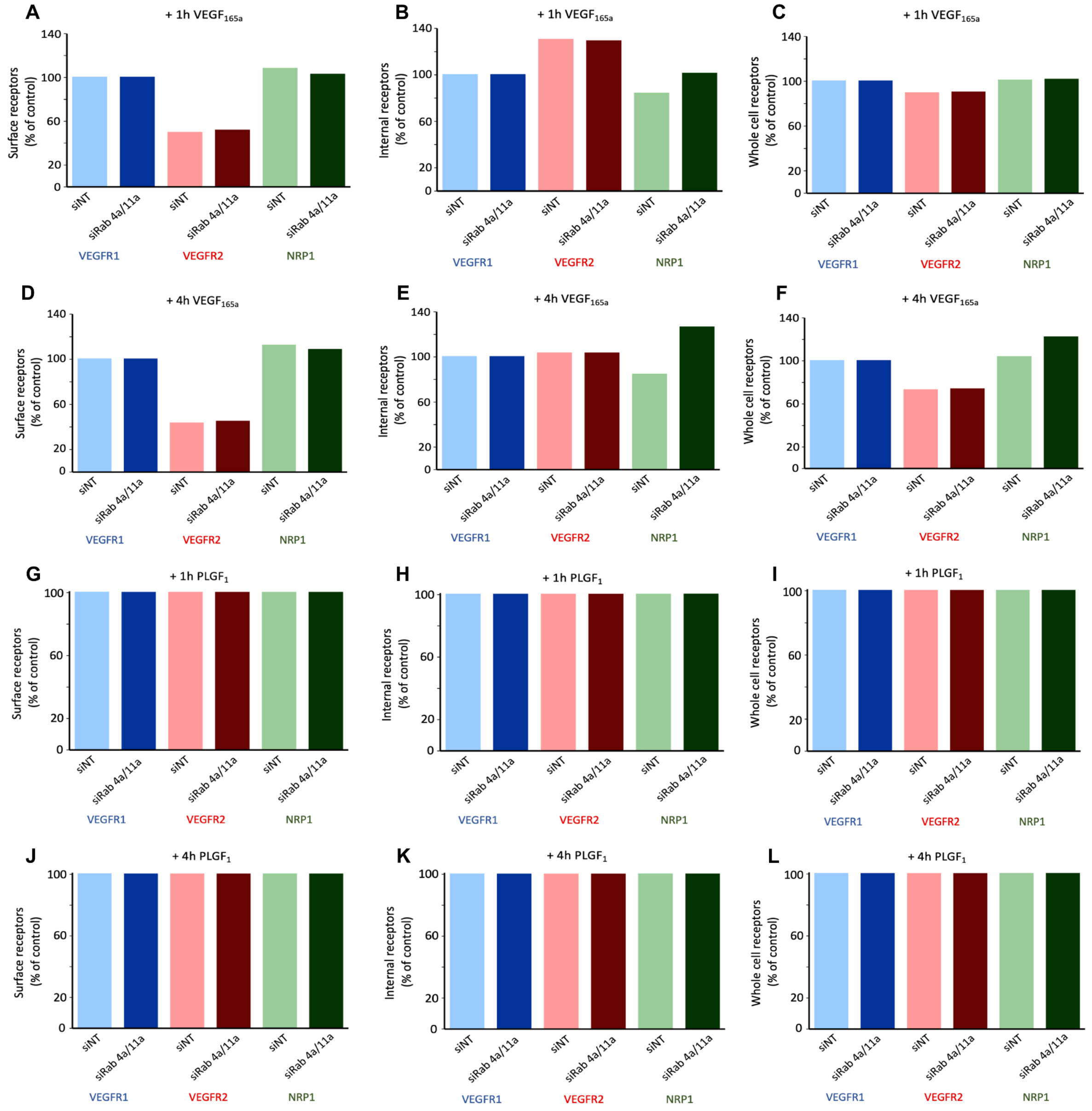

**Figure S8.** Surface, internal and whole cell levels of total (unligated and ligated) VEGFR1, VEGFR2, and NRP1 after Rab4a/Rab11a knockdown, compared to control (no siRNA treatment) under **A-C,** 1 hour of HUVEC treatment with 50 ng.mL$^{-1}$ of VEGF$_{165a}$. **D-F,** 4 hours of HUVEC treatment with 50 ng.mL$^{-1}$ of VEGF$_{165a}$. **G-I,** 1 hour of HUVEC treatment with 50 ng.mL$^{-1}$ of PLGF $_{1}$. **J-L,** 4 hours of HUVEC treatment with 50 ng.mL$^{-1}$ of PLGF$_{1}$.

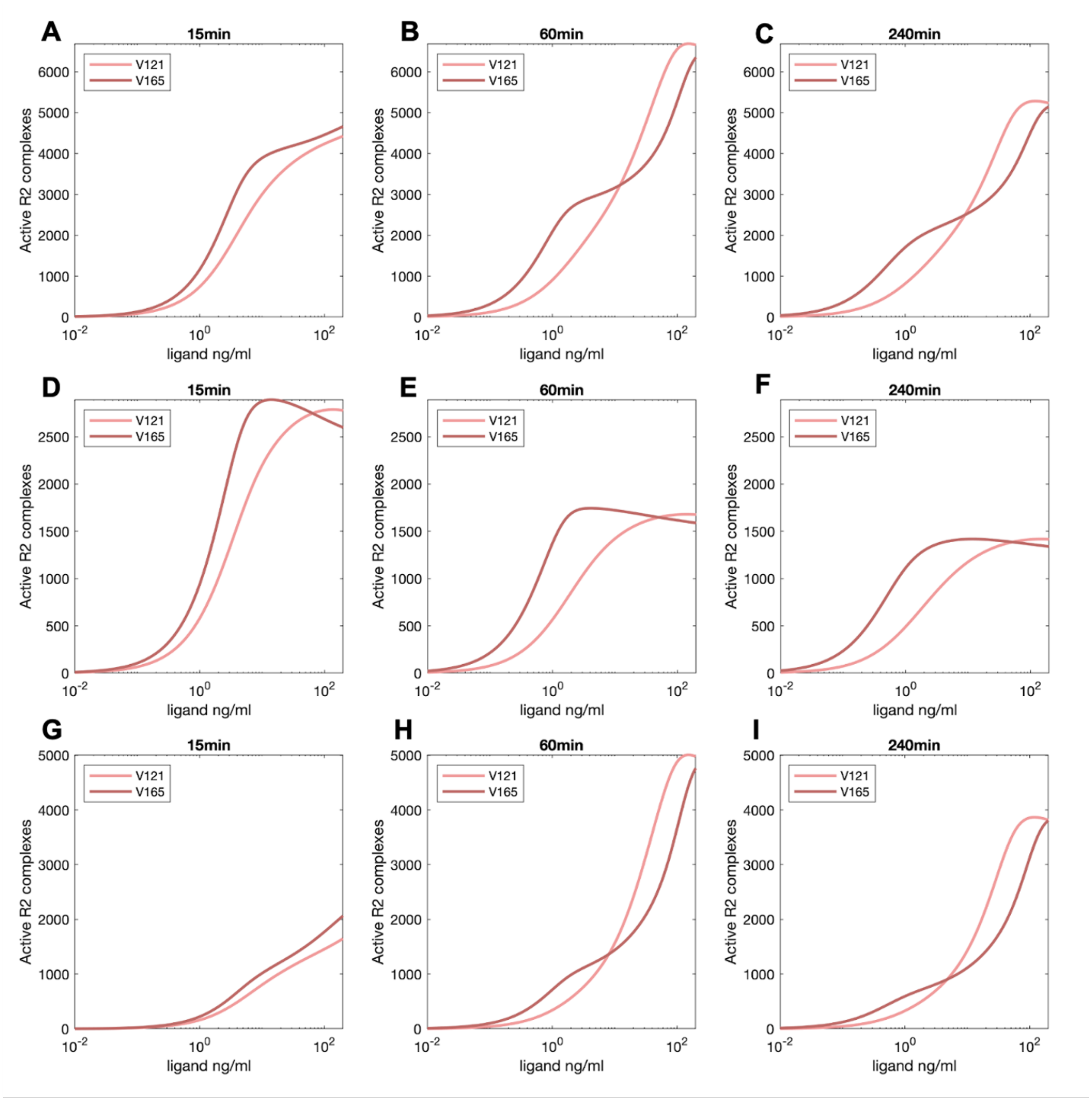

**Figure S9. Active VEGFR2 following VEGF treatment.** Previous supplemental figures S2-S7 have illustrated localization of all receptors, without regard to whether they are in complexes with other receptors or ligands. Here, we focus on active, ligated receptors. Number of active (ligand-dimerized) VEGFR2 receptors on the whole cell (A-C), cell surface (D-F), and internally (G-I) following 15 min, 60 min, or 240 min of treatment with $VEGF_{121a}$ or $VEGF_{165a}$.

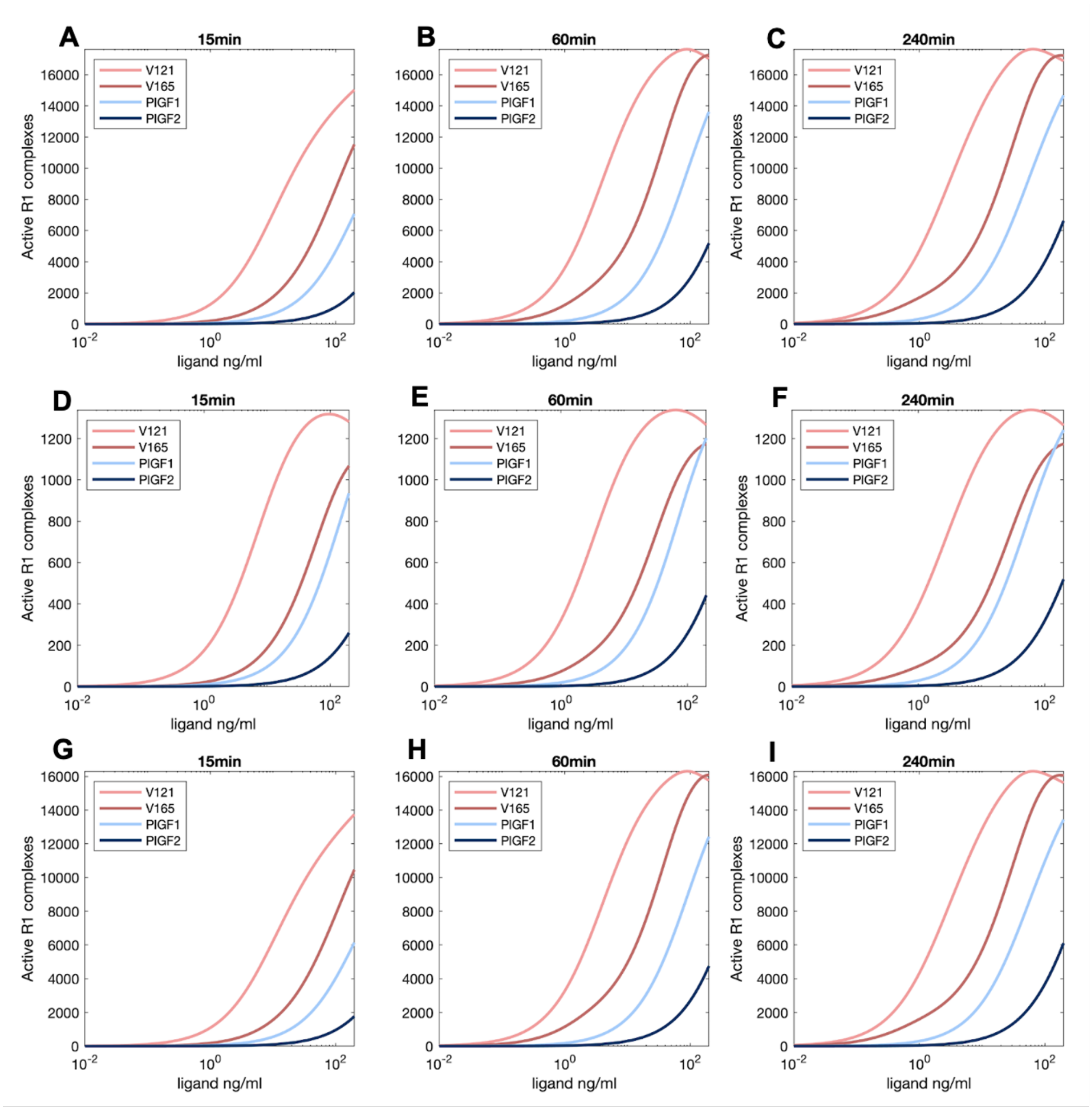

**Figure S10. Active VEGFR1 following VEGF or PLGF treatment.** Number of active (ligand-dimerized) VEGFR1 receptors on the whole cell (A-C), cell surface (D-F), and internally (G-I) following 15 min, 60 min, or 240 min of treatment with $VEGF_{121a}$, $VEGF_{165a}$, $PLGF_1$, or $PLGF_2$.

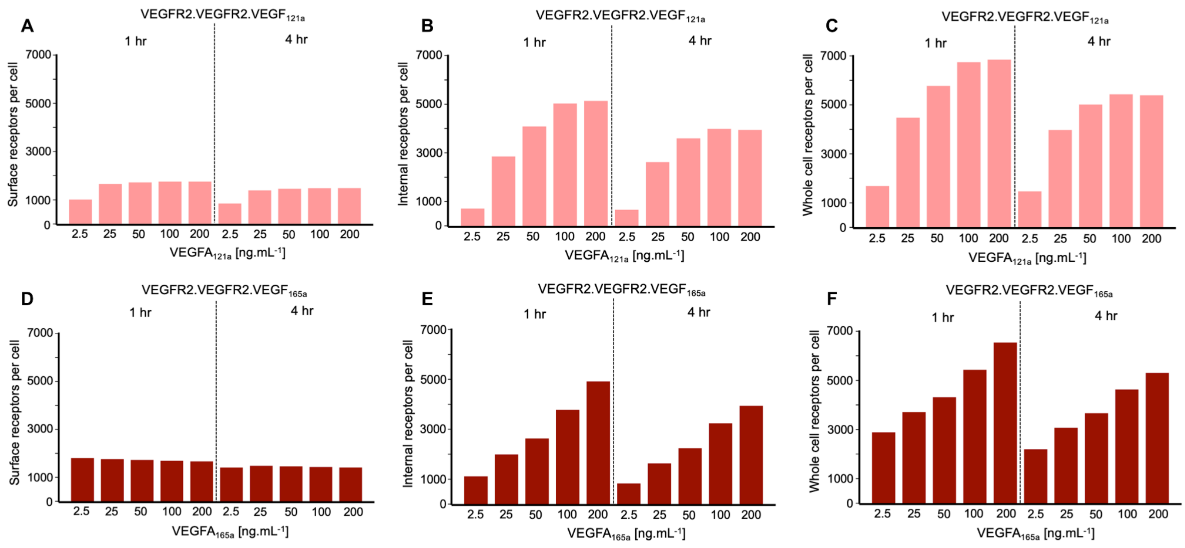

**Figure S11. Ligand dose-dependent distribution of ligated VEGFR2.** different single ligand doses 2.5, 25, 50, 100, 200 ng.mL$^{-1}$ of **A-C,** $VEGF_{121a}$ treatment or **D-F,** $VEGF_{165a}$ treatment.

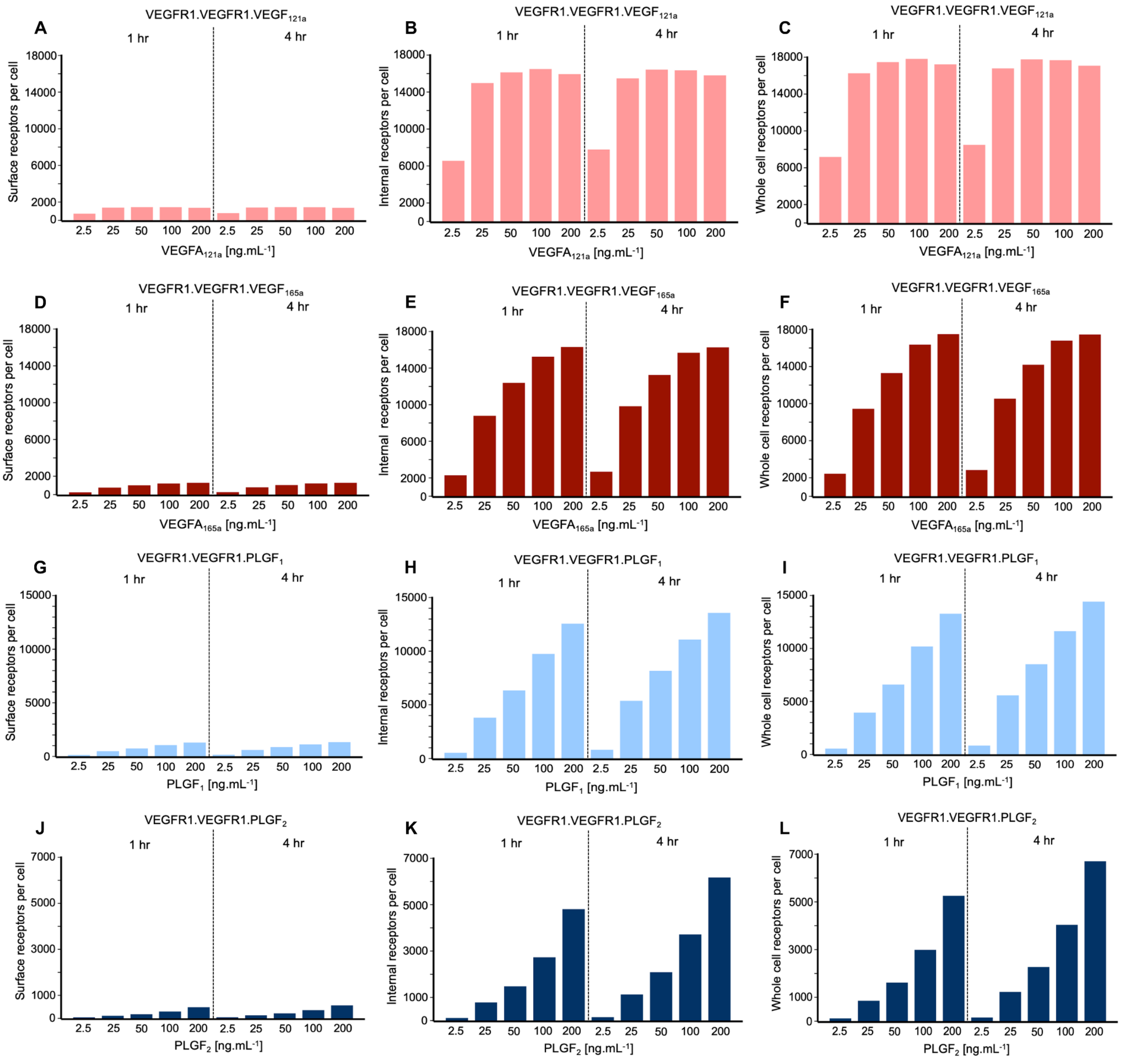

**Figure S12. Ligand dose-dependent distribution of ligated VEGFR1.** different single ligand doses 2.5, 25, 50, 100, 200 ng.mL$^{-1}$ of **A-C,** $VEGF_{121a}$, **D-F,** $VEGF_{165a}$, **G-I,** $PLGF_1$ or **J-L,** $PLGF_2$.

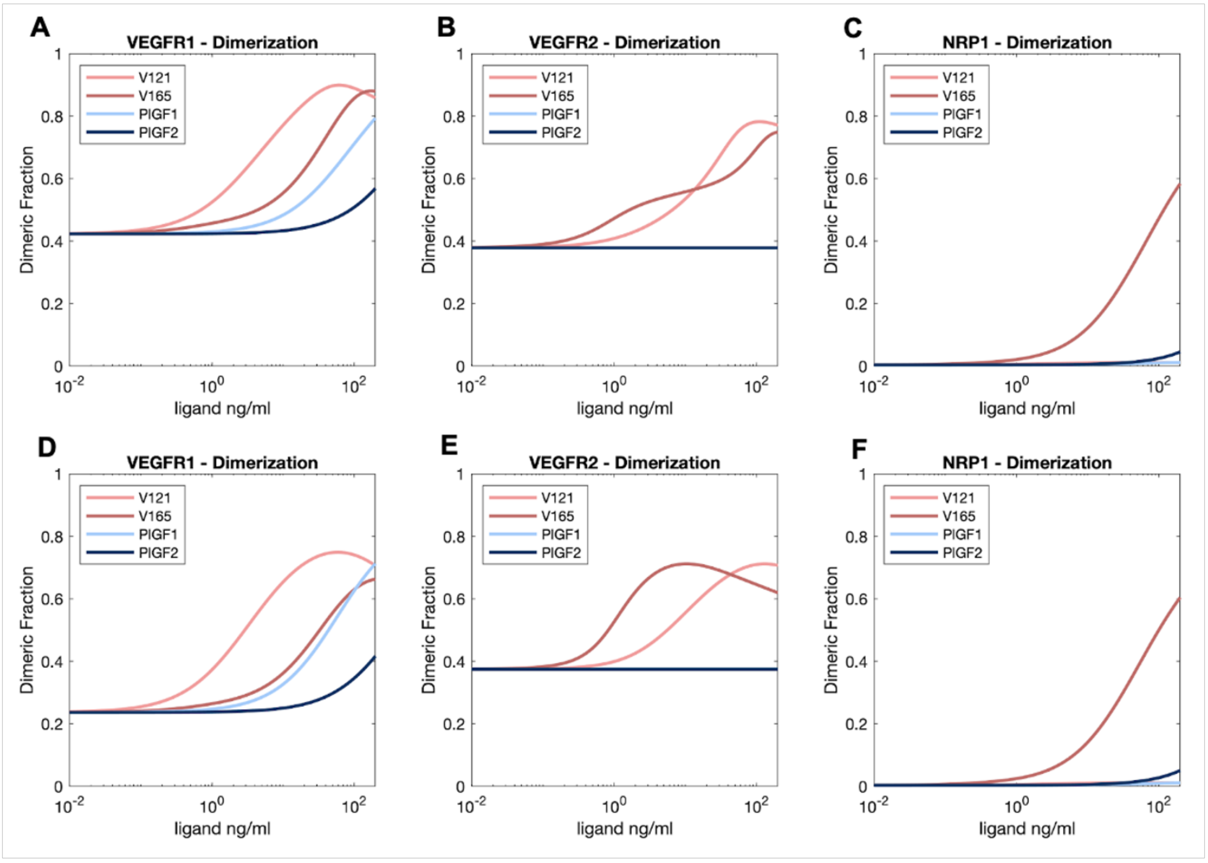

**Figure S13. Dimerization of VEGFR1, VEGFR2 and NRP1.** VEGF receptors can be dimerized in the absence of ligand; they can also be ligand-bound and dimerized without being active (if the ligand is not bound to both receptors). These graphs show the impact on receptor dimerization (not activation) across the whole cell (A-C), or on the cell surface (D-F) after 240 min of treatment with 50 ng.ml$^{-1}$ VEGF$_{121a}$, VEGF$_{165a}$, PLGF$_1$, or PLGF$_2$.

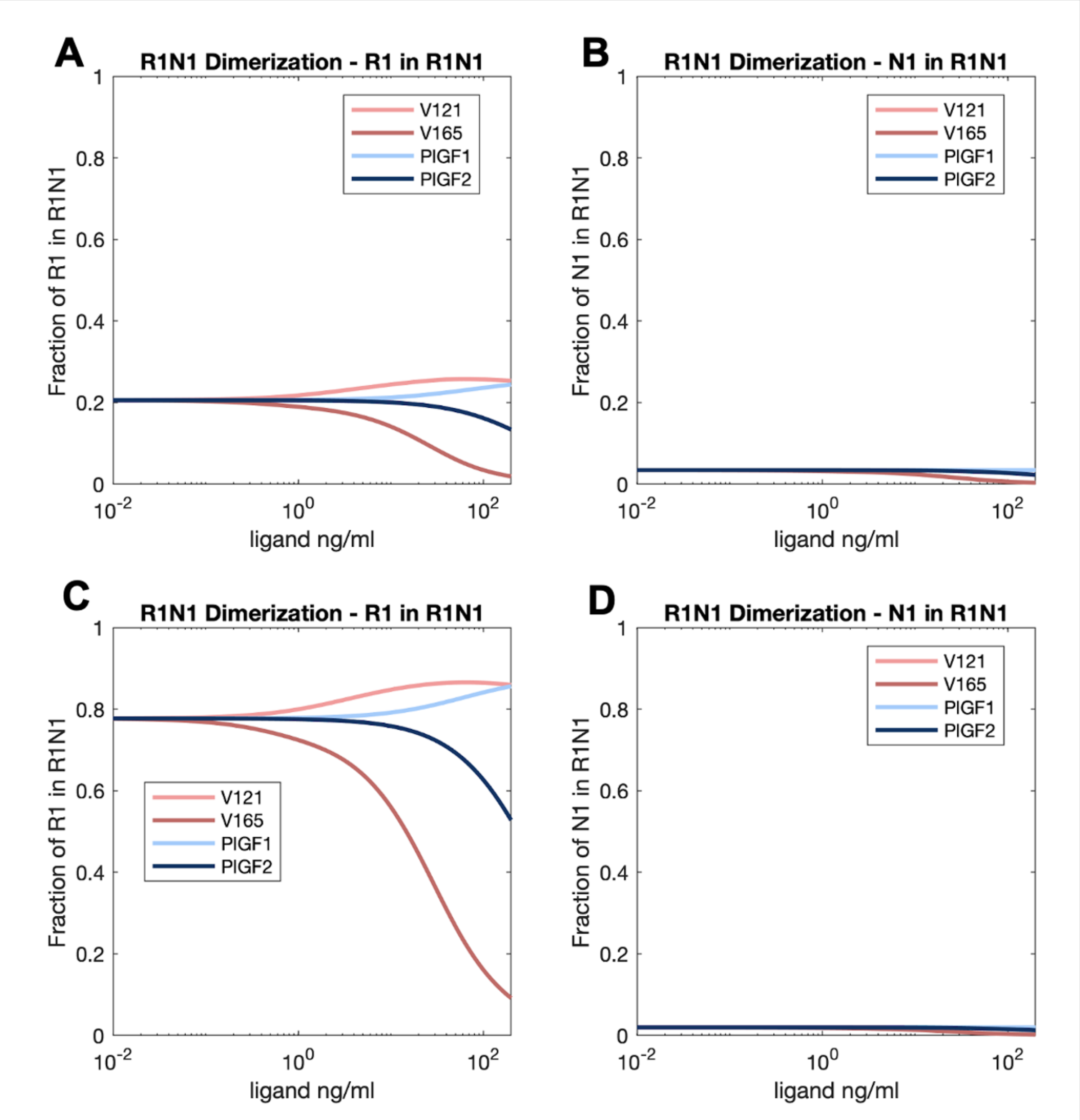

**Figure S14. Fraction of VEGFR1-NRP1 heterodimers.** VEGFR1 and NRP1 can associate without ligands (Fuh *et al*, 2000), and the VEGFR1-NRP1 complex formation affected by the presence of ligands that can bind to the complex ($VEGF_{121a}$ and $PLGF_1$) and those that cannot ($VEGF_{165a}$ and $PLGF_2$). These graphs show the impact of VEGFR1-NRP1 dimerization across the whole cell (A-B) or on the cell surface (C-D) after 240 minutes of 50 $ng.ml^{-1}$ $VEGF_{121a}$ , $VEGF_{165a}$ , $PLGF_1$ , or $PLGF_2$.

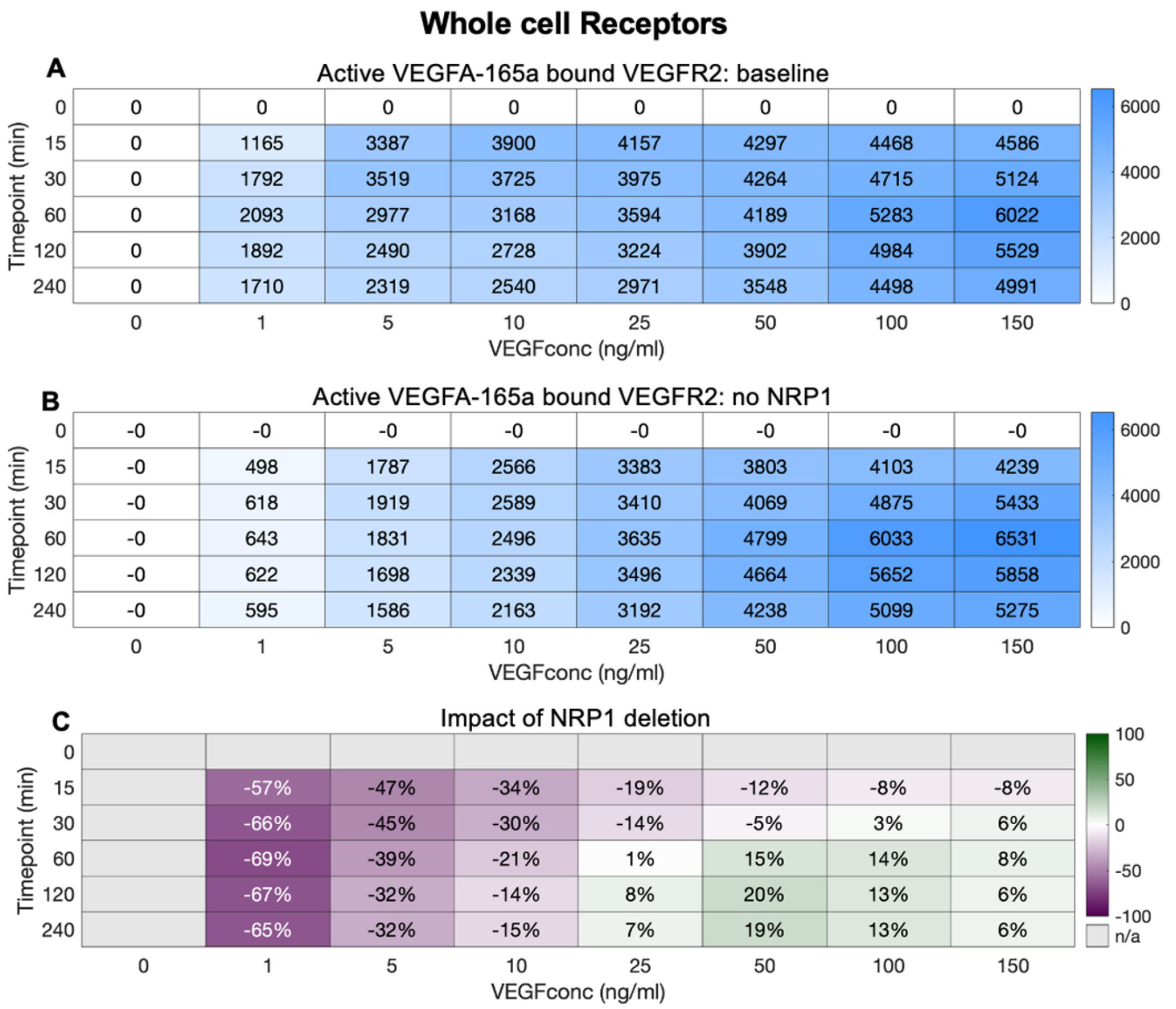

**Figure S15. Impact of NRP1 expression on whole cell VEGFR2 activation by VEGF$_{165a}$. A,** VEGFR2.VEGF$_{165a}$.VEGFR2 levels for different initial VEGF$_{165a}$ concentrations. **B,** VEGFR2.VEGF$_{165a}$.VEGFR2 levels in the absence of NRP1 for different initial VEGF$_{165a}$ concentrations. **C,** percent change in VEGFR2.VEGF$_{165a}$.VEGFR2 levels due to loss of NRP1 for different initial VEGF$_{165a}$ concentrations.

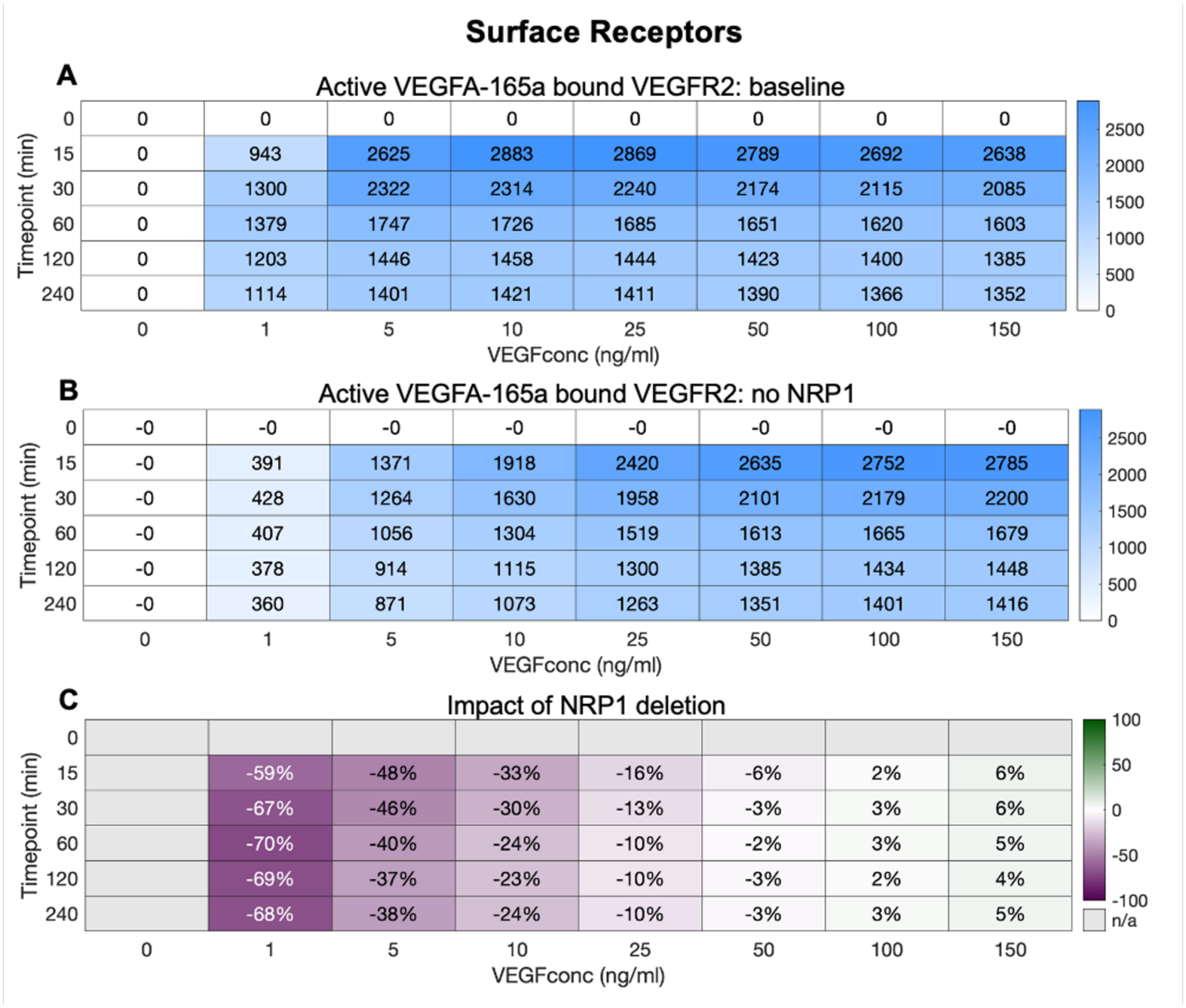

**Figure S16. Impact of NRP1 expression on surface VEGFR2 activation by $VEGF_{165a}$. A,** $VEGFR2.VEGF_{165a}.VEGFR2$ levels for different initial $VEGF_{165a}$ concentrations. **B,** $VEGFR2.VEGF_{165a}.VEGFR2$ levels in the absence of NRP1 for different initial $VEGF_{165a}$ concentrations. **C,** percent change in $VEGFR2.VEGF_{165a}.VEGFR2$ levels due to loss of NRP1 for different initial $VEGF_{165a}$ concentrations.

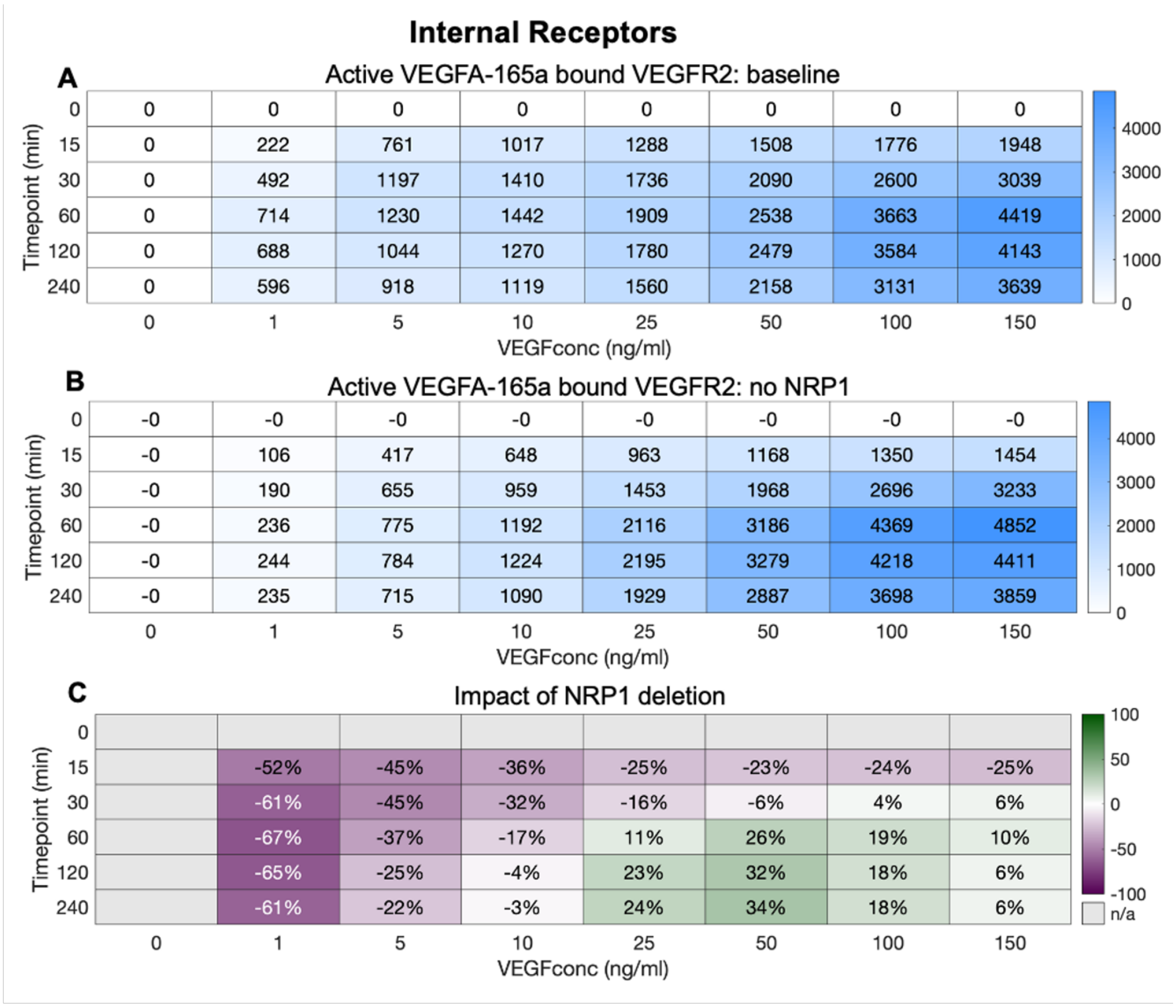

**Figure S17. Impact of NRP1 expression on internal VEGFR2 activation by VEGF$_{165a}$. A,** VEGFR2.VEGF$_{165a}$.VEGFR2 levels for different initial VEGF$_{165a}$ concentrations. **B,** VEGFR2.VEGF$_{165a}$.VEGFR2 levels in the absence of NRP1 for different initial VEGF$_{165a}$ concentrations. **C,** percent change in VEGFR2.VEGF$_{165a}$.VEGFR2 levels due to loss of NRP1 for different initial VEGF$_{165a}$ concentrations.

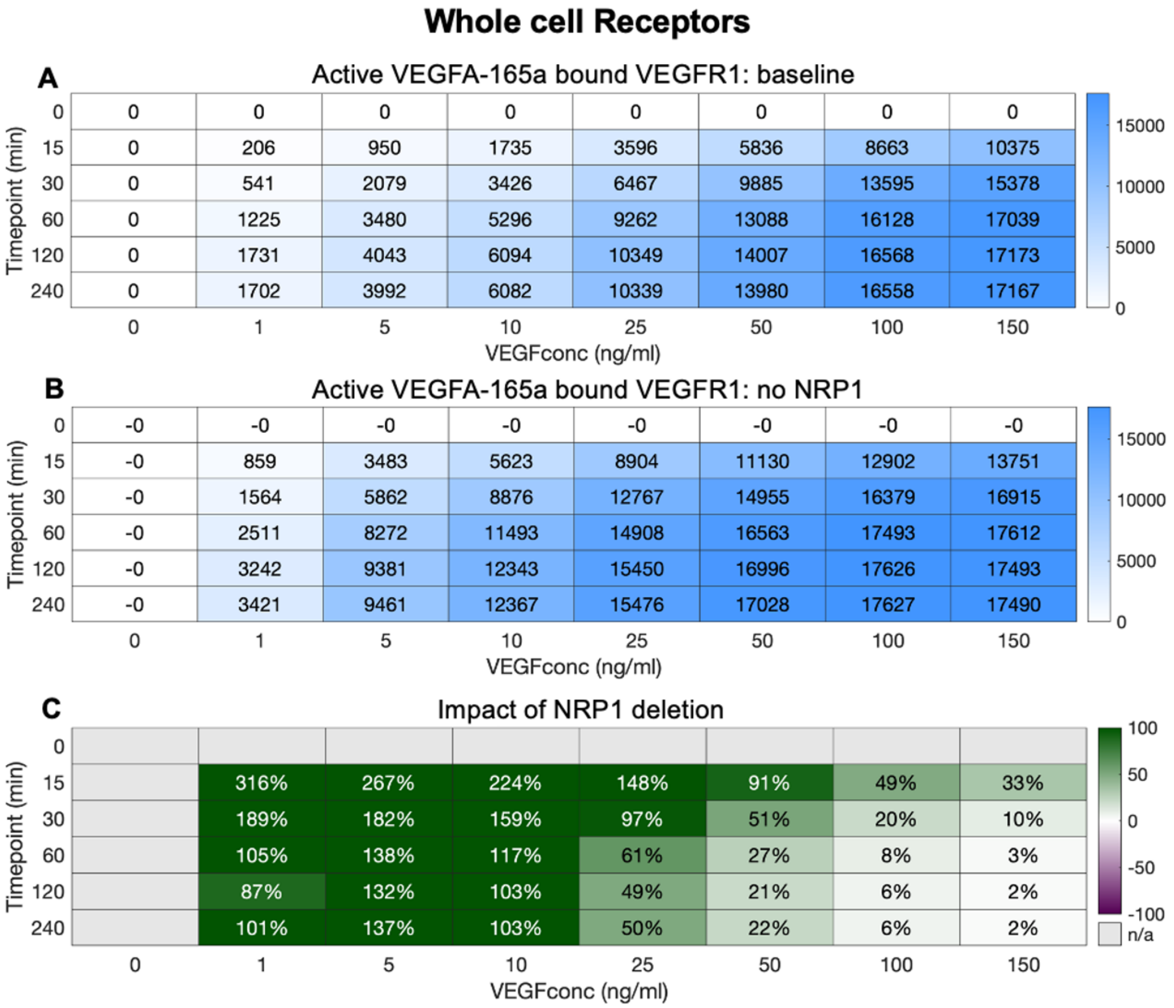

**Figure S18. Impact of NRP1 expression on whole cell VEGFR1 activation by $VEGF_{165a}$. A,** VEGFR1.$VEGF_{165a}$.VEGFR1 levels for different initial $VEGF_{165a}$ concentrations. **B,** VEGFR1.$VEGF_{165a}$.VEGFR1 levels in the absence of NRP1 for different initial $VEGF_{165a}$ concentrations. **C,** percent change in VEGFR1.$VEGF_{165a}$.VEGFR1 levels due to loss of NRP1 for different initial $VEGF_{165a}$ concentrations.

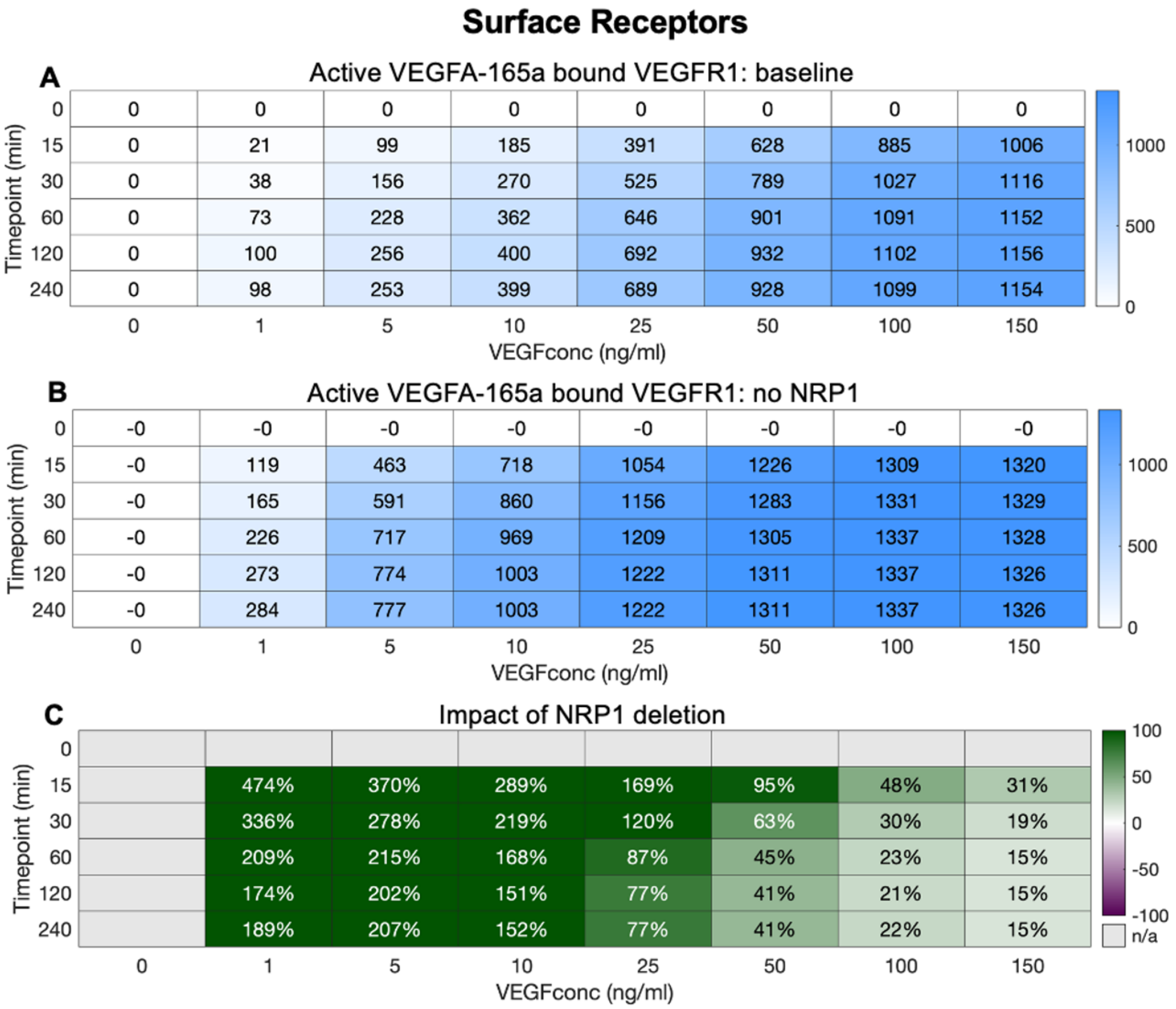

**Figure S19. Impact of NRP1 expression on surface VEGFR1 activation by VEGF$_{165a}$. A,** VEGFR1.VEGF$_{165a}$.VEGFR1 levels for different initial VEGF$_{165a}$ concentrations. **B,** VEGFR1.VEGF$_{165a}$.VEGFR1 levels in the absence of NRP1 for different initial VEGF$_{165a}$ concentrations **C,** percent change in VEGFR1.VEGF$_{165a}$.VEGFR1 levels due to loss of NRP1 for different initial VEGF$_{165a}$ concentrations.

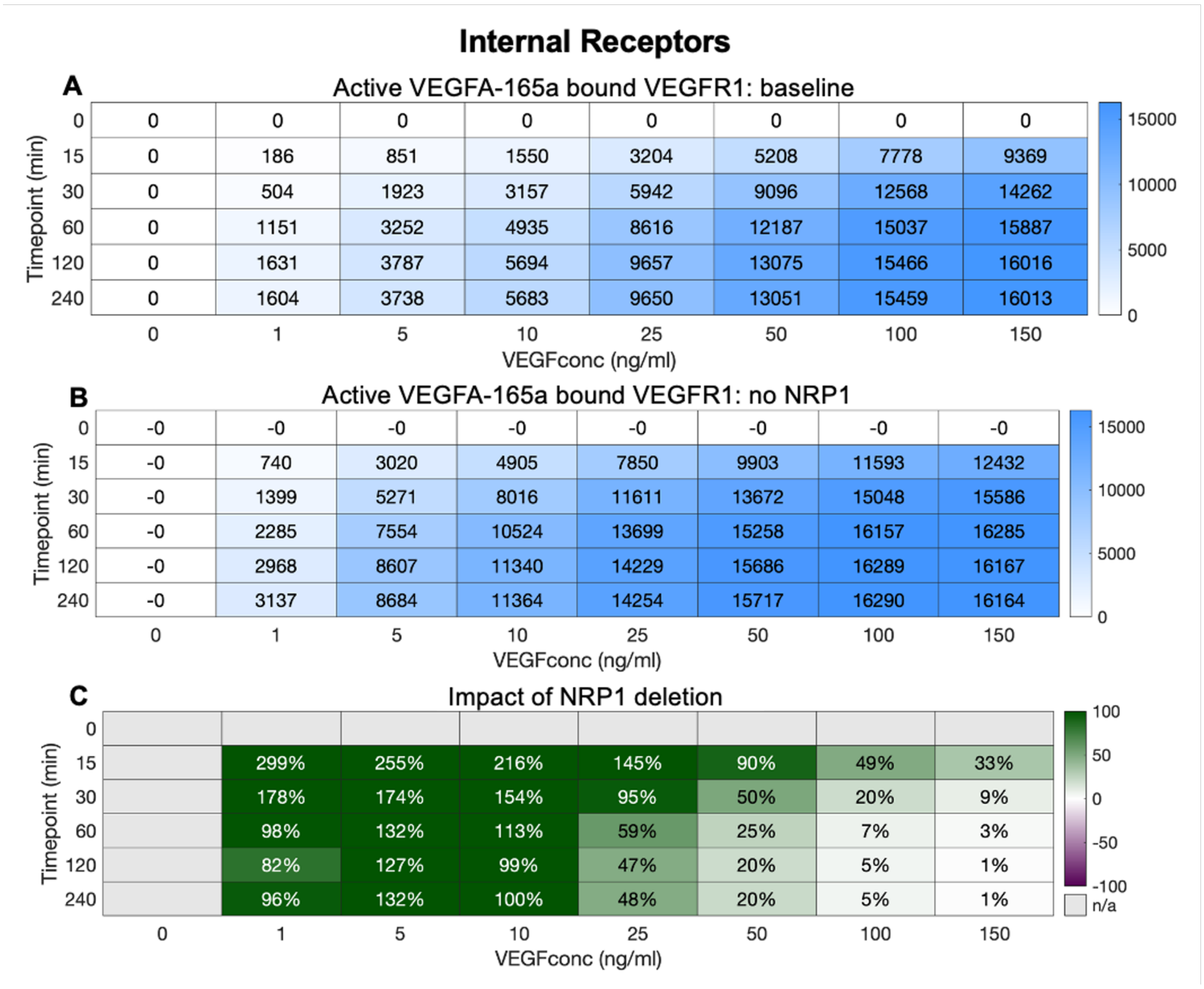

**Figure S20. Impact of NRP1 expression on internal VEGFR1 activation by VEGF$_{165a}$. A,** VEGFR1.VEGF$_{165a}$.VEGFR1 levels for different initial VEGF$_{165a}$ concentrations. **B,** VEGFR1.VEGF$_{165a}$.VEGFR1 levels in the absence of NRP1 for different initial VEGF$_{165a}$ concentrations. **C,** percent change in VEGFR1.VEGF$_{165a}$.VEGFR1 levels due to loss of NRP1 for different initial VEGF$_{165a}$ concentrations.

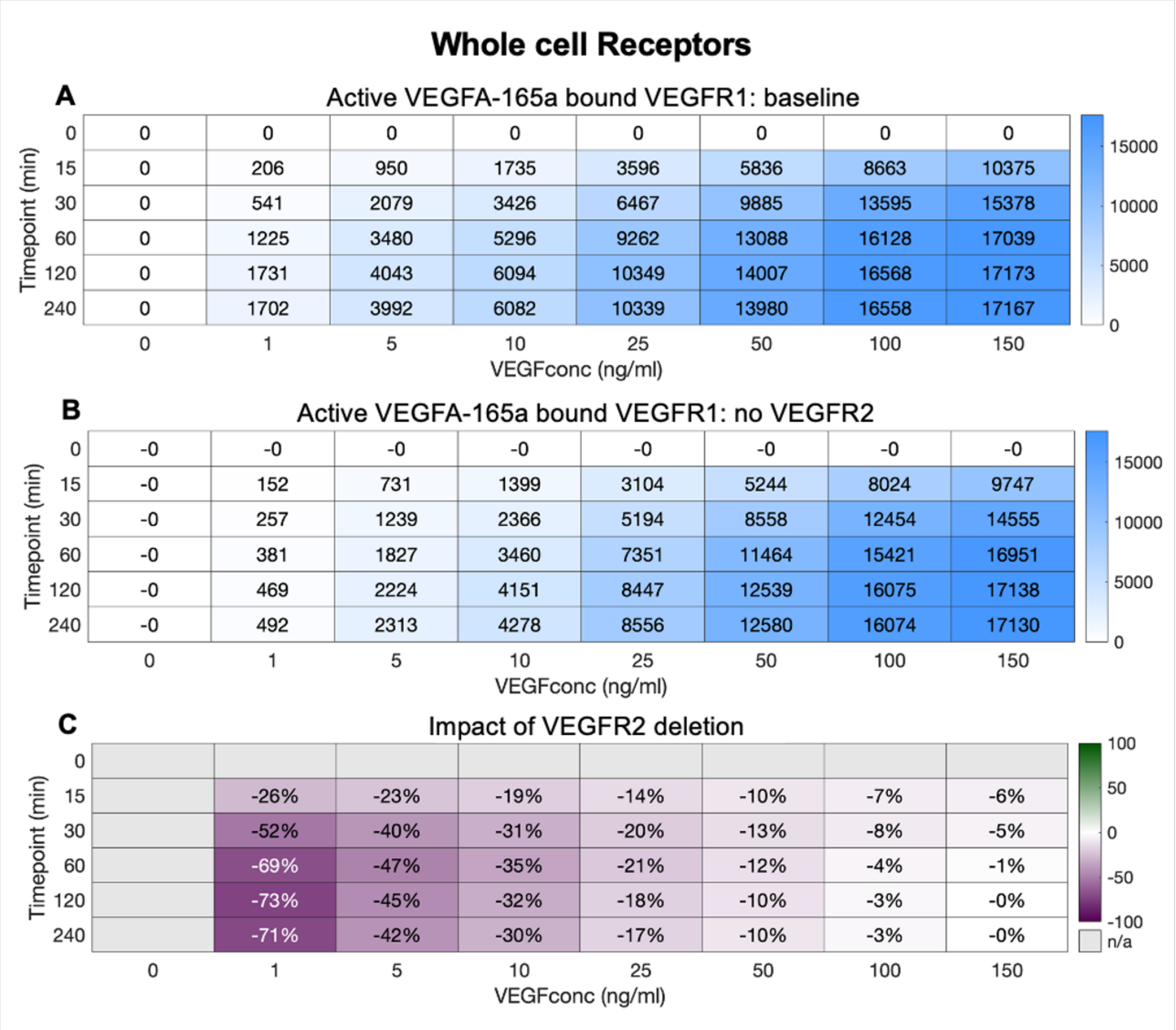

**Figure S21. Impact of VEGFR2 expression on whole cell VEGFR1 activation by $VEGF_{165a}$.** **A,** $VEGFR1.VEGF_{165a}.VEGFR1$ levels for different initial $VEGF_{165a}$ concentrations. **B,** $VEGFR1.VEGF_{165a}.VEGFR1$ levels in the absence of VEGFR2 for different initial $VEGF_{165a}$ concentrations. **C,** percent change in $VEGFR1.VEGF_{165a}.VEGFR1$ levels due to loss of VEGFR2 for different initial $VEGF_{165a}$ concentrations.

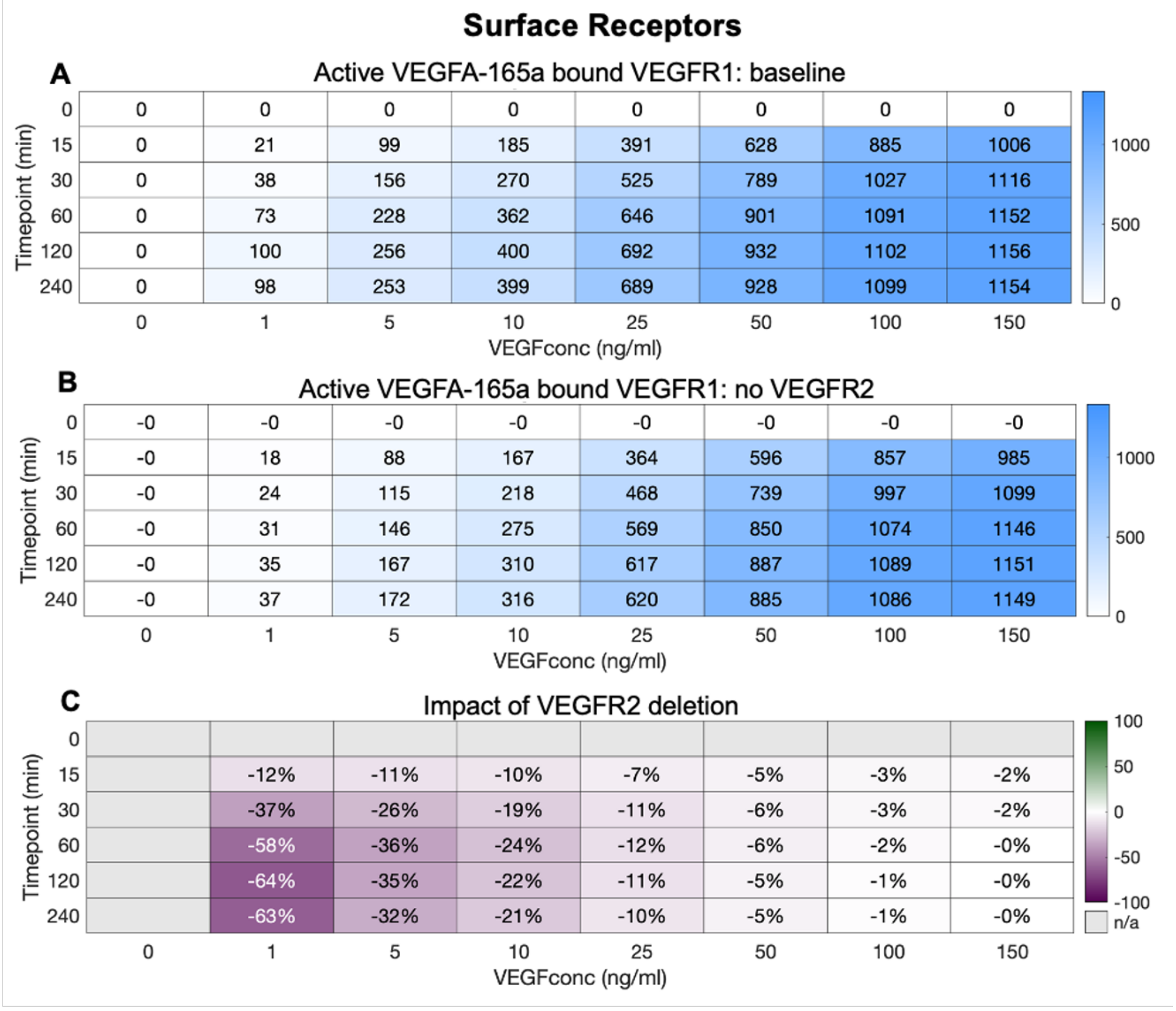

**Figure S22. Impact of VEGFR2 expression on surface VEGFR1 activation by VEGF$_{165a}$. A,** VEGFR1.VEGF$_{165a}$.VEGFR1 levels for different initial VEGF$_{165a}$ concentrations. **B,** VEGFR1.VEGF$_{165a}$.VEGFR1 levels in the absence of VEGFR2 for different initial VEGF$_{165a}$ concentrations. **C,** percent change in VEGFR1.VEGF$_{165a}$.VEGFR1 levels due to VEGFR2 for different initial VEGF$_{165a}$ concentrations.

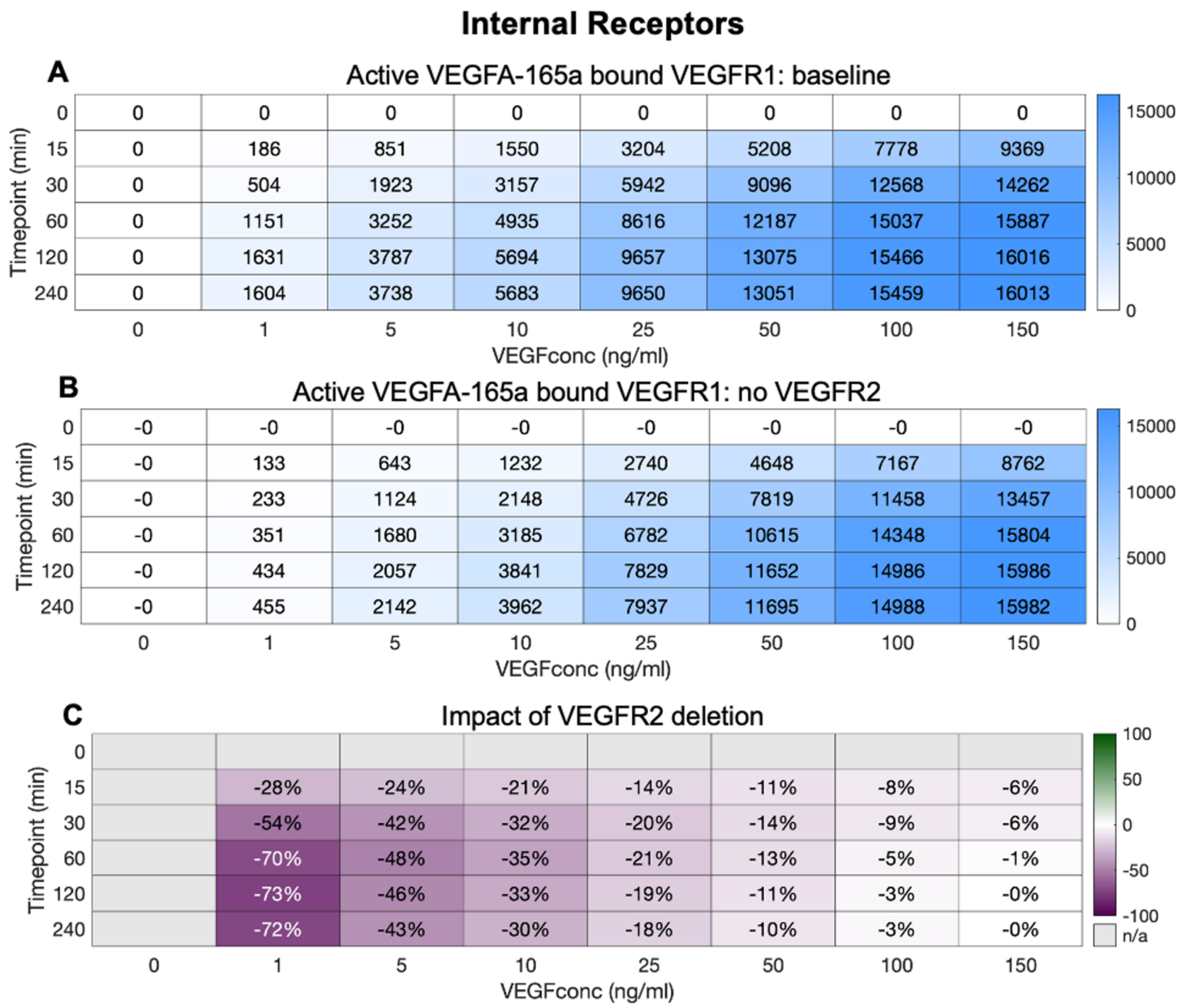

**Figure S23. Impact of VEGFR2 expression on internal VEGFR1 activation by $VEGF_{165a}$. A,** $VEGFR1.VEGF_{165a}.VEGFR1$ levels for different initial $VEGF_{165a}$ concentrations. **B,** $VEGFR1.VEGF_{165a}.VEGFR1$ levels in the absence of VEGFR2 for different initial $VEGF_{165a}$ concentrations. **C,** percent change in $VEGFR1.VEGF_{165a}.VEGFR1$ levels due to VEGFR2 for different initial $VEGF_{165a}$ concentrations.

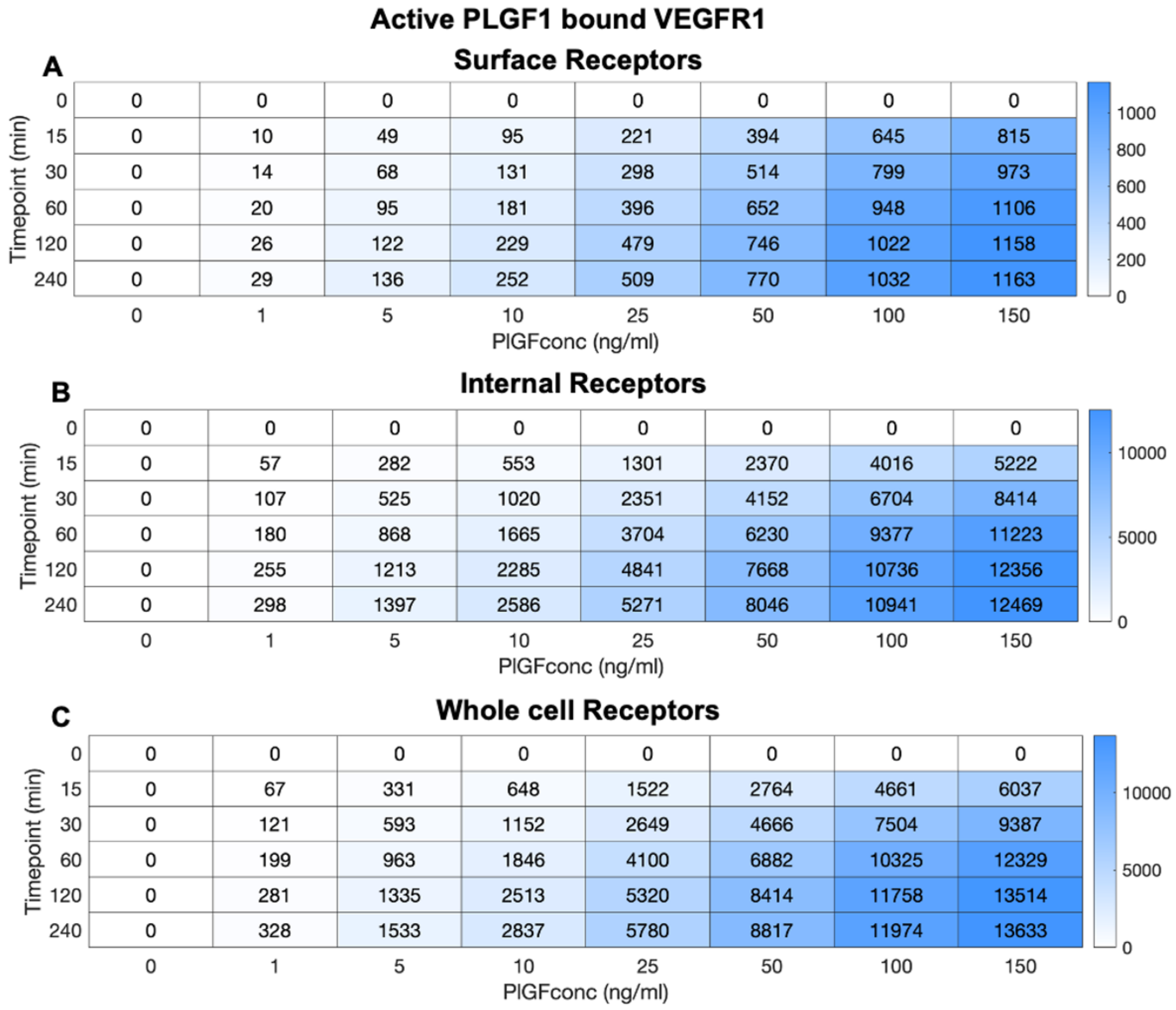

**Figure S24. Induction of active VEGFR1 by PLGF$_1$. A,** Surface, **B,** Internal, and **C,** Whole cell levels of VEGFR1.PLGF1.VEGFR1 complexes at different timepoints over 4 hours and under varying PLGF1 concentration.

**Active VEGFA-165a bound VEGFR1**

**Whole cell receptors**

| PlGFconc (ng/ml) \ VEGFconc (ng/ml) | 0 | 1 | 5 | 10 | 25 | 50 | 100 | 150 |
|---|---|---|---|---|---|---|---|---|
| 0 | 0 | 1731 | 4043 | 6094 | 10349 | 14007 | 16568 | 17173 |
| 1 | 0 | 1721 | 4012 | 6045 | 10278 | 13935 | 16516 | 17136 |
| 5 | 0 | 1684 | 3892 | 5858 | 10003 | 13657 | 16309 | 16991 |
| 10 | 0 | 1640 | 3755 | 5642 | 9681 | 13325 | 16058 | 16814 |
| 25 | 0 | 1530 | 3409 | 5094 | 8837 | 12425 | 15353 | 16307 |
| 50 | 0 | 1393 | 2988 | 4420 | 7742 | 11180 | 14317 | 15539 |
| 100 | 0 | 1211 | 2462 | 3567 | 6257 | 9340 | 12643 | 14236 |
| 150 | 0 | 1089 | 2136 | 3039 | 5289 | 8040 | 11350 | 13172 |

Colour scale: 0, 5000, 10000, 15000

**Surface receptors**

| PlGFconc (ng/ml) \ VEGFconc (ng/ml) | 0 | 1 | 5 | 10 | 25 | 50 | 100 | 150 |
|---|---|---|---|---|---|---|---|---|
| 0 | 0 | 100 | 256 | 400 | 692 | 932 | 1102 | 1156 |
| 1 | 0 | 99 | 254 | 396 | 686 | 926 | 1099 | 1154 |
| 5 | 0 | 96 | 243 | 380 | 663 | 904 | 1083 | 1143 |
| 10 | 0 | 92 | 231 | 361 | 637 | 878 | 1064 | 1129 |
| 25 | 0 | 82 | 202 | 316 | 569 | 809 | 1012 | 1092 |
| 50 | 0 | 70 | 167 | 261 | 482 | 714 | 936 | 1035 |
| 100 | 0 | 56 | 127 | 196 | 370 | 578 | 815 | 939 |
| 150 | 0 | 48 | 104 | 158 | 300 | 485 | 722 | 861 |

Colour scale: 0, 500, 1000

**Internal receptors**

| PlGFconc (ng/ml) \ VEGFconc (ng/ml) | 0 | 1 | 5 | 10 | 25 | 50 | 100 | 150 |
|---|---|---|---|---|---|---|---|---|
| 0 | 0 | 1631 | 3787 | 5694 | 9657 | 13075 | 15466 | 16016 |
| 1 | 0 | 1622 | 3758 | 5649 | 9592 | 13009 | 15417 | 15982 |
| 5 | 0 | 1588 | 3649 | 5478 | 9340 | 12753 | 15226 | 15849 |
| 10 | 0 | 1549 | 3523 | 5281 | 9044 | 12447 | 14994 | 15685 |
| 25 | 0 | 1448 | 3207 | 4779 | 8269 | 11616 | 14341 | 15216 |
| 50 | 0 | 1322 | 2821 | 4159 | 7260 | 10466 | 13381 | 14504 |
| 100 | 0 | 1155 | 2335 | 3371 | 5887 | 8761 | 11828 | 13298 |
| 150 | 0 | 1042 | 2032 | 2882 | 4989 | 7555 | 10628 | 12311 |

Colour scale: 0, 5000, 10000, 15000

**Fig S25. PLGF-VEGF competition: impact on active VEGF-bound VEGFR1 complexes.** Predicted level of active VEGFR1.VEGF.VEGFR1 complexes, following two hours of ligand treatment at different doses for $PLGF_1$ and $VEGF_{165}$, across the whole cell (top), on the cell surface (middle), and intracellularly (bottom).

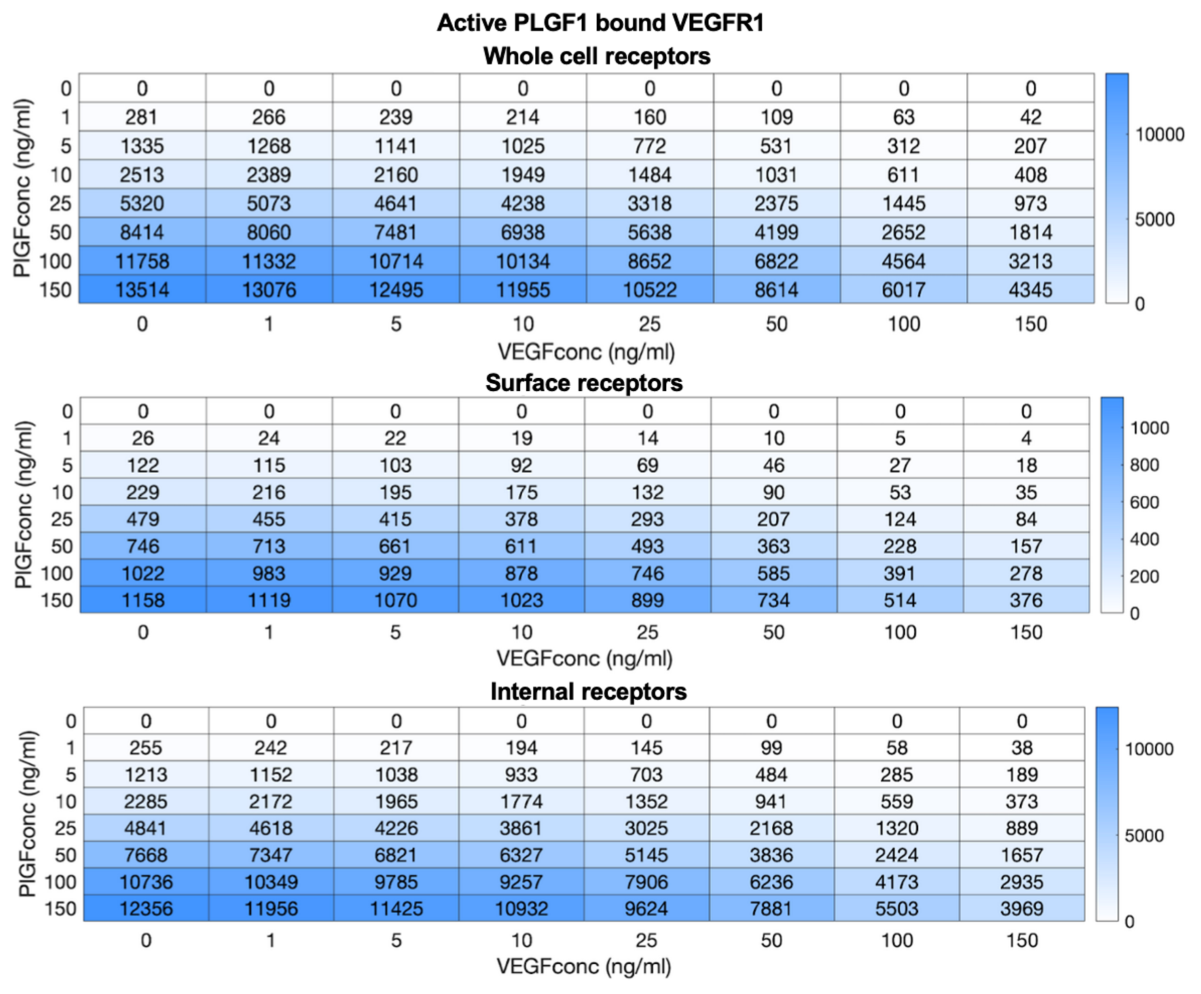

**Fig S26. PLGF-VEGF competition: impact on active PLGF-bound VEGFR1 complexes.** Predicted level of active VEGFR1.PLGF.VEGFR1 complexes, following two hours of ligand treatment at different doses for $PLGF_1$ and $VEGF_{165}$, across the whole cell (top), on the cell surface (middle), and intracellularly (bottom).

### VEGF-165a concentration

#### Cell surface

| PlGFconc (ng/ml) \ VEGFconc (ng/ml) | 0 | 1 | 5 | 10 | 25 | 50 | 100 | 150 |
|---|---|---|---|---|---|---|---|---|
| 0 | 0 | 22 | 112 | 225 | 564 | 1130 | 2264 | 3399 |
| 1 | 0 | 22 | 112 | 225 | 564 | 1130 | 2264 | 3399 |
| 5 | 0 | 22 | 112 | 225 | 564 | 1130 | 2264 | 3400 |
| 10 | 0 | 22 | 112 | 225 | 564 | 1130 | 2264 | 3400 |
| 25 | 0 | 22 | 112 | 225 | 564 | 1130 | 2265 | 3400 |
| 50 | 0 | 22 | 112 | 225 | 564 | 1131 | 2265 | 3400 |
| 100 | 0 | 22 | 112 | 225 | 565 | 1131 | 2265 | 3400 |
| 150 | 0 | 22 | 112 | 225 | 565 | 1131 | 2265 | 3400 |

#### Internal

| PlGFconc (ng/ml) \ VEGFconc (ng/ml) | 0 | 1 | 5 | 10 | 25 | 50 | 100 | 150 |
|---|---|---|---|---|---|---|---|---|
| 0 | 0 | 5 | 16 | 30 | 80 | 195 | 615 | 1458 |
| 1 | 0 | 5 | 16 | 30 | 80 | 195 | 615 | 1457 |
| 5 | 0 | 5 | 16 | 30 | 81 | 195 | 617 | 1455 |
| 10 | 0 | 5 | 16 | 31 | 81 | 196 | 618 | 1453 |
| 25 | 0 | 5 | 17 | 32 | 82 | 198 | 624 | 1449 |
| 50 | 0 | 6 | 19 | 34 | 85 | 204 | 634 | 1445 |
| 100 | 0 | 8 | 22 | 38 | 93 | 217 | 657 | 1439 |
| 150 | 0 | 10 | 25 | 43 | 101 | 232 | 679 | 1433 |

### PLGF1 concentration

#### Cell surface

| PlGFconc (ng/ml) \ VEGFconc (ng/ml) | 0 | 1 | 5 | 10 | 25 | 50 | 100 | 150 |
|---|---|---|---|---|---|---|---|---|
| 0 | 0 | 0 | 0 | 0 | 0 | 0 | 0 | 0 |
| 1 | 34 | 34 | 34 | 34 | 34 | 34 | 34 | 34 |
| 5 | 168 | 168 | 168 | 168 | 168 | 168 | 168 | 168 |
| 10 | 336 | 336 | 336 | 336 | 337 | 337 | 337 | 337 |
| 25 | 841 | 841 | 841 | 841 | 841 | 841 | 842 | 842 |
| 50 | 1682 | 1682 | 1683 | 1683 | 1683 | 1683 | 1683 | 1683 |
| 100 | 3366 | 3366 | 3366 | 3366 | 3366 | 3366 | 3367 | 3367 |
| 150 | 5049 | 5049 | 5049 | 5049 | 5049 | 5050 | 5050 | 5050 |

#### Internal

| PlGFconc (ng/ml) \ VEGFconc (ng/ml) | 0 | 1 | 5 | 10 | 25 | 50 | 100 | 150 |
|---|---|---|---|---|---|---|---|---|
| 0 | 0 | 0 | 0 | 0 | 0 | 0 | 0 | 0 |
| 1 | 4 | 5 | 5 | 5 | 6 | 7 | 11 | 14 |
| 5 | 22 | 23 | 24 | 26 | 30 | 37 | 53 | 72 |
| 10 | 44 | 47 | 49 | 52 | 60 | 74 | 107 | 144 |
| 25 | 117 | 124 | 131 | 137 | 155 | 190 | 273 | 365 |
| 50 | 248 | 266 | 279 | 291 | 328 | 395 | 560 | 737 |
| 100 | 529 | 575 | 606 | 632 | 709 | 844 | 1160 | 1472 |
| 150 | 819 | 903 | 959 | 1004 | 1129 | 1334 | 1775 | 2178 |

**Fig S27. PLGF-VEGF competition: ligand levels.** Predicted level of free (unbound) $VEGF_{165}$ (top) or $PLGF_1$ (bottom), at the cell surface or intracellularly, following two hours of ligand treatment at different doses for $PLGF_1$ and $VEGF_{165}$. Ligand levels are given in units of #/cell.

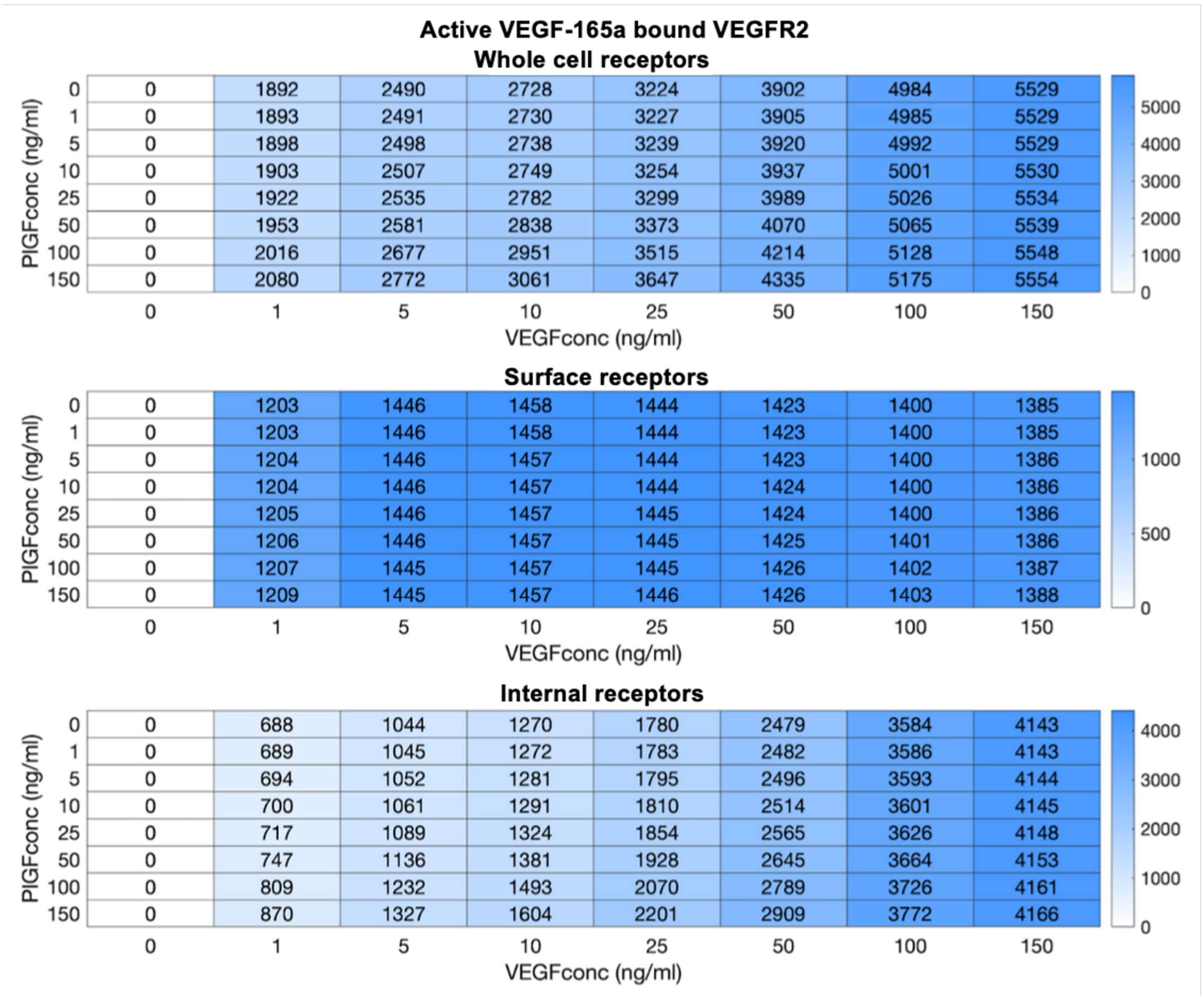

**Fig S28. PLGF-VEGF competition: impact on active VEGF-bound VEGFR2 complexes.** Predicted level of active VEGFR2.VEGF.VEGFR2 complexes, following two hours of ligand treatment at different doses for $PLGF_1$ and $VEGF_{165}$, across the whole cell (top), on the cell surface (middle), and intracellularly (bottom).

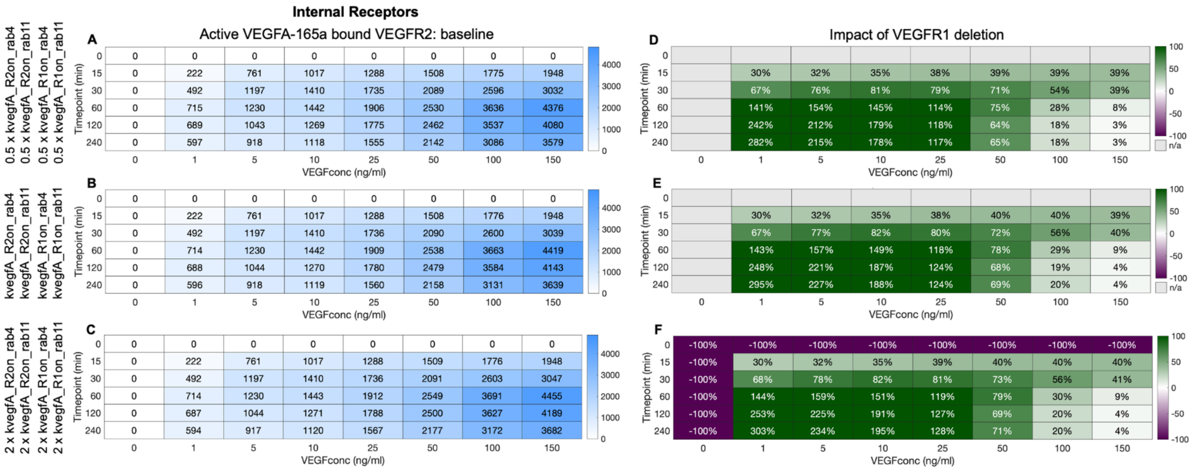

**Figure S29. Effect of change in endosomal pH on VEGFR2 activation and decoy effect.** Simulations with slower (top) or faster (bottom) rate constants in the endosomes than on the cell surface, due to pH differences. Effect of 4 hours of 50 ng.mL$^{-1}$ VEGF$_{165a}$ treatment on the intracellular levels of VEGFR2.VEGF$_{165a}$.VEGFR2 in HUVECs.

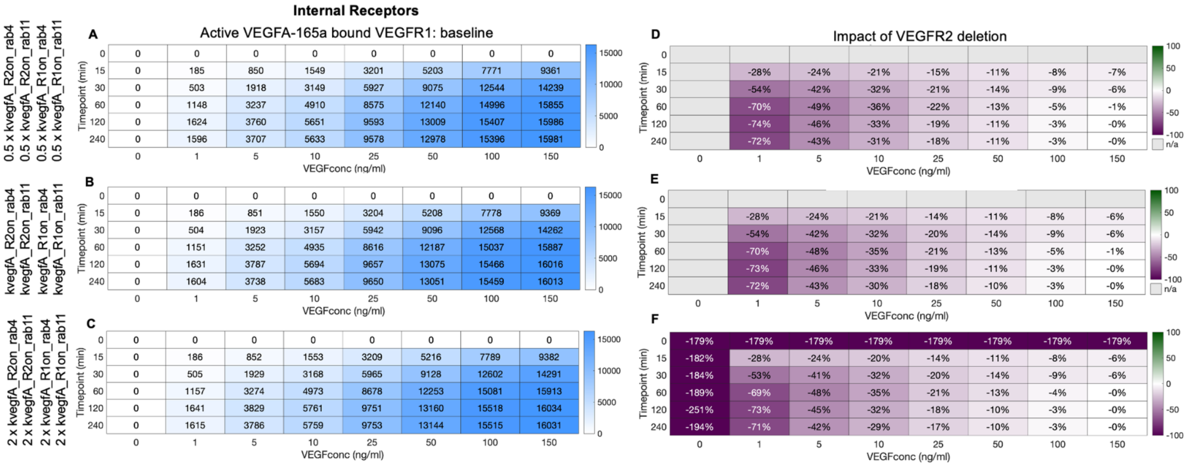

**Figure S30. Effect of change in endosomal pH on VEGFR1 activation and decoy effect.** Simulations with slower (top) or faster (bottom) rate constants in the endosomes than on the cell surface, due to pH differences. Effect of 4 hours of 50 ng.mL$^{-1}$ VEGF$_{165a}$ treatment on the intracellular levels of VEGFR1.VEGF$_{165a}$.VEGFR1 in HUVECs.
